# Transcontinental Spread of HPAI H5N1 from South America to Antarctica via Avian Vectors

**DOI:** 10.1101/2025.09.06.674605

**Authors:** Xu Ruifeng, Gao Minhao, Zhang Nailou, Wei Zhenhua, Wang Zheng, Zhang Lei, Liu Yang, Zheng Zhenhua, Chen Liulin, Ding Haitao, Wang Wei

**Affiliations:** Wuhan Institute of Virology, Chinese Academy of Sciences, Wuhan City, Hubei Province, 430071, China; Polar Research Institute of China, Shanghai, 200136, China

**Author notes:** Corresponding author: No. 262, Jinlong Avenue, Zhengdian Subdistrict, Jiangxia District, Wuhan City, Hubei Province, China, 027-87998197. These authors contributed equally to this work (Xu Ruifeng, Gao Minhao).

## Abstract

Our study has for the first time identified H5N1 strains (clade 2.3.4.4b, genotype B3.2) in brown skuas from the Fildes Peninsula, South Shetland Islands, Antarctica. These findings indicate that highly pathogenic avian influenza H5N1 viruses is now actively circulating in Antarctic ecosystems, representing a significant expansion of its geographic range.

## Dear editor

Highly Pathogenic Avian Influenza (HPAI) H5N1 viruses originating from the A/Goose/Guangdong/1/96 (Gs/GD) lineage have demonstrated rapid genomic evolution and remarkable cross-species transmission capabilities. The clade 2.3.4.4b outbreaks have intensified significantly since 2020, spreading rapidly across Asia, Europe, Africa, North America, and South America, with substantial impacts on poultry and wildlife (Peacock et al., 2024). Notably, this clade has demonstrated unprecedented adaptive evolution in mammalian hosts, with confirmed infections in dairy cattle and domestic cats on farms (Caserta et al., 2024), as well as mass mortality events in southern elephant seals (Uhart et al., 2024). Human infections have also been documented (Garg et al., 2025), though sustained human-to-human transmission has not been observed.

The Antarctic region encompasses the Antarctic ice shelves, surrounding waters, and all island territories south of the Antarctic Convergence (Antarctic Polar Front), a marine boundary where cold Antarctic waters meet the warmer sub-Antarctic waters(Moore et al., 1999). This region supports unique ecosystems that serve as critical habitats for numerous avian and marine mammal species. Despite their geographical isolation, wildlife, such as brown skuas (*Stercorarius antarcticus*), southern giant petrels (*Macronectes giganteus*),Southern Elephant Seal(*Mirounga leonina*), Antarctic Fur Seal(*Arctocephalus gazella*), regularly breed in Antarctica but partially migrate to the coasts of Chile and Argentina during winter. Growing evidence suggests that viruses from South America may be spreading southward to Antarctica. In 2023, HPAI H5N1 was detected in sub-Antarctic regions including South Georgia (54°15′S, 36°45′W) and the Falkland Islands (51°42′S, 57°51′W)(Banyard et al., 2024). By 2024, the virus had reached James Ross Island (64°10′S, 57°45′W) in Antarctica, where it was identified in brown skuas, though no signs of HPAI were observed in wildlife on Fildes Peninsula(Bennett-Laso et al., 2024).

During China’s 41st Antarctic Scientific Expedition, we collected biological samples in the vicinity of the Great Wall Station (62°12′59”S, 58°57′52”W) on Fildes Peninsula, located in western King George Island of the South Shetland Islands, Antarctica. The samples included pharyngeal and anal swabs from deceased Antarctic fur seals, brain tissue and fecal samples from dead brown skuas that exhibited clinical signs of disheveled plumage, lethargy and inability to fly prior to death, as well as fresh fecal samples from gentoo penguins (*Pygoscelis papua*), southern giant petrels, and southern elephant seals (**Appendix Fig. S1 and Appendix Table S1**).

Real-time RT-PCR screening initially detected H5 positivity in four brown skua samples and one elephant seal fecal sample, along with H7 positivity in one fur seal sample, though subsequent retesting failed to confirm positivity in the elephant seal and fur seal specimens. To investigate the potential origins, genetic characteristics, and evolution of these viruses, we performed next-generation sequencing (NGS) and phylogenetic analyses, successfully obtaining complete genome sequences for all four H5N1-positive skua samples. The H5N1 viruses identified on Fildes Peninsula, Antarctica were formally designated as A/brown skua/Fildes Peninsula/B1/2024(H5N1), A/brown skua/Fildes Peninsula/B2/2024(H5N1), A/brown skua/Fildes Peninsula/B3/2024(H5N1), and A/brown skua/Fildes Peninsula/F4/2024(H5N1), with their complete genome sequences deposited in the Global Initiative on Sharing All Influenza Data (GISAID) under accession numbers EPI_ISL_19847535, EPI_ISL_19847536, EPI_ISL_19847538, and EPI_ISL_19847539, respectively. The genome sequence between viruses possessed high homology with each other.

To further characterize the obtained viral strains, we retrieved 514 representative H5N1 sequences encompassing all eight gene segments from the GISAID database for genotyping (**Appendix Table S2**). Subtyping based on hemagglutinin (HA) and neuraminidase (NA) protein sequences confirmed these isolates as H5N1 subtype, belonging to clade 2.3.4.4b genotype B3.2 (**Appendix Fig. S2**). Additionally, we retrieved 1,143 representative H5N1 clade 2.3.4.4b sequences, from the National Center for Biotechnology Information (NCBI) for phylogenetic analysis **(Appendix Table S3)**. Phylogenetic analysis revealed that viral sequences from Fildes Peninsula clustered closely with H5N1 strains originating from South America, particularly demonstrating strong phylogenetic affinity with Peruvian and Chilean isolates across all genomic segments (**Fig.1**). These genetically similar viruses exhibit broad circulation among both avian and marine mammal populations, and South America as the predominant geographic reservoir for this viral lineage (**Appendix Fig. S3** and **Appendix Fig. S4**).

**Figure 1.**
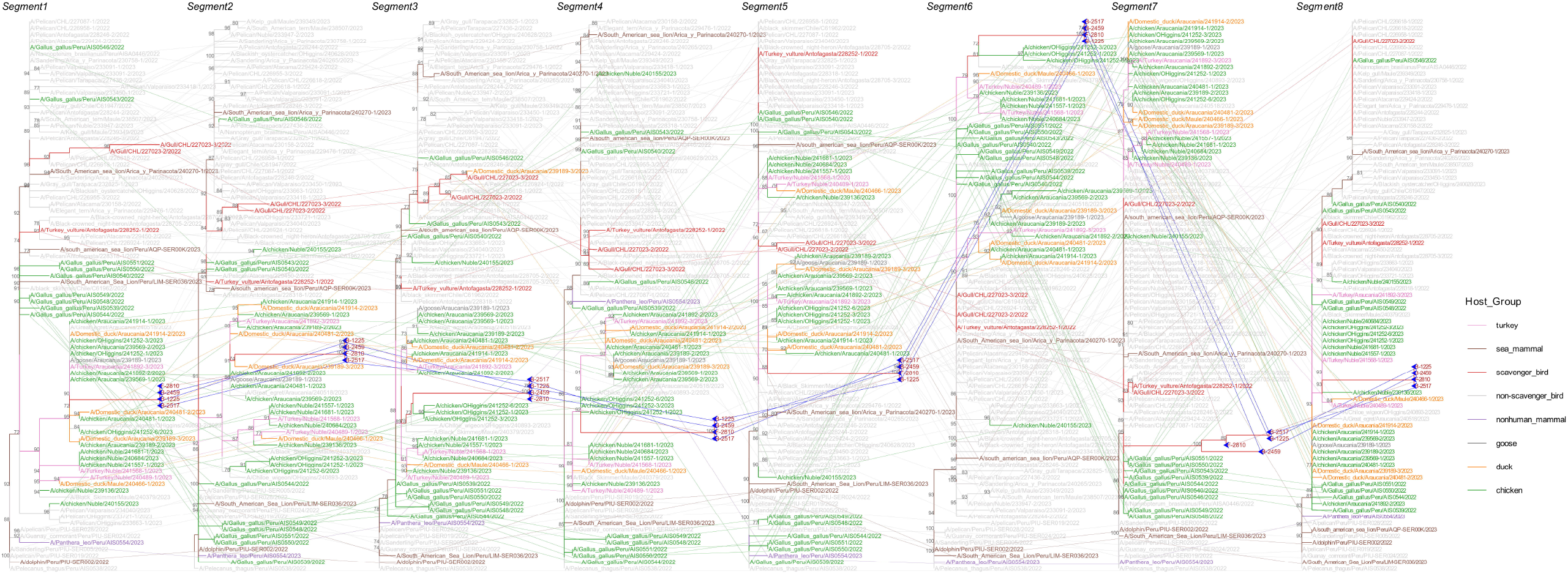
Phylogenetic incongruence analyses. Maximum likelihood trees for the all eight genomic segments (PB2, PB1, PA, HA, NP, NA, MP, and NS) from equivalent strains were connected across the trees. The phylogenetic branches are color-coded to identify host species: pink lines characterize turkey, brown lines depict sea mammal, red lines exemplify scavenger bird, gray lines typify non-scavenger bird, purple lines portray nonhuman mammal, dark gray lines illustrate goose, orange lines embody duck, and green lines personify chicken. Bright blue lines represent phylogenetic connections between the genomic segments of the H5N1 strain detected on Fildes Peninsula.

To investigate the potential introduction pathway of HPAI H5N1 to Fildes Peninsula, we performed discrete phylogeographic analysis using 90 closely related HA and NA sequences identified in our initial screening. The spatiotemporal reconstruction indicated Peru as the likely source country for the HPAI H5N1 virus detected on Fildes Peninsula, with transmission occurring through Chile before reaching Antarctica. Molecular dating traced the HA sequences to a Chilean strain sampled on December 3, 2022 (95% HPD: November 13-December 10, 2022), while the NA sequences originated from Chilean viruses circulating between November 25, 2022 (95% HPD: November 10, 2022-January 7, 2023). This spatiotemporal reconstruction reveals a transmission pathway whereby the virus spilled over from South American poultry populations to brown skuas, subsequently reaching Fildes Peninsula’s wildlife through the birds’ natural migratory routes. (**Appendix Fig. S5 A and B**).

Comparative analysis with the reference strain A/Goose/Guangdong/1/96 identified the NA protein contained a 17-amino-acid deletion (positions 58-74) in the stalk region. This deletion pattern was identical to that observed in the H5N1 strain A/brown skua/Torgersen Island/o81-b82/2024 isolated from Torgersen Island, it was absent in 1143 globally sampled H5N1 clade 2.3.4.4b variants. However, continued genomic surveillance is warranted to assess its epidemiological significance. Notably, the Fildes Peninsula isolates harbored an E57K substitution in the NA gene, absent in the Torgersen Island strain. Multiple studies have documented NA stalk deletions in various influenza strains: A/Hong Kong/159/97 (H5N1) possesses a 19-aa deletion (positions 54-72) that reduces viral release capacity; A/chicken/Hubei/327/2004 (H5N1) contains a 20-aa deletion (positions 49-68); while A/Puerto Rico/8/34 (H1N1) and A/WSN/33 (H1N1) exhibit 15-aa (positions 63-77) and 16-aa (positions 57-72) deletions, respectively (Matrosovich et al., 1999; Martin et al., 2009). Reverse genetics studies demonstrate that NA stalk length influences viral replication kinetics -viruses with shorter stalks replicate significantly faster than their long-stalk counterparts. NA stalk length correlates with H5N1 virulence and pathogenicity, as stalk-deletion recombinants display attenuated pathogenicity in ducks(Tumpey et al., 2002). The increasing prevalence of NA stalk deletions in avian H5N1 viral isolates suggests potential adaptive advantages that may enhance cellular fitness and expand host tropism. While these mutations could significantly alter viral pathogenicity, transmissibility, and host specificity, their precise biological consequences require further experimental validation.

This study reports the first detection of HPAI H5N1 in brown skuas on Fildes Peninsula, Antarctic Peninsula, characterized by a unique 17-aa NA stalk deletion (58–74) currently found only in Antarctic strains. The ecological and evolutionary significance of this Antarctic-specific deletion warrants investigation. It should be noted that this study has limitations including restricted species coverage and small sample size. These constraints underscore the need for more systematic and continuous monitoring efforts to obtain more representative samples. Given the increasing human activities in Antarctica and its unique ecological significance, such surveillance is not only critical for understanding viral ecology but also essential for early detection of potential zoonotic transmission risks.

## Acknowledgments

This work was supported by the National Key Research and Development Program of China (2024YFC2813605,2023YFC2605504). We thank the Center for Instrumental Analysis and Metrology and Institutional Center for Shared Technologies and Facilities of the Wuhan Institute of Virology for providing technical assistance.

## Author Bio

Xu Ruifeng is a Research Technician at the Wuhan Institute of Virology, Chinese Academy of Sciences, specializing in virology and pathogenic microorganism studies.

Gao Minhao is a member of the research team at the Polar Research Institute of China. With extensive experience in Antarctic expeditions, he has participated in multiple Chinese National Antarctic Research Expeditions, conducting critical fieldwork in extreme environments.

## Notes

### Competing Interest Statement

The authors have declared no competing interest.

## Reference

Banyard AC, Bennison A, Byrne AMP, Reid SM, Lynton-Jenkins JG, Mollett B, De Silva D, Peers-Dent J, Finlayson K, Hall R, Blockley F, Blyth M, Falchieri M, Fowler Z, Fitzcharles EM, Brown IH, James J. 2024. Detection and spread of high pathogenicity avian influenza virus h5n1 in the antarctic region. Nature Communications, 15.

Bennett-Laso B, Berazay B, Muñoz G, Ariyama N, Enciso N, Braun C, Krüger L, Barták M, González-Aravena M, Neira V. 2024. Confirmation of highly pathogenic avian influenza h5n1 in skuas, antarctica 2024. Frontiers in Veterinary Science, 11.

Caserta LC, Frye EA, Butt SL, Laverack M, Nooruzzaman M, Covaleda LM, Thompson AC, Koscielny MP, Cronk B, Johnson A, Kleinhenz K, Edwards EE, Gomez G, Hitchener G, Martins M, Kapczynski DR, Suarez DL, Alexander Morris ER, Hensley T, Beeby JS, Lejeune M, Swinford AK, Elvinger F, Dimitrov KM, Diel DG. 2024. Spillover of highly pathogenic avian influenza h5n1 virus to dairy cattle. Nature, 634: 669–676.

Garg S, Reinhart K, Couture A, Kniss K, Davis CT, Kirby MK, Murray EL, Zhu S, Kraushaar V, Wadford DA, Drehoff C, Kohnen A, Owen M, Morse J, Eckel S, Goswitz J, Turabelidze G, Krager S, Unutzer A, Gonzales ER, Abdul Hamid C, Ellington S, Mellis AM, Budd A, Barnes JR, Biggerstaff M, Jhung MA, Richmond-Crum M, Burns E, Shimabukuro TT, Uyeki TM, Dugan VG, Reed C, Olsen SJ. 2025. Highly pathogenic avian influenza a(h5n1) virus infections in humans. New England Journal of Medicine, 392: 843–854.

Martin DP, Zhou H, Yu Z, Hu Y, Tu J, Zou W, Peng Y, Zhu J, Li Y, Zhang A, Yu Z, Ye Z, Chen H, Jin M. 2009. The special neuraminidase stalk-motif responsible for increased virulence and pathogenesis of h5n1 influenza a virus. PLoS ONE, 4.

Matrosovich M, Zhou N, Kawaoka Y, Webster R. 1999. The surface glycoproteins of h5 influenza viruses isolated from humans, chickens, and wild aquatic birds have distinguishable properties Journal of Vironal., 73: 1146–1155.

Moore JK, Abbott MR, Richman JG. 1999. Location and dynamics of the antarctic polar front from satellite sea surface temperature data. Journal of Geophysical Research: Oceans, 104: 3059–3073.

Peacock TP, Moncla L, Dudas G, VanInsberghe D, Sukhova K, Lloyd-Smith JO, Worobey M, Lowen AC, Nelson MI. 2024. The global h5n1 influenza panzootic in mammals. Nature, 637: 304–313.

Tumpey TM, Suarez DL, Perkins LEL, Senne DA, Lee J-g, Lee Y-J, Mo I-P, Sung H-W, Swayne DE. 2002. Characterization of a highly pathogenic h5n1 avian influenza a virus isolated from duck meat. Journal of Virology, 76: 6344–6355.

Uhart MM, Vanstreels RET, Nelson MI, Olivera V, Campagna J, Zavattieri V, Lemey P, Campagna C, Falabella V, Rimondi A. 2024. Epidemiological data of an influenza a/h5n1 outbreak in elephant seals in argentina indicates mammal-to-mammal transmission. Nature Communications, 15.

