## Appendix Files for "Transcontinental Spread of HPAI H5N1 from South America to Antarctica via Avian Vectors": Appendix.docx

**Materials and Methods**

**Sample Collection**

During the 41st Chinese Antarctic Research Expedition, field investigations were conducted across Fildes Peninsula and surrounding areas from December 2024 to February 2025. Expedition members wearing full personal protective equipment collected influenza samples from both live animals and atypical mortality cases using standardized protocols for sterilization, disinfection, and hazardous waste disposal. Spatial distribution of sampling sites was mapped using Google earth. A total of 11 biological samples were obtained, including: oropharyngeal swabs, cloacal swabs, fecal samples, brain tissue specimens. All samples were preserved in cryotubes containing 1 mL RNA later and stored at 4°C after collection, and preserved at -80°C.

**qPCR**

For viral genetic testing, we utilized a portable real-time fluorescence PCR system and Influenza A Virus with H5/H7 Subtype Nucleic Acid Detection Kit (PCR-fluorescent probe method) (Zhongkeshengyi science and technology Co., Beijing, China), following the manufacturer’s instructions for avian influenza virus screening. Samples with Ct values ≤ 35, accompanied by a characteristic S-shaped amplification curve, were identified as avian influenza virus-positive.

**RNA Extraction**

The processed samples were subjected to RNA extraction using the MiniBEST Viral RNA/DNA Extraction Kit Ver. 5.0 (Takara, Japan) following the manufacturer's protocol. The extracted RNA was immediately stored at -80°C until downstream molecular analysis. For each extraction procedure, RNase-free water was included as a negative control.

**Next-generation sequencing**

Influenza virus-positive samples identified by PCR-fluorescent probe assay were selected for Next-generation sequencing (NGS) to analyze genomic characteristics. Total RNA libraries were constructed using the VAHTS Universal V8 RNA-seq Library Prep Kit for Illumina (Vazyme, Nanjing, China) in accordance with the manufacturer's protocol. Approximately 1 μg of total RNA was reverse-transcribed into cDNA, followed by fragmentation, adapter ligation, PCR amplification, and purification. Library quality control was performed using an Agilent 2100 Bioanalyzer. Final sequencing was conducted on an Illumina NovaSeq platform (San Diego, CA, USA).

**Sequencing data assembly and bioinformatic analysis**

According to the method described by Zhang et al.(Zhang et al., 2025) and Yu et al.(Yu et al., 2014), we performed bioinformatics analysis on whole-genome sequencing data to assemble sequences of eight different influenza virus segments. Briefly, raw sequencing reads were first subjected to quality assessment and removal of adapter sequences/low-quality reads using Fastp. Quality-controlled sequences were then aligned against a host genome database using Kraken2 to eliminate host-derived sequences. De novo assembly was conducted with Megahit, followed by assembly polishing with Pilon and sequence consensus optimization using CAP3. The improved sequences served as references for reassembling high-quality reads to generate final reference-based assemblies. 514 H5N1 genomic sequences were downloaded from the GISAID database and NCBI, yielding eight datasets corresponding to the eight gene segments of H5N1. Each dataset underwent multiple sequence alignment using MUSCLE, followed by phylogenetic analysis that incorporated the H5N1 sequences obtained in our study. Maximum likelihood phylogenetic trees were constructed using FastTree, with subsequent visualization performed using the ggtree package. For spatiotemporal phylogenetic analysis, we employed Nextstrain to infer the viral evolutionary origin, mutation rate, and transmission patterns.

**Appendix Reference**

Yu X, Jin T, Cui Y, Pu X, Li J, Xu J, Liu G, Jia H, Liu D, Song S, Yu Y, Xie L, Huang R, Ding H, Kou Y, Zhou Y, Wang Y, Xu X, Yin Y, Wang J, Guo C, Yang X, Hu L, Wu X, Wang H, Liu J, Zhao G, Zhou J, Pan J, Gao GF, Yang R, Wang J, Lyles DS. 2014. Influenza h7n9 and h9n2 viruses: Coexistence in poultry linked to human h7n9 infection and genome characteristics. Journal of Virology, 88: 3423-3431.

Zhang N, Hu B, Zhang L, Gan M, Ding Q, Pan K, Wei J, Xu W, Chen D, Zheng S, Cai K, Zheng Z. 2025. Virome landscape of wild rodents and shrews in central china. Microbiome, 13.

**Appendix Figure legends**

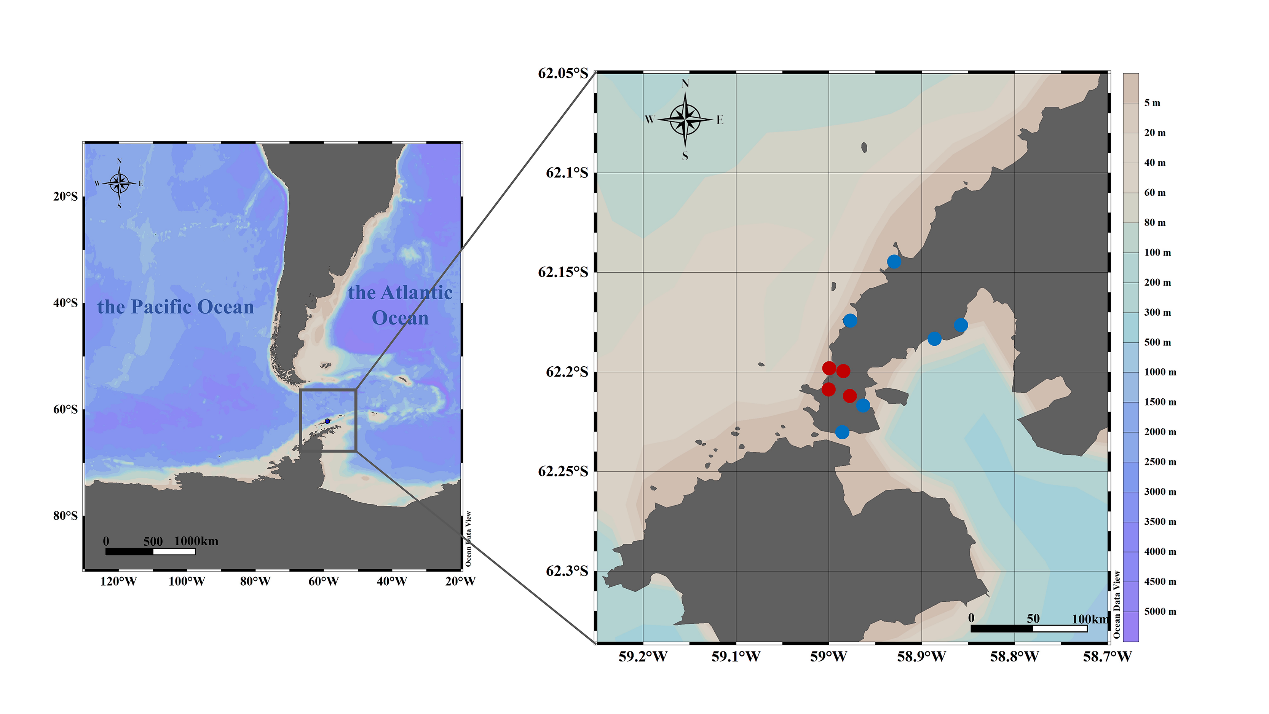

**Appendix Figure S1.** Collection sites sampled. Map of Fildes Peninsula, South Shetland Islands, Antarctic Peninsula showing sampling locations of animal specimens collected in this study. Complete H5N1 virus sequences were obtained from the samples marked in red.

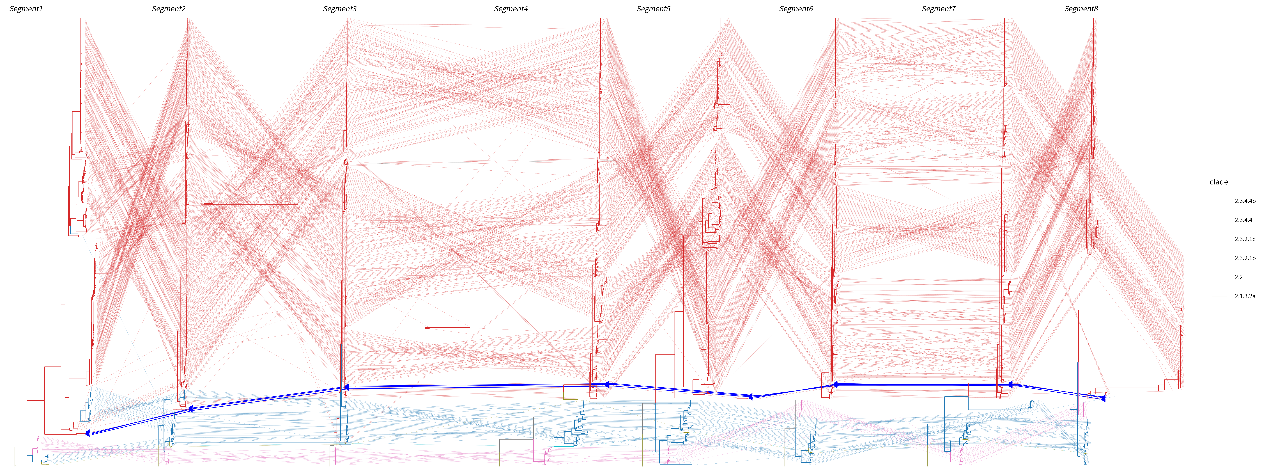

**Appendix Figure S2.** Phylogenetic analysis. A maximum likelihood phylogenetic tree was constructed using 514 representative H5N1 sequences downloaded from the GISAID combined with sequences generated from Fildes Peninsula samples, encompassing all eight genomic segments (PB2, PB1, PA, HA, NP, NA, MP, and NS). Tip labels and connecting branches were color-coded according to the legend. Red denotes subtype 2.3.4.4b, olive green signifies 2.3.4.4, blue indicates 2.3.2.1c, cyan marks 2.3.2.1b, pink symbolizes 2.2, and light olive green represents 2.1.3.2a. Bright blue lines connect all eight genomic segments of the H5N1 strain identified on Fildes Island.

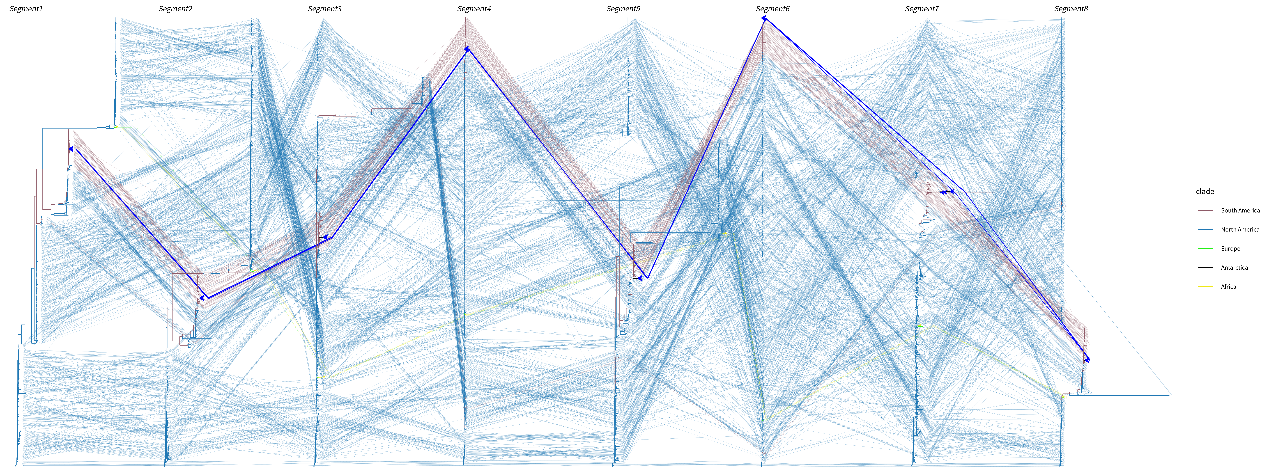

**Appendix Figure S3.** Phylogenetic analysis. A maximum likelihood phylogenetic tree was constructed using 1143 representative H5N1 sequences downloaded from NCBI combined with sequences generated from Fildes Peninsula samples, encompassing all eight genomic segments (PB2, PB1, PA, HA, NP, NA, MP, and NS). The phylogenetic tree's color distinguishes geographic origins with brown indicating South America, blue symbolizing North America, green representing Europe, black denoting Antarctica, and yellow signifying Africa. Bright blue lines connect all eight genomic segments of the H5N1 strain identified on Fildes Island.

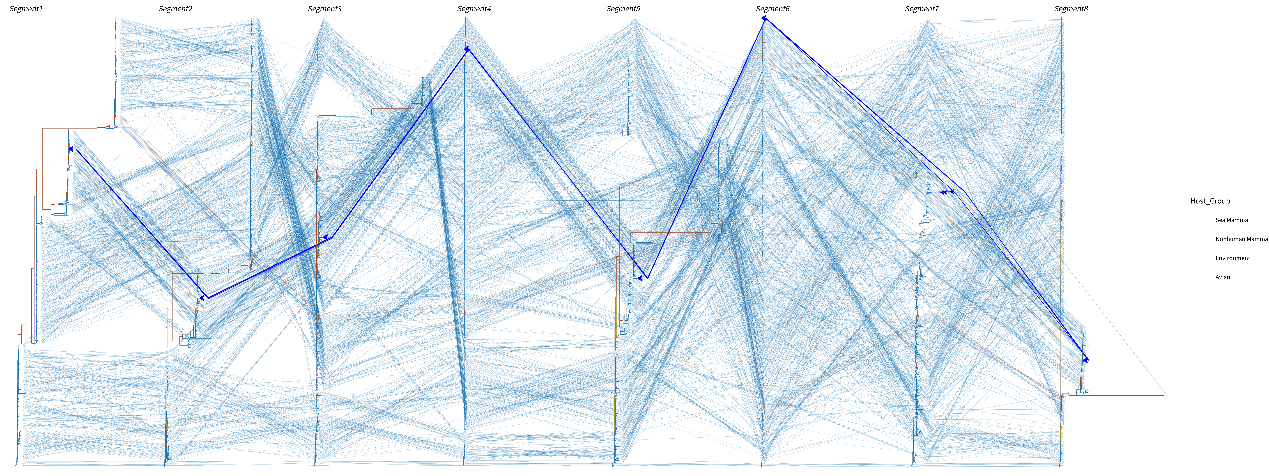

**Appendix Figure S4.** Phylogenetic analysis. A maximum likelihood phylogenetic tree was constructed using 1143 representative H5N1 sequences downloaded from NCBI combined with sequences generated from Fildes Peninsula samples, encompassing all eight genomic segments (PB2, PB1, PA, HA, NP, NA, MP, and NS). The color of the phylogenetic tree classifies host origins: brown corresponds to Sea Mammal, orange reflects Nonhuman Mammal, purple designates Environment, and sky blue characterizes Avian. Bright blue lines connect all eight genomic segments of the H5N1 strain identified on Fildes Island.

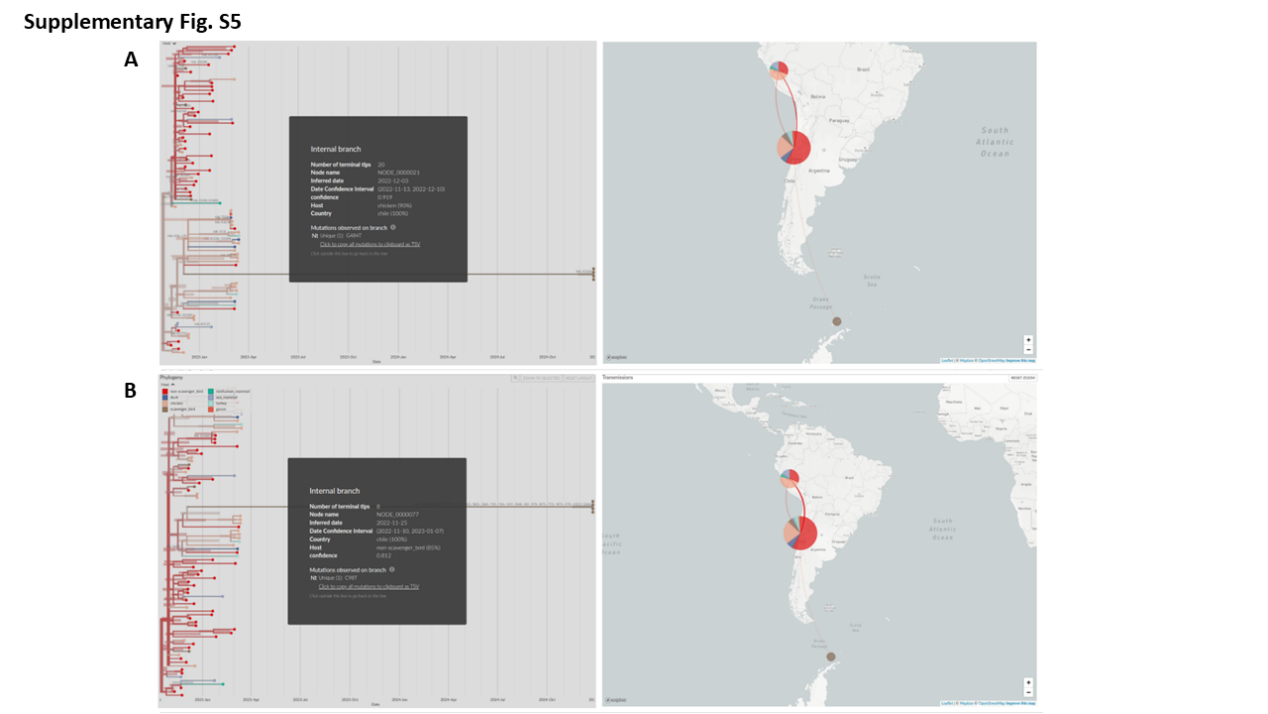

**Appendix Figure S5.** Temporal-geographic-phylogenetic analysis of the HA (A) and NA (B) sequences. The connecting lines indicate potential transmission routes. The color-coded legend illustrates host species classification: red squares denote non-scavenger bird, dark blue squares signify duck, light orange squares represent chicken, brown squares indicate scavenger bird, teal squares symbolize nonhuman mammal, light purple squares correspond to sea mammal, pale cyan squares stand for turkey, and orange-red squares designate goose.
