## Appendix Files for "Transcontinental Spread of HPAI H5N1 from South America to Antarctica via Avian Vectors": AppendixTable1.pdf

**Appendix Table S1.** Detailed information for all the samples collected.

| Number | location | Collection date | Species | sample | Positive | GISAID ID |
|---|---|---|---|---|---|---|
| B-2459 | 62.19919829°S, 58.99613846°W | 2024/12/10 | Brown Skua | Brain tissue | YES | EPI_ISL19847535 |
| B-1225 | 62.198969°S, 58.993061°W | 2024/12/25 | Brown Skua | Brain tissue | YES | EPI_ISL19847536 |
| B-2517 | 62.208744°S, 59.000028°W | 2024/12/26 | Brown Skua | Brain tissue | YES | EPI_ISL19847538 |
| F-2810 | 62.21208001°S, 58.97715781°W | 2024/12/26 | Brown Skua | fecal sample | YES | EPI_ISL19847539 |
| F-2117 | 62.17426349°S, 58.97736702°W | 2025/1/17 | elephant seal | fecal sample | NO | |
| Y-2217 | 62.14466710°S, 58.92984766°W | 2025/1/17 | Fur seal | throat swab | NO | |
| G-2117 | 62.14466710°S, 58.92984766°W | 2025/1/17 | Fur seal | anal swab | NO | |
| F-2122 | 62.23039287°S, 58.98536134°W | 2025/1/22 | Brown Skua | fecal sample | NO | |
| F-2213 | 62.17656773°S, 58.857727090°W | 2025/2/13 | Gentoo Penguin | fecal sample | NO | |
| F-2313 | 62.18331919°S, 58.886164579°W | 2025/2/13 | Southern Giant Petrel | fecal sample | NO | |
| F-2216 | 62.21690135°S, 58.96290321°W | 2025/2/16 | Brown Skua | fecal sample | NO | |
