## Appendix Files for "Transcontinental Spread of HPAI H5N1 from South America to Antarctica via Avian Vectors": AppendixTable2.pdf

**Appendix Table S2.** Detailed metadata of GISAID reference sequences used for genotyping the sequences

| Isolate_Id | Isolate_Name | Clade | Location | Host | Collection_D | Subtype |
|---|---|---|---|---|---|---|
| EPI_ISL_199411 | A/goose/Shandong/03.10_HZ/2015 | 2.3.2.1c | Asia / China | Goose | 2015/3/10 | A / H5N1 |
| EPI_ISL_199413 | A/goose/Shandong/GS/2013 | 2.3.2.1c | Asia / China | Goose | 2013/1/1 | A / H5N1 |
| EPI_ISL_259927 | A/goose/Guangdong/SH7/2013 | 2.3.4.4 | Asia / China / | Goose | 2013/9/20 | A / H5N1 |
| EPI_ISL_12572661 | A/goose/Guizhou/S1541/2022 | 2.3.4.4b | Asia / China / | Goose | 2022/2/22 | A / H5N1 |
| EPI_ISL_12572662 | A/goose/Hunan/SE284/2022 | 2.3.4.4b | Asia / China / | Goose | 2022/1/5 | A / H5N1 |
| EPI_ISL_19033551 | A/domestic_goose/Poland/H48-D1/2 | 2.3.4.4b | Europe / Pola | Goose | 2024/2/13 | A / H5N1 |
| EPI_ISL_19033553 | A/domestic_goose/Poland/H49-W/2 | 2.3.4.4b | Europe / Pola | Goose | 2024/2/13 | A / H5N1 |
| EPI_ISL_201001 | A/Black-necked_Grebe/Inner_Mong | 2.3.2.1c | Asia / China / | Anser ind | 2015/5/12 | A / H5N1 |
| EPI_ISL_201002 | A/Black-necked_Grebe/Inner_Mong | 2.3.2.1c | Asia / China / | Anser ind | 2015/5/12 | A / H5N1 |
| EPI_ISL_201003 | A/Black-necked_Grebe/Inner_Mong | 2.3.2.1c | Asia / China / | Anser ind | 2015/5/12 | A / H5N1 |
| EPI_ISL_201004 | A/Black-necked_Grebe/Inner_Mong | 2.3.2.1c | Asia / China / | Anser ind | 2015/5/12 | A / H5N1 |
| EPI_ISL_201005 | A/Black-necked_Grebe/Inner_Mong | 2.3.2.1c | Asia / China / | Anser ind | 2015/5/12 | A / H5N1 |
| EPI_ISL_200998 | A/Bar-headed_Goose/Qinghai/BTY2 | 2.3.2.1c | Asia / China / | Anser ind | 2015/7/14 | A / H5N1 |
| EPI_ISL_200999 | A/Great_black-headed_Gull/Qingha | 2.3.2.1c | Asia / China / | Anser ind | 2015/7/14 | A / H5N1 |
| EPI_ISL_201000 | A/Great_black-headed_Gull/Qingha | 2.3.2.1c | Asia / China / | Anser ind | 2015/7/14 | A / H5N1 |
| EPI_ISL_201006 | A/Black-necked_Grebe/Inner_Mong | 2.3.2.1c | Asia / China / | Anser ind | 2015/5/12 | A / H5N1 |
| EPI_ISL_201007 | A/Black-necked_Grebe/Inner_Mong | 2.3.2.1c | Asia / China / | Anser ind | 2015/5/12 | A / H5N1 |
| EPI_ISL_201008 | A/Black-necked_Grebe/Inner_Mong | 2.3.2.1c | Asia / China / | Anser ind | 2015/5/12 | A / H5N1 |
| EPI_ISL_201009 | A/Black-necked_Grebe/Inner_Mong | 2.3.2.1c | Asia / China / | Anser ind | 2015/5/12 | A / H5N1 |
| EPI_ISL_201010 | A/Black-necked_Grebe/Inner_Mong | 2.3.2.1c | Asia / China / | Anser ind | 2015/5/12 | A / H5N1 |
| EPI_ISL_14619009 | A/Hartlaubs_gull/South_Africa/210 | 2.3.4.4b | Africa / South | Gull | 2021/5/17 | A / H5N1 |
| EPI_ISL_14637676 | A/Hartlaubs_gull/21050385/2021 | 2.3.4.4b | Africa / South | Gull | 2021/5/20 | A / H5N1 |
| EPI_ISL_14638201 | A/Hartlaubs_gull/South_Africa/210 | 2.3.4.4b | Africa / South | Gull | 2021/5/20 | A / H5N1 |
| EPI_ISL_14639383 | A/Kelp_gull/South_Africa/2105038 | 2.3.4.4b | Africa / South | Gull | 2021/5/20 | A / H5N1 |
| EPI_ISL_14640323 | A/Hartlaubs_gull/South_Africa/210 | 2.3.4.4b | Africa / South | Gull | 2021/5/20 | A / H5N1 |
| EPI_ISL_14918030 | A/ostrich/South_Africa/21060311/2 | 2.3.4.4b | Africa / South | Ostrich | 2021/6/14 | A / H5N1 |
| EPI_ISL_14918305 | A/ostrich/South_Africa/21060357/2 | 2.3.4.4b | Africa / South | Ostrich | 2021/6/21 | A / H5N1 |
| EPI_ISL_14918327 | A/ostrich/South_Africa/21060425/2 | 2.3.4.4b | Africa / South | Ostrich | 2021/6/23 | A / H5N1 |
| EPI_ISL_14933722 | A/ostrich/South_Africa/21070586/2 | 2.3.4.4b | Africa / South | Ostrich | 2021/7/28 | A / H5N1 |
| EPI_ISL_19174942 | A/Swift_tern/South_Africa/2404017 | 2.3.4.4b | Africa / South | Ostrich | 2024/4/22 | A / H5N1 |
| EPI_ISL_4651963 | A/common_pheasant_/Sweden/SVA | 2.3.4.4b | Europe / Swe | Phasanius | 2021/9/22 | A / H5N1 |
| EPI_ISL_19033338 | A/mute_swan/Poland/MB085-N/202 | 2.3.4.4b | Europe / Pola | Swan | 2024/2/8 | A / H5N1 |
| EPI_ISL_19033559 | A/mute_swan/Poland/MB055-L1/20 | 2.3.4.4b | Europe / Pola | Swan | 2024/2/1 | A / H5N1 |
| EPI_ISL_19033560 | A/mute_swan/Poland/MB055-L2/20 | 2.3.4.4b | Europe / Pola | Swan | 2024/2/1 | A / H5N1 |
| EPI_ISL_19033561 | A/mute_swan/Poland/MB055-L4/20 | 2.3.4.4b | Europe / Pola | Swan | 2024/2/1 | A / H5N1 |
| EPI_ISL_19033562 | A/mute_swan/Poland/MB070-L1/20 | 2.3.4.4b | Europe / Pola | Swan | 2024/2/19 | A / H5N1 |
| EPI_ISL_171121 | A/whooper_swan/Henan/SMX1/201 | 2.3.2.1c | Asia / China / | Cygnus c | 2015/1/4 | A / H5N1 |
| EPI_ISL_171141 | A/whooper_swan/Henan/SMX3/201 | 2.3.2.1c | Asia / China / | Cygnus c | 2015/1/4 | A / H5N1 |
| EPI_ISL_171165 | A/whooper_swan/Henan/SMX4/201 | 2.3.2.1c | Asia / China / | Cygnus c | 2015/1/5 | A / H5N1 |
| EPI_ISL_171186 | A/whooper_swan/Henan/SMX9/201 | 2.3.2.1c | Asia / China / | Cygnus c | 2015/1/5 | A / H5N1 |
| EPI_ISL_19603132 | A/mute_swan/Poland/MB418N3/20 | 2.3.4.4b | Europe / Pola | Cygnus o | 2024/10/11 | A / H5N1 |
| EPI_ISL_19603135 | A/mute_swan/Poland/MB418N4/20 | 2.3.4.4b | Europe / Pola | Cygnus o | 2024/10/21 | A / H5N1 |
| EPI_ISL_19603471 | A/mute_swan/Poland/MB422NJ/20 | 2.3.4.4b | Europe / Pola | Cygnus o | 2024/10/26 | A / H5N1 |
| EPI_ISL_19603472 | A/mute_swan/Poland/MB424NJ/20 | 2.3.4.4b | Europe / Pola | Cygnus o | 2024/10/28 | A / H5N1 |
| EPI_ISL_19603485 | A/mute_swan/Poland/MB428NMJ/2 | 2.3.4.4b | Europe / Pola | Cygnus o | 2024/10/31 | A / H5N1 |
| EPI_ISL_19603543 | A/mute_swan/Poland/MB429N/202 | 2.3.4.4b | Europe / Pola | Cygnus o | 2024/11/4 | A / H5N1 |
| EPI_ISL_19033332 | A/turkey/Poland/H68-T2/2024 | 2.3.4.4b | Europe / Pola | Turkey | 2024/2/23 | A / H5N1 |
| EPI_ISL_19033334 | A/turkey/Poland/H75-T1/2024 | 2.3.4.4b | Europe / Pola | Turkey | 2024/2/26 | A / H5N1 |
| EPI_ISL_19033335 | A/turkey/Poland/H80-T3/2024 | 2.3.4.4b | Europe / Pola | Turkey | 2024/2/23 | A / H5N1 |
| EPI_ISL_19033336 | A/turkey/Poland/H81-T1/2024 | 2.3.4.4b | Europe / Pola | Turkey | 2024/2/27 | A / H5N1 |
| EPI_ISL_19033546 | A/turkey/Poland/H38-T4/2024 | 2.3.4.4b | Europe / Pola | Turkey | 2024/2/7 | A / H5N1 |
| EPI_ISL_19033547 | A/turkey/Poland/H40-T2/2024 | 2.3.4.4b | Europe / Pola | Turkey | 2024/2/8 | A / H5N1 |
| EPI_ISL_19033548 | A/turkey/Poland/H43-T2/2024 | 2.3.4.4b | Europe / Pola | Turkey | 2024/2/11 | A / H5N1 |
| EPI_ISL_19033550 | A/turkey/Poland/H47-T4/2024 | 2.3.4.4b | Europe / Pola | Turkey | 2024/2/13 | A / H5N1 |
| EPI_ISL_19033556 | A/turkey/Poland/H63-N/2024 | 2.3.4.4b | Europe / Pola | Turkey | 2024/2/19 | A / H5N1 |
| EPI_ISL_212420 | A/Great_Cormorant/Hubei/VI128/2 | 2.3.2.1c | Asia / China / | Other avi | 2015/2/5 | A / H5N1 |
| EPI_ISL_212421 | A/Great_Cormorant/Hubei/VI132/2 | 2.3.2.1c | Asia / China / | Other avi | 2015/2/5 | A / H5N1 |
| EPI_ISL_212422 | A/Great_Cormorant/Hubei/VI133/2 | 2.3.2.1c | Asia / China / | Other avi | 2015/2/5 | A / H5N1 |
| EPI_ISL_14639837 | A/Spotted_eagle_owl/South_Africa/ | 2.3.4.4b | Africa / South | Other avi | 2021/5/20 | A / H5N1 |
| EPI_ISL_14645331 | A/Black-headed_heron/South_Afric | 2.3.4.4b | Africa / South | Other avi | 2021/5/25 | A / H5N1 |

| | | | | | | |
|---|---|---|---|---|---|---|
| EPI_ISL_14973559 | A/Cape_gannet/South_Africa/21100 | 2.3.4.4b | Africa / South | Other avi | 2021/10/7 | A / H5N1 |
| EPI_ISL_12572664 | A/wild_duck/Hebei/SD012/2021 | 2.3.4.4b | Asia / China / | Other avi | 2021/11/25 | A / H5N1 |
| EPI_ISL_19603087 | A/blackbird/Poland/MB406M/2024 | 2.3.4.4b | Europe / Pola | Other avi | 2024/10/18 | A / H5N1 |
| EPI_ISL_199157 | A/environment/Sichuan/12.11_CDL | 2.3.2.1c | Asia / China | Environm | 2014/12/11 | A / H5N1 |
| EPI_ISL_199286 | A/environment/Jiangsu/01.20_TCC | 2.3.2.1c | Asia / China | Environm | 2015/1/20 | A / H5N1 |
| EPI_ISL_198753 | A/environment/Jilin/04.25_NA001/ | 2.3.2.1c | Asia / China | Environm | 2015/4/25 | A / H5N1 |
| EPI_ISL_198842 | A/environment/Jilin/04.25_MHK01 | 2.3.2.1c | Asia / China | Environm | 2015/4/25 | A / H5N1 |
| EPI_ISL_198848 | A/environment/Jilin/04.25_MHK01 | 2.3.2.1c | Asia / China | Environm | 2015/4/25 | A / H5N1 |
| EPI_ISL_17655676 | A/environment/Kagoshima/KU-B20 | 2.3.4.4b | Asia / Japan | Environm | 2021/12/20 | A / H5N1 |
| EPI_ISL_18472731 | A/environment/Kagoshima/KU-6A/ | 2.3.4.4b | Asia / Japan | Environm | 2022/11/1 | A / H5N1 |
| EPI_ISL_18472732 | A/environment/Kagoshima/KU-B1/ | 2.3.4.4b | Asia / Japan | Environm | 2022/11/2 | A / H5N1 |
| EPI_ISL_18472733 | A/environment/Kagoshima/KU-B2/ | 2.3.4.4b | Asia / Japan | Environm | 2022/11/3 | A / H5N1 |
| EPI_ISL_18472734 | A/environment/Kagoshima/KU-B3/ | 2.3.4.4b | Asia / Japan | Environm | 2022/11/3 | A / H5N1 |
| EPI_ISL_18472735 | A/environment/Kagoshima/KU-B4/ | 2.3.4.4b | Asia / Japan | Environm | 2022/11/4 | A / H5N1 |
| EPI_ISL_18472736 | A/environment/Kagoshima/KU-G1/ | 2.3.4.4b | Asia / Japan | Environm | 2022/11/4 | A / H5N1 |
| EPI_ISL_18472737 | A/environment/Kagoshima/KU-G2/ | 2.3.4.4b | Asia / Japan | Environm | 2022/11/4 | A / H5N1 |
| EPI_ISL_18472738 | A/environment/Kagoshima/KU-G3/ | 2.3.4.4b | Asia / Japan | Environm | 2022/11/4 | A / H5N1 |
| EPI_ISL_18472739 | A/environment/Kagoshima/KU-B5/ | 2.3.4.4b | Asia / Japan | Environm | 2022/11/5 | A / H5N1 |
| EPI_ISL_18472740 | A/environment/Kagoshima/KU-B6/ | 2.3.4.4b | Asia / Japan | Environm | 2022/11/5 | A / H5N1 |
| EPI_ISL_18472741 | A/environment/Kagoshima/KU-B7/ | 2.3.4.4b | Asia / Japan | Environm | 2022/11/5 | A / H5N1 |
| EPI_ISL_18472742 | A/environment/Kagoshima/KU-G5/ | 2.3.4.4b | Asia / Japan | Environm | 2022/11/7 | A / H5N1 |
| EPI_ISL_18472743 | A/environment/Kagoshima/KU-H1/ | 2.3.4.4b | Asia / Japan | Environm | 2022/11/7 | A / H5N1 |
| EPI_ISL_18472744 | A/environment/Kagoshima/KU-B11 | 2.3.4.4b | Asia / Japan | Environm | 2022/11/8 | A / H5N1 |
| EPI_ISL_18472749 | A/environment/Kagoshima/KU-J7/2 | 2.3.4.4b | Asia / Japan | Environm | 2022/11/9 | A / H5N1 |
| EPI_ISL_18472750 | A/environment/Kagoshima/KU-J8/2 | 2.3.4.4b | Asia / Japan | Environm | 2022/11/9 | A / H5N1 |
| EPI_ISL_18472756 | A/environment/Kagoshima/KU-B8/ | 2.3.4.4b | Asia / Japan | Environm | 2022/11/6 | A / H5N1 |
| EPI_ISL_18472757 | A/environment/Kagoshima/KU-D1/ | 2.3.4.4b | Asia / Japan | Environm | 2022/11/6 | A / H5N1 |
| EPI_ISL_18472758 | A/environment/Kagoshima/KU-D2/ | 2.3.4.4b | Asia / Japan | Environm | 2022/11/6 | A / H5N1 |
| EPI_ISL_18472759 | A/environment/Kagoshima/KU-D3/ | 2.3.4.4b | Asia / Japan | Environm | 2022/11/6 | A / H5N1 |
| EPI_ISL_18472760 | A/environment/Kagoshima/KU-D4/ | 2.3.4.4b | Asia / Japan | Environm | 2022/11/6 | A / H5N1 |
| EPI_ISL_18472761 | A/environment/Kagoshima/KU-G4/ | 2.3.4.4b | Asia / Japan | Environm | 2022/11/7 | A / H5N1 |
| EPI_ISL_18472762 | A/environment/Kagoshima/KU-D6/ | 2.3.4.4b | Asia / Japan | Environm | 2022/11/7 | A / H5N1 |
| EPI_ISL_18472764 | A/environment/Kagoshima/KU-D7/ | 2.3.4.4b | Asia / Japan | Environm | 2022/11/8 | A / H5N1 |
| EPI_ISL_18509885 | A/environment/Kagoshima/KU-B3/ | 2.3.4.4b | Asia / Japan | Environm | 2023/11/7 | A / H5N1 |
| EPI_ISL_18509886 | A/environment/Kagoshima/KU-C1/ | 2.3.4.4b | Asia / Japan | Environm | 2023/11/7 | A / H5N1 |
| EPI_ISL_18509887 | A/environment/Kagoshima/KU-C2/ | 2.3.4.4b | Asia / Japan | Environm | 2023/11/7 | A / H5N1 |
| EPI_ISL_18509888 | A/environment/Kagoshima/KU-C3/ | 2.3.4.4b | Asia / Japan | Environm | 2023/11/7 | A / H5N1 |
| EPI_ISL_18509889 | A/environment/Kagoshima/KU-C4/ | 2.3.4.4b | Asia / Japan | Environm | 2023/11/7 | A / H5N1 |
| EPI_ISL_18509890 | A/environment/Kagoshima/KU-D4/ | 2.3.4.4b | Asia / Japan | Environm | 2023/11/7 | A / H5N1 |
| EPI_ISL_18509891 | A/environment/Kagoshima/KU-E1/ | 2.3.4.4b | Asia / Japan | Environm | 2023/11/7 | A / H5N1 |
| EPI_ISL_18509892 | A/environment/Kagoshima/KU-G4/ | 2.3.4.4b | Asia / Japan | Environm | 2023/11/7 | A / H5N1 |
| EPI_ISL_18509893 | A/environment/Kagoshima/KU-H1/ | 2.3.4.4b | Asia / Japan | Environm | 2023/11/7 | A / H5N1 |
| EPI_ISL_18509894 | A/environment/Kagoshima/KU-B2/ | 2.3.4.4b | Asia / Japan | Environm | 2023/11/7 | A / H5N1 |
| EPI_ISL_18509898 | A/environment/Kagoshima/KU-H2/ | 2.3.4.4b | Asia / Japan | Environm | 2023/11/7 | A / H5N1 |
| EPI_ISL_18612259 | A/environment/Kagoshima/KU-D6/ | 2.3.4.4b | Asia / Japan | Environm | 2023/11/20 | A / H5N1 |
| EPI_ISL_18615595 | A/environment/Kagoshima/KU-D1/ | 2.3.4.4b | Asia / Japan / | Environm | 2023/11/6 | A / H5N1 |
| EPI_ISL_18615596 | A/environment/Kagoshima/KU-D2/ | 2.3.4.4b | Asia / Japan | Environm | 2023/11/7 | A / H5N1 |
| EPI_ISL_18615597 | A/environment/Kagoshima/KU-D3/ | 2.3.4.4b | Asia / Japan | Environm | 2023/11/6 | A / H5N1 |
| EPI_ISL_18615598 | A/environment/Kagoshima/KU-G1/ | 2.3.4.4b | Asia / Japan / | Environm | 2023/11/6 | A / H5N1 |
| EPI_ISL_18615599 | A/environment/Kagoshima/KU-G2/ | 2.3.4.4b | Asia / Japan | Environm | 2023/11/6 | A / H5N1 |
| EPI_ISL_18615600 | A/environment/Kagoshima/KU-G3/ | 2.3.4.4b | Asia / Japan | Environm | 2023/11/6 | A / H5N1 |
| EPI_ISL_18634686 | A/environment/Kagoshima/KU-B4- | 2.3.4.4b | Asia / Japan / | Environm | 2023/11/13 | A / H5N1 |
| EPI_ISL_18634687 | A/environment/Kagoshima/KU-G5- | 2.3.4.4b | Asia / Japan / | Environm | 2023/11/13 | A / H5N1 |
| EPI_ISL_18634688 | A/environment/Kagoshima/KU-H3/ | 2.3.4.4b | Asia / Japan / | Environm | 2023/11/27 | A / H5N1 |
| EPI_ISL_18651023 | A/environment/Kagoshima/KU-D11 | 2.3.4.4b | Asia / Japan | Environm | 2023/12/4 | A / H5N1 |
| EPI_ISL_18651569 | A/environment/Kagoshima/KU-G8/ | 2.3.4.4b | Asia / Japan | Environm | 2023/12/4 | A / H5N1 |
| EPI_ISL_18651570 | A/environment/Kagoshima/KU-H4/ | 2.3.4.4b | Asia / Japan | Environm | 2023/12/4 | A / H5N1 |
| EPI_ISL_18651571 | A/environment/Kagoshima/KU-I1/2 | 2.3.4.4b | Asia / Japan | Environm | 2023/12/4 | A / H5N1 |
| EPI_ISL_18651572 | A/environment/Kagoshima/KU-I2/2 | 2.3.4.4b | Asia / Japan | Environm | 2023/12/4 | A / H5N1 |
| EPI_ISL_18770555 | A/environment/Kagoshima/KU-E3/2 | 2.3.4.4b | Asia / Japan | Environm | 2023/12/19 | A / H5N1 |
| EPI_ISL_18770597 | A/environment/Kagoshima/KU-B11 | 2.3.4.4b | Asia / Japan / | Environm | 2023/12/18 | A / H5N1 |
| EPI_ISL_18935776 | A/environment/Kagoshima/KU-B10 | 2.3.4.4b | Asia / Japan | Environm | 2023/12/11 | A / H5N1 |

| | | | | | | |
|---|---|---|---|---|---|---|
| EPI_ISL_18935782 | A/environment/Kagoshima/KU-C7/2 | 2.3.4.4b | Asia / Japan | Environm | 2023/12/11 | A / H5N1 |
| EPI_ISL_19624044 | A/environment/Kagoshima/KU-24-I | 2.3.4.4b | Asia / Japan | Environm | 2024/11/4 | A / H5N1 |
| EPI_ISL_19624045 | A/environment/Kagoshima/KU-24-I | 2.3.4.4b | Asia / Japan | Environm | 2024/11/4 | A / H5N1 |
| EPI_ISL_19624046 | A/environment/Kagoshima/KU-24-I | 2.3.4.4b | Asia / Japan | Environm | 2024/11/4 | A / H5N1 |
| EPI_ISL_19624047 | A/environment/Kagoshima/KU-24-I | 2.3.4.4b | Asia / Japan | Environm | 2024/11/11 | A / H5N1 |
| EPI_ISL_19624050 | A/environment/Kagoshima/KU-24-I | 2.3.4.4b | Asia / Japan | Environm | 2024/11/4 | A / H5N1 |
| EPI_ISL_19624051 | A/environment/Kagoshima/KU-24-I | 2.3.4.4b | Asia / Japan | Environm | 2024/11/4 | A / H5N1 |
| EPI_ISL_19624052 | A/environment/Kagoshima/KU-24-I | 2.3.4.4b | Asia / Japan | Environm | 2024/11/4 | A / H5N1 |
| EPI_ISL_19624053 | A/environment/Kagoshima/KU-24-C | 2.3.4.4b | Asia / Japan | Environm | 2024/11/4 | A / H5N1 |
| EPI_ISL_19624054 | A/environment/Kagoshima/KU-24-C | 2.3.4.4b | Asia / Japan | Environm | 2024/11/4 | A / H5N1 |
| EPI_ISL_19624055 | A/environment/Kagoshima/KU-24-4 | 2.3.4.4b | Asia / Japan | Environm | 2024/11/11 | A / H5N1 |
| EPI_ISL_19624056 | A/environment/Kagoshima/KU-24-4 | 2.3.4.4b | Asia / Japan | Environm | 2024/11/11 | A / H5N1 |
| EPI_ISL_19624057 | A/environment/Kagoshima/KU-24-5 | 2.3.4.4b | Asia / Japan | Environm | 2024/11/11 | A / H5N1 |
| EPI_ISL_19645343 | A/environment/Kagoshima/KU-24-6 | 2.3.4.4b | Asia / Japan | Environm | 2024/11/11 | A / H5N1 |
| EPI_ISL_19645344 | A/environment/Kagoshima/KU-24-I | 2.3.4.4b | Asia / Japan | Environm | 2024/11/11 | A / H5N1 |
| EPI_ISL_19645345 | A/environment/Kagoshima/KU-24-C | 2.3.4.4b | Asia / Japan | Environm | 2024/11/11 | A / H5N1 |
| EPI_ISL_19645350 | A/environment/Kagoshima/KU-24-C | 2.3.4.4b | Asia / Japan | Environm | 2024/11/18 | A / H5N1 |
| EPI_ISL_19645351 | A/environment/Kagoshima/KU-24-J | 2.3.4.4b | Asia / Japan | Environm | 2024/11/11 | A / H5N1 |
| EPI_ISL_19645354 | A/environment/Kagoshima/KU-24-C | 2.3.4.4b | Asia / Japan | Environm | 2024/11/11 | A / H5N1 |
| EPI_ISL_19645356 | A/environment/Kagoshima/KU-24-J | 2.3.4.4b | Asia / Japan | Environm | 2024/11/11 | A / H5N1 |
| EPI_ISL_19645358 | A/environment/Kagoshima/KU-24-J | 2.3.4.4b | Asia / Japan | Environm | 2024/11/11 | A / H5N1 |
| EPI_ISL_19700404 | A/environment/Kagoshima/KU-24-C | 2.3.4.4b | Asia / Japan | Environm | 2024/11/4 | A / H5N1 |
| EPI_ISL_18529945 | A/environment/Kagoshima/KU-D5/2 | 2.3.4.4b | Asia / Japan | Environm | 2023/11/14 | A / H5N1 |
| EPI_ISL_18509834 | A/environment/Kagoshima/KU-B1/2 | 2.3.4.4b | Asia / Japan / | Water sar | 2023/11/7 | A / H5N1 |
| EPI_ISL_19534182 | A/environment/Kagoshima/KU-24B | 2.3.4.4b | Asia / Japan | Water sar | 2024/11/4 | A / H5N1 |
| EPI_ISL_19534183 | A/environment/Kagoshima/KU-24C1/2024 | | Asia / Japan / | Water sar | 2024/11/4 | A / H5N1 |
| EPI_ISL_19534184 | A/environment/Kagoshima/KU-24D | 2.3.4.4b | Asia / Japan / | Water sar | 2024/11/4 | A / H5N1 |
| EPI_ISL_19534185 | A/environment/Kagoshima/KU-24G | 2.3.4.4b | Asia / Japan | Water sar | 2024/11/4 | A / H5N1 |
| EPI_ISL_171297 | A/enviroment/Henan/SMX1/2015 | 2.3.2.1c | Asia / China / | Feces | 2015/1/5 | A / H5N1 |
| EPI_ISL_142593 | A/environment/Hangzhou/109-2/201 | 2.3.2.1c | Asia / China / | Feces | 2013/4/12 | A / H5N1 |
| EPI_ISL_81453 | A/MDCK/Germany/P18-escape/200 | 2.2 | Europe / Gern | Laborator | 2008 | A / H5N1 |
| EPI_ISL_81454 | A/MDCK/Germany/P30-escape/200 | 2.2 | Europe / Gern | Laborator | 2008 | A / H5N1 |
| EPI_ISL_81455 | A/MDCK/Germany/P50-escape/200 | 2.2 | Europe / Gern | Laborator | 2008 | A / H5N1 |
| EPI_ISL_81456 | A/MDCK/Germany/P100b-escape/2 | 2.2 | Europe / Gern | Laborator | 2009 | A / H5N1 |
| EPI_ISL_81457 | A/MDCK/Germany/PP100b-escape/ | 2.2 | Europe / Gern | Laborator | 2009 | A / H5N1 |
| EPI_ISL_81458 | A/MDCK/Germany/PP100c-escape/ | 2.2 | Europe / Gern | Laborator | 2009 | A / H5N1 |
| EPI_ISL_81459 | A/MDCK/Germany/P100c-escape/2 | 2.2 | Europe / Gern | Laborator | 2009 | A / H5N1 |
| EPI_ISL_81460 | A/MDCK/Germany/CoP100-control | 2.2 | Europe / Gern | Laborator | 2009 | A / H5N1 |
| EPI_ISL_81461 | A/MDCK/Germany/CoP50-control/2 | 2.2 | Europe / Gern | Laborator | 2008 | A / H5N1 |
| EPI_ISL_81462 | A/MDCK/Germany/Q18-escape/200 | 2.2 | Europe / Gern | Laborator | 2009 | A / H5N1 |
| EPI_ISL_81463 | A/MDCK/Germany/Q30-escape/200 | 2.2 | Europe / Gern | Laborator | 2009 | A / H5N1 |
| EPI_ISL_81464 | A/MDCK/Germany/Q50-escape/200 | 2.2 | Europe / Gern | Laborator | 2009 | A / H5N1 |
| EPI_ISL_81465 | A/MDCK/Germany/Q100b-escape/2 | 2.2 | Europe / Gern | Laborator | 2009 | A / H5N1 |
| EPI_ISL_81466 | A/MDCK/Germany/QQ100b-escape | 2.2 | Europe / Gern | Laborator | 2009 | A / H5N1 |
| EPI_ISL_81467 | A/MDCK/Germany/QQ100a-escape | 2.2 | Europe / Gern | Laborator | 2009 | A / H5N1 |
| EPI_ISL_81468 | A/MDCK/Germany/Q100c-escape/2 | 2.2 | Europe / Gern | Laborator | 2009 | A / H5N1 |
| EPI_ISL_81469 | A/MDCK/Germany/CoQ100-contro | 2.2 | Europe / Gern | Laborator | 2009 | A / H5N1 |
| EPI_ISL_81470 | A/MDCK/Germany/CoQ50-control/ | 2.2 | Europe / Gern | Laborator | 2009 | A / H5N1 |
| EPI_ISL_88043 | A/cygnus_cygnus/Germany/R65.1/2 | 2.2 | Europe / Gern | Laborator | 2008 | A / H5N1 |
| EPI_ISL_97050 | A/hens_egg/Germany/EscEgg50A-e | 2.2 | Europe / Gern | Laborator | 2009 | A / H5N1 |
| EPI_ISL_97051 | A/hens_egg/Germany/CoJ50-control | 2.2 | Europe / Gern | Laborator | 2009 | A / H5N1 |
| EPI_ISL_17075747 | A/Jiangsu/NJ210/2023 | 2.3.4.4b | Asia / China / | Human | 2023/2/10 | A / H5N1 |
| EPI_ISL_98854 | A/Indonesia/NIHRD11767/2011 | 2.1.3.2a | Asia / Indones | Human | 2011/10/7 | A / H5N1 |
| EPI_ISL_98855 | A/Indonesia/NIHRD11771/2011 | 2.1.3.2a | Asia / Indones | Human | 2011/10/7 | A / H5N1 |
| EPI_ISL_100272 | A/Guangdong-Shenzhen/1/2011 | 2.3.2.1b | Asia / China / | Human | 2011/12/28 | A / H5N1 |
| EPI_ISL_198892 | A/pigeon/Jilin/04.25_CCHL019-O/2 | 2.3.2.1c | Asia / China | Avian | 2015/4/25 | A / H5N1 |
| EPI_ISL_199060 | A/Phalacrocorax/Hubei/01.09_V1/2 | 2.3.2.1c | Asia / China | Avian | 2015/1/9 | A / H5N1 |
| EPI_ISL_199061 | A/Phalacrocorax/Hubei/01.09_V4/2 | 2.3.2.1c | Asia / China | Avian | 2015/1/9 | A / H5N1 |
| EPI_ISL_18472583 | A/hooded_crane/Kagoshima/KU-65/ | 2.3.4.4b | Asia / Japan | Avian | 2022/11/14 | A / H5N1 |
| EPI_ISL_18472584 | A/hooded_crane/Kagoshima/KU-86/ | 2.3.4.4b | Asia / Japan | Avian | 2022/11/19 | A / H5N1 |
| EPI_ISL_18472585 | A/white-naped_crane/Kagoshima/K | 2.3.4.4b | Asia / Japan | Avian | 2022/11/19 | A / H5N1 |
| EPI_ISL_18472586 | A/hooded_crane/Kagoshima/KU-88/ | 2.3.4.4b | Asia / Japan | Avian | 2022/11/19 | A / H5N1 |

| | | | | | | |
|---|---|---|---|---|---|---|
| EPI_ISL_18472587 | A/white-naped_crane/Kagoshima/K | 2.3.4.4b | Asia / Japan | Avian | 2022/12/7 | A / H5N1 |
| EPI_ISL_18472588 | A/hooded_crane/Kagoshima/KU-26 | 2.3.4.4b | Asia / Japan | Avian | 2023/1/22 | A / H5N1 |
| EPI_ISL_18472589 | A/hooded_crane/Kagoshima/KU-2/2 | 2.3.4.4b | Asia / Japan | Avian | 2022/11/1 | A / H5N1 |
| EPI_ISL_18472590 | A/hooded_crane/Kagoshima/KU-3/2 | 2.3.4.4b | Asia / Japan | Avian | 2022/11/2 | A / H5N1 |
| EPI_ISL_18472591 | A/hooded_crane/Kagoshima/KU-4/2 | 2.3.4.4b | Asia / Japan | Avian | 2022/11/3 | A / H5N1 |
| EPI_ISL_18472592 | A/hooded_crane/Kagoshima/KU-5/2 | 2.3.4.4b | Asia / Japan | Avian | 2022/11/3 | A / H5N1 |
| EPI_ISL_18472593 | A/hooded_crane/Kagoshima/KU-6/2 | 2.3.4.4b | Asia / Japan | Avian | 2022/11/4 | A / H5N1 |
| EPI_ISL_18472594 | A/hooded_crane/Kagoshima/KU-7/2 | 2.3.4.4b | Asia / Japan | Avian | 2022/11/4 | A / H5N1 |
| EPI_ISL_18472595 | A/hooded_crane/Kagoshima/KU-8/2 | 2.3.4.4b | Asia / Japan | Avian | 2022/11/4 | A / H5N1 |
| EPI_ISL_18472596 | A/hooded_crane/Kagoshima/KU-9/2 | 2.3.4.4b | Asia / Japan | Avian | 2022/11/4 | A / H5N1 |
| EPI_ISL_18472597 | A/hooded_crane/Kagoshima/KU-10/ | 2.3.4.4b | Asia / Japan | Avian | 2022/11/5 | A / H5N1 |
| EPI_ISL_18472598 | A/hooded_crane/Kagoshima/KU-11/ | 2.3.4.4b | Asia / Japan | Avian | 2022/11/5 | A / H5N1 |
| EPI_ISL_18472599 | A/hooded_crane/Kagoshima/KU-12/ | 2.3.4.4b | Asia / Japan | Avian | 2022/11/5 | A / H5N1 |
| EPI_ISL_18472600 | A/hooded_crane/Kagoshima/KU-13/ | 2.3.4.4b | Asia / Japan | Avian | 2022/11/6 | A / H5N1 |
| EPI_ISL_18472601 | A/hooded_crane/Kagoshima/KU-14/ | 2.3.4.4b | Asia / Japan | Avian | 2022/11/6 | A / H5N1 |
| EPI_ISL_18472602 | A/hooded_crane/Kagoshima/KU-15/ | 2.3.4.4b | Asia / Japan | Avian | 2022/11/6 | A / H5N1 |
| EPI_ISL_18472603 | A/hooded_crane/Kagoshima/KU-16/ | 2.3.4.4b | Asia / Japan | Avian | 2022/11/6 | A / H5N1 |
| EPI_ISL_18472604 | A/hooded_crane/Kagoshima/KU-19/ | 2.3.4.4b | Asia / Japan | Avian | 2022/11/6 | A / H5N1 |
| EPI_ISL_18472605 | A/hooded_crane/Kagoshima/KU-20/ | 2.3.4.4b | Asia / Japan | Avian | 2022/11/7 | A / H5N1 |
| EPI_ISL_18472606 | A/hooded_crane/Kagoshima/KU-22/ | 2.3.4.4b | Asia / Japan | Avian | 2022/11/7 | A / H5N1 |
| EPI_ISL_18472607 | A/hooded_crane/Kagoshima/KU-23/ | 2.3.4.4b | Asia / Japan | Avian | 2022/11/7 | A / H5N1 |
| EPI_ISL_18472608 | A/hooded_crane/Kagoshima/KU-24/ | 2.3.4.4b | Asia / Japan | Avian | 2022/11/7 | A / H5N1 |
| EPI_ISL_18472609 | A/hooded_crane/Kagoshima/KU-26/ | 2.3.4.4b | Asia / Japan | Avian | 2022/11/8 | A / H5N1 |
| EPI_ISL_18472610 | A/hooded_crane/Kagoshima/KU-27/ | 2.3.4.4b | Asia / Japan | Avian | 2022/11/8 | A / H5N1 |
| EPI_ISL_18472611 | A/hooded_crane/Kagoshima/KU-28/ | 2.3.4.4b | Asia / Japan | Avian | 2022/11/8 | A / H5N1 |
| EPI_ISL_18472612 | A/hooded_crane/Kagoshima/KU-29/ | 2.3.4.4b | Asia / Japan | Avian | 2022/11/8 | A / H5N1 |
| EPI_ISL_18472613 | A/hooded_crane/Kagoshima/KU-30/ | 2.3.4.4b | Asia / Japan | Avian | 2022/11/8 | A / H5N1 |
| EPI_ISL_18472614 | A/hooded_crane/Kagoshima/KU-31/ | 2.3.4.4b | Asia / Japan | Avian | 2022/11/8 | A / H5N1 |
| EPI_ISL_18472615 | A/hooded_crane/Kagoshima/KU-32/ | 2.3.4.4b | Asia / Japan | Avian | 2022/11/8 | A / H5N1 |
| EPI_ISL_18472616 | A/hooded_crane/Kagoshima/KU-33/ | 2.3.4.4b | Asia / Japan | Avian | 2022/11/9 | A / H5N1 |
| EPI_ISL_18472617 | A/hooded_crane/Kagoshima/KU-35/ | 2.3.4.4b | Asia / Japan | Avian | 2022/11/9 | A / H5N1 |
| EPI_ISL_18472618 | A/hooded_crane/Kagoshima/KU-36/ | 2.3.4.4b | Asia / Japan | Avian | 2022/11/9 | A / H5N1 |
| EPI_ISL_18472619 | A/hooded_crane/Kagoshima/KU-38/ | 2.3.4.4b | Asia / Japan | Avian | 2022/11/9 | A / H5N1 |
| EPI_ISL_18472620 | A/hooded_crane/Kagoshima/KU-39/ | 2.3.4.4b | Asia / Japan | Avian | 2022/11/9 | A / H5N1 |
| EPI_ISL_18472621 | A/hooded_crane/Kagoshima/KU-40/ | 2.3.4.4b | Asia / Japan | Avian | 2022/11/10 | A / H5N1 |
| EPI_ISL_18472622 | A/hooded_crane/Kagoshima/KU-41/ | 2.3.4.4b | Asia / Japan | Avian | 2022/11/10 | A / H5N1 |
| EPI_ISL_18472623 | A/hooded_crane/Kagoshima/KU-42/ | 2.3.4.4b | Asia / Japan | Avian | 2022/11/10 | A / H5N1 |
| EPI_ISL_18472624 | A/hooded_crane/Kagoshima/KU-43/ | 2.3.4.4b | Asia / Japan | Avian | 2022/11/10 | A / H5N1 |
| EPI_ISL_18472625 | A/hooded_crane/Kagoshima/KU-44/ | 2.3.4.4b | Asia / Japan | Avian | 2022/11/10 | A / H5N1 |
| EPI_ISL_18472626 | A/hooded_crane/Kagoshima/KU-45/ | 2.3.4.4b | Asia / Japan | Avian | 2022/11/10 | A / H5N1 |
| EPI_ISL_18472627 | A/hooded_crane/Kagoshima/KU-46/ | 2.3.4.4b | Asia / Japan | Avian | 2022/11/10 | A / H5N1 |
| EPI_ISL_18472628 | A/hooded_crane/Kagoshima/KU-47/ | 2.3.4.4b | Asia / Japan | Avian | 2022/11/10 | A / H5N1 |
| EPI_ISL_18472629 | A/hooded_crane/Kagoshima/KU-48/ | 2.3.4.4b | Asia / Japan | Avian | 2022/11/10 | A / H5N1 |
| EPI_ISL_18472630 | A/hooded_crane/Kagoshima/KU-50/ | 2.3.4.4b | Asia / Japan | Avian | 2022/11/10 | A / H5N1 |
| EPI_ISL_18472631 | A/hooded_crane/Kagoshima/KU-51/ | 2.3.4.4b | Asia / Japan | Avian | 2022/11/11 | A / H5N1 |
| EPI_ISL_18472632 | A/hooded_crane/Kagoshima/KU-52/ | 2.3.4.4b | Asia / Japan | Avian | 2022/11/11 | A / H5N1 |
| EPI_ISL_18472633 | A/hooded_crane/Kagoshima/KU-53/ | 2.3.4.4b | Asia / Japan | Avian | 2022/11/11 | A / H5N1 |
| EPI_ISL_18472634 | A/hooded_crane/Kagoshima/KU-54/ | 2.3.4.4b | Asia / Japan | Avian | 2022/11/12 | A / H5N1 |
| EPI_ISL_18472635 | A/hooded_crane/Kagoshima/KU-55/ | 2.3.4.4b | Asia / Japan | Avian | 2022/11/12 | A / H5N1 |
| EPI_ISL_18472636 | A/hooded_crane/Kagoshima/KU-56/ | 2.3.4.4b | Asia / Japan | Avian | 2022/11/12 | A / H5N1 |
| EPI_ISL_18472637 | A/hooded_crane/Kagoshima/KU-57/ | 2.3.4.4b | Asia / Japan | Avian | 2022/11/12 | A / H5N1 |
| EPI_ISL_18472638 | A/hooded_crane/Kagoshima/KU-58/ | 2.3.4.4b | Asia / Japan | Avian | 2022/11/13 | A / H5N1 |
| EPI_ISL_18472639 | A/hooded_crane/Kagoshima/KU-59/ | 2.3.4.4b | Asia / Japan | Avian | 2022/11/13 | A / H5N1 |
| EPI_ISL_18472640 | A/hooded_crane/Kagoshima/KU-60/ | 2.3.4.4b | Asia / Japan | Avian | 2022/11/14 | A / H5N1 |
| EPI_ISL_18472641 | A/hooded_crane/Kagoshima/KU-61/ | 2.3.4.4b | Asia / Japan | Avian | 2022/11/14 | A / H5N1 |
| EPI_ISL_18472642 | A/white-naped_crane/Kagoshima/K | 2.3.4.4b | Asia / Japan | Avian | 2022/11/14 | A / H5N1 |
| EPI_ISL_18472643 | A/hooded_crane/Kagoshima/KU-63/ | 2.3.4.4b | Asia / Japan | Avian | 2022/11/14 | A / H5N1 |
| EPI_ISL_18472644 | A/white-naped_crane/Kagoshima/K | 2.3.4.4b | Asia / Japan | Avian | 2022/11/14 | A / H5N1 |
| EPI_ISL_18472645 | A/hooded_crane/Kagoshima/KU-67/ | 2.3.4.4b | Asia / Japan | Avian | 2022/11/15 | A / H5N1 |
| EPI_ISL_18472646 | A/hooded_crane/Kagoshima/KU-68/ | 2.3.4.4b | Asia / Japan | Avian | 2022/11/15 | A / H5N1 |
| EPI_ISL_18472647 | A/hooded_crane/Kagoshima/KU-69/ | 2.3.4.4b | Asia / Japan | Avian | 2022/11/15 | A / H5N1 |
| EPI_ISL_18472648 | A/hooded_crane/Kagoshima/KU-71/ | 2.3.4.4b | Asia / Japan | Avian | 2022/11/15 | A / H5N1 |

| | | | | | | |
|---|---|---|---|---|---|---|
| EPI_ISL_18472649 | A/hooded_crane/Kagoshima/KU-73/ | 2.3.4.4b | Asia / Japan | Avian | 2022/11/16 | A / H5N1 |
| EPI_ISL_18472650 | A/hooded_crane/Kagoshima/KU-74/ | 2.3.4.4b | Asia / Japan | Avian | 2022/11/16 | A / H5N1 |
| EPI_ISL_18472651 | A/hooded_crane/Kagoshima/KU-75/ | 2.3.4.4b | Asia / Japan | Avian | 2022/11/16 | A / H5N1 |
| EPI_ISL_18472652 | A/hooded_crane/Kagoshima/KU-76/ | 2.3.4.4b | Asia / Japan | Avian | 2022/11/16 | A / H5N1 |
| EPI_ISL_18472653 | A/hooded_crane/Kagoshima/KU-77/ | 2.3.4.4b | Asia / Japan | Avian | 2022/11/17 | A / H5N1 |
| EPI_ISL_18472654 | A/white-naped_crane/Kagoshima/K | 2.3.4.4b | Asia / Japan | Avian | 2022/11/17 | A / H5N1 |
| EPI_ISL_18472655 | A/hooded_crane/Kagoshima/KU-79/ | 2.3.4.4b | Asia / Japan | Avian | 2022/11/17 | A / H5N1 |
| EPI_ISL_18472656 | A/hooded_crane/Kagoshima/KU-80/ | 2.3.4.4b | Asia / Japan | Avian | 2022/11/17 | A / H5N1 |
| EPI_ISL_18472657 | A/white-naped_crane/Kagoshima/K | 2.3.4.4b | Asia / Japan | Avian | 2022/11/17 | A / H5N1 |
| EPI_ISL_18472658 | A/white-naped_crane/Kagoshima/K | 2.3.4.4b | Asia / Japan | Avian | 2022/11/18 | A / H5N1 |
| EPI_ISL_18472659 | A/white-naped_crane/Kagoshima/K | 2.3.4.4b | Asia / Japan | Avian | 2022/11/18 | A / H5N1 |
| EPI_ISL_18472660 | A/hooded_crane/Kagoshima/KU-89/ | 2.3.4.4b | Asia / Japan | Avian | 2022/11/19 | A / H5N1 |
| EPI_ISL_18472661 | A/white-naped_crane/Kagoshima/K | 2.3.4.4b | Asia / Japan | Avian | 2022/11/20 | A / H5N1 |
| EPI_ISL_18472662 | A/hooded_crane/Kagoshima/KU-94/ | 2.3.4.4b | Asia / Japan | Avian | 2022/11/21 | A / H5N1 |
| EPI_ISL_18472663 | A/white-naped_crane/Kagoshima/K | 2.3.4.4b | Asia / Japan | Avian | 2022/11/21 | A / H5N1 |
| EPI_ISL_18472664 | A/hooded_crane/Kagoshima/KU-96/ | 2.3.4.4b | Asia / Japan | Avian | 2022/11/21 | A / H5N1 |
| EPI_ISL_18472665 | A/white-naped_crane/Kagoshima/K | 2.3.4.4b | Asia / Japan | Avian | 2022/11/21 | A / H5N1 |
| EPI_ISL_18472666 | A/hooded_crane/Kagoshima/KU-10 | 2.3.4.4b | Asia / Japan | Avian | 2022/11/21 | A / H5N1 |
| EPI_ISL_18472667 | A/hooded_crane/Kagoshima/KU-10 | 2.3.4.4b | Asia / Japan | Avian | 2022/11/22 | A / H5N1 |
| EPI_ISL_18472668 | A/hooded_crane/Kagoshima/KU-10 | 2.3.4.4b | Asia / Japan | Avian | 2022/11/22 | A / H5N1 |
| EPI_ISL_18472669 | A/white-naped_crane/Kagoshima/K | 2.3.4.4b | Asia / Japan | Avian | 2022/11/22 | A / H5N1 |
| EPI_ISL_18472670 | A/hooded_crane/Kagoshima/KU-11 | 2.3.4.4b | Asia / Japan | Avian | 2022/11/22 | A / H5N1 |
| EPI_ISL_18472671 | A/white-naped_crane/Kagoshima/K | 2.3.4.4b | Asia / Japan | Avian | 2022/11/22 | A / H5N1 |
| EPI_ISL_18472672 | A/hooded_crane/Kagoshima/KU-11 | 2.3.4.4b | Asia / Japan | Avian | 2022/11/22 | A / H5N1 |
| EPI_ISL_18472673 | A/white-naped_crane/Kagoshima/K | 2.3.4.4b | Asia / Japan | Avian | 2022/11/23 | A / H5N1 |
| EPI_ISL_18472674 | A/hooded_crane/Kagoshima/KU-11 | 2.3.4.4b | Asia / Japan | Avian | 2022/11/23 | A / H5N1 |
| EPI_ISL_18472675 | A/hooded_crane/Kagoshima/KU-11 | 2.3.4.4b | Asia / Japan | Avian | 2022/11/23 | A / H5N1 |
| EPI_ISL_18472676 | A/hooded_crane/Kagoshima/KU-11 | 2.3.4.4b | Asia / Japan | Avian | 2022/11/23 | A / H5N1 |
| EPI_ISL_18472677 | A/hooded_crane/Kagoshima/KU-11 | 2.3.4.4b | Asia / Japan | Avian | 2022/11/24 | A / H5N1 |
| EPI_ISL_18472678 | A/hooded_crane/Kagoshima/KU-12 | 2.3.4.4b | Asia / Japan | Avian | 2022/11/24 | A / H5N1 |
| EPI_ISL_18472679 | A/hooded_crane/Kagoshima/KU-12 | 2.3.4.4b | Asia / Japan | Avian | 2022/11/24 | A / H5N1 |
| EPI_ISL_18472680 | A/hooded_crane/Kagoshima/KU-12 | 2.3.4.4b | Asia / Japan | Avian | 2022/11/24 | A / H5N1 |
| EPI_ISL_18472681 | A/hooded_crane/Kagoshima/KU-12 | 2.3.4.4b | Asia / Japan | Avian | 2022/11/25 | A / H5N1 |
| EPI_ISL_18472682 | A/white-naped_crane/Kagoshima/K | 2.3.4.4b | Asia / Japan | Avian | 2022/11/25 | A / H5N1 |
| EPI_ISL_18472683 | A/hooded_crane/Kagoshima/KU-12 | 2.3.4.4b | Asia / Japan | Avian | 2022/11/26 | A / H5N1 |
| EPI_ISL_18472684 | A/hooded_crane/Kagoshima/KU-12 | 2.3.4.4b | Asia / Japan | Avian | 2022/11/26 | A / H5N1 |
| EPI_ISL_18472685 | A/hooded_crane/Kagoshima/KU-13 | 2.3.4.4b | Asia / Japan | Avian | 2022/11/26 | A / H5N1 |
| EPI_ISL_18472686 | A/hooded_crane/Kagoshima/KU-13 | 2.3.4.4b | Asia / Japan | Avian | 2022/11/26 | A / H5N1 |
| EPI_ISL_18472687 | A/hooded_crane/Kagoshima/KU-13 | 2.3.4.4b | Asia / Japan | Avian | 2022/11/27 | A / H5N1 |
| EPI_ISL_18472688 | A/white-naped_crane/Kagoshima/K | 2.3.4.4b | Asia / Japan | Avian | 2022/11/27 | A / H5N1 |
| EPI_ISL_18472689 | A/hooded_crane/Kagoshima/KU-13 | 2.3.4.4b | Asia / Japan | Avian | 2022/11/27 | A / H5N1 |
| EPI_ISL_18472690 | A/hooded_crane/Kagoshima/KU-13 | 2.3.4.4b | Asia / Japan | Avian | 2022/11/27 | A / H5N1 |
| EPI_ISL_18472691 | A/hooded_crane/Kagoshima/KU-14 | 2.3.4.4b | Asia / Japan | Avian | 2022/11/28 | A / H5N1 |
| EPI_ISL_18472692 | A/white-naped_crane/Kagoshima/K | 2.3.4.4b | Asia / Japan | Avian | 2022/11/28 | A / H5N1 |
| EPI_ISL_18472693 | A/hooded_crane/Kagoshima/KU-14 | 2.3.4.4b | Asia / Japan | Avian | 2022/11/28 | A / H5N1 |
| EPI_ISL_18472694 | A/hooded_crane/Kagoshima/KU-14 | 2.3.4.4b | Asia / Japan | Avian | 2022/11/28 | A / H5N1 |
| EPI_ISL_18472695 | A/hooded_crane/Kagoshima/KU-14 | 2.3.4.4b | Asia / Japan | Avian | 2022/11/28 | A / H5N1 |
| EPI_ISL_18472696 | A/white-naped_crane/Kagoshima/K | 2.3.4.4b | Asia / Japan | Avian | 2022/11/29 | A / H5N1 |
| EPI_ISL_18472697 | A/hooded_crane/Kagoshima/KU-14 | 2.3.4.4b | Asia / Japan | Avian | 2022/11/29 | A / H5N1 |
| EPI_ISL_18472698 | A/hooded_crane/Kagoshima/KU-14 | 2.3.4.4b | Asia / Japan | Avian | 2022/11/29 | A / H5N1 |
| EPI_ISL_18472699 | A/hooded_crane/Kagoshima/KU-15 | 2.3.4.4b | Asia / Japan | Avian | 2022/11/30 | A / H5N1 |
| EPI_ISL_18472700 | A/hooded_crane/Kagoshima/KU-15 | 2.3.4.4b | Asia / Japan | Avian | 2022/11/30 | A / H5N1 |
| EPI_ISL_18472701 | A/white-naped_crane/Kagoshima/K | 2.3.4.4b | Asia / Japan | Avian | 2022/11/30 | A / H5N1 |
| EPI_ISL_18472702 | A/hooded_crane/Kagoshima/KU-16 | 2.3.4.4b | Asia / Japan | Avian | 2022/12/1 | A / H5N1 |
| EPI_ISL_18472703 | A/hooded_crane/Kagoshima/KU-16 | 2.3.4.4b | Asia / Japan | Avian | 2022/12/2 | A / H5N1 |
| EPI_ISL_18472704 | A/white-naped_crane/Kagoshima/K | 2.3.4.4b | Asia / Japan | Avian | 2022/12/1 | A / H5N1 |
| EPI_ISL_18472705 | A/hooded_crane/Kagoshima/KU-17 | 2.3.4.4b | Asia / Japan | Avian | 2022/12/3 | A / H5N1 |
| EPI_ISL_18472706 | A/white-naped_crane/Kagoshima/K | 2.3.4.4b | Asia / Japan | Avian | 2022/12/3 | A / H5N1 |
| EPI_ISL_18472707 | A/hooded_crane/Kagoshima/KU-17 | 2.3.4.4b | Asia / Japan | Avian | 2022/12/3 | A / H5N1 |
| EPI_ISL_18472708 | A/white-naped_crane/Kagoshima/K | 2.3.4.4b | Asia / Japan | Avian | 2022/12/4 | A / H5N1 |
| EPI_ISL_18472709 | A/white-naped_crane/Kagoshima/K | 2.3.4.4b | Asia / Japan | Avian | 2022/12/4 | A / H5N1 |
| EPI_ISL_18472710 | A/white-naped_crane/Kagoshima/K | 2.3.4.4b | Asia / Japan | Avian | 2022/12/5 | A / H5N1 |

| | | | | | | |
|---|---|---|---|---|---|---|
| EPI_ISL_18472711 | A/hooded_crane/Kagoshima/KU-18 | 2.3.4.4b | Asia / Japan | Avian | 2022/12/6 | A / H5N1 |
| EPI_ISL_18472712 | A/hooded_crane/Kagoshima/KU-18( | 2.3.4.4b | Asia / Japan | Avian | 2022/12/7 | A / H5N1 |
| EPI_ISL_18472713 | A/hooded_crane/Kagoshima/KU-18' | 2.3.4.4b | Asia / Japan | Avian | 2022/12/8 | A / H5N1 |
| EPI_ISL_18472714 | A/hooded_crane/Kagoshima/KU-19( | 2.3.4.4b | Asia / Japan | Avian | 2022/12/8 | A / H5N1 |
| EPI_ISL_18472715 | A/hooded_crane/Kagoshima/KU-19 | 2.3.4.4b | Asia / Japan | Avian | 2022/12/10 | A / H5N1 |
| EPI_ISL_18472716 | A/hooded_crane/Kagoshima/KU-19: | 2.3.4.4b | Asia / Japan | Avian | 2022/12/10 | A / H5N1 |
| EPI_ISL_18472717 | A/hooded_crane/Kagoshima/KU-20 | 2.3.4.4b | Asia / Japan | Avian | 2022/12/12 | A / H5N1 |
| EPI_ISL_18472718 | A/hooded_crane/Kagoshima/KU-21: | 2.3.4.4b | Asia / Japan | Avian | 2022/12/15 | A / H5N1 |
| EPI_ISL_18508592 | A/black_kite/Kagoshima/KU-140/2( | 2.3.4.4b | Asia / Japan / | Avian | 2022/11/28 | A / H5N1 |
| EPI_ISL_18612256 | A/Eurasian_wigeon/Kagoshima/KU- | 2.3.4.4b | Asia / Japan | Avian | 2023/11/15 | A / H5N1 |
| EPI_ISL_18612263 | A/Common_teal/Kagoshima/KU-6/2 | 2.3.4.4b | Asia / Japan | Avian | 2023/11/27 | A / H5N1 |
| EPI_ISL_18651568 | A/hooded_crane/Kagoshima/KU-12/ | 2.3.4.4b | Asia / Japan | Avian | 2023/12/11 | A / H5N1 |
| EPI_ISL_18770562 | A/white-naped_crane/Kagoshima/K | 2.3.4.4b | Asia / Japan | Avian | 2023/12/12 | A / H5N1 |
| EPI_ISL_18770563 | A/white-naped_crane/Kagoshima/K | 2.3.4.4b | Asia / Japan | Avian | 2023/12/15 | A / H5N1 |
| EPI_ISL_18770564 | A/hooded_crane/Kagoshima/KU-17/ | 2.3.4.4b | Asia / Japan | Avian | 2023/12/16 | A / H5N1 |
| EPI_ISL_18770565 | A/hooded_crane/Kagoshima/KU-21/ | 2.3.4.4b | Asia / Japan | Avian | 2023/12/21 | A / H5N1 |
| EPI_ISL_19624025 | A/Hooded_crane/Kagoshima/KU-24 | 2.3.4.4b | Asia / Japan | Avian | 2024/11/17 | A / H5N1 |
| EPI_ISL_19624026 | A/Hooded_crane/Kagoshima/KU-24 | 2.3.4.4b | Asia / Japan | Avian | 2024/11/16 | A / H5N1 |
| EPI_ISL_19645340 | A/Hooded_crane/Kagoshima/KU-24 | 2.3.4.4b | Asia / Japan | Avian | 2024/11/18 | A / H5N1 |
| EPI_ISL_19645341 | A/Eurasian_wigeon/Kagoshima/KU- | 2.3.4.4b | Asia / Japan | Avian | 2024/11/18 | A / H5N1 |
| EPI_ISL_19645342 | A/Hooded_crane/Kagoshima/KU-24 | 2.3.4.4b | Asia / Japan | Avian | 2024/11/21 | A / H5N1 |
| EPI_ISL_19645346 | A/Hooded_crane/Kagoshima/KU-24 | 2.3.4.4b | Asia / Japan | Avian | 2024/11/21 | A / H5N1 |
| EPI_ISL_19645347 | A/White-naped_crane/Kagoshima/K | 2.3.4.4b | Asia / Japan | Avian | 2024/11/21 | A / H5N1 |
| EPI_ISL_19645348 | A/Hooded_crane/Kagoshima/KU-24 | 2.3.4.4b | Asia / Japan | Avian | 2024/11/21 | A / H5N1 |
| EPI_ISL_19645349 | A/Hooded_crane/Kagoshima/KU-24 | 2.3.4.4b | Asia / Japan | Avian | 2024/11/22 | A / H5N1 |
| EPI_ISL_19645361 | A/Hooded_crane/Kagoshima/KU-24 | 2.3.4.4b | Asia / Japan | Avian | 2024/11/18 | A / H5N1 |
| EPI_ISL_19645362 | A/Hooded_crane/Kagoshima/KU-24 | 2.3.4.4b | Asia / Japan | Avian | 2024/11/18 | A / H5N1 |
| EPI_ISL_19645363 | A/Hooded_crane/Kagoshima/KU-24 | 2.3.4.4b | Asia / Japan | Avian | 2024/11/18 | A / H5N1 |
| EPI_ISL_19645364 | A/Hooded_crane/Kagoshima/KU-24 | 2.3.4.4b | Asia / Japan | Avian | 2024/11/19 | A / H5N1 |
| EPI_ISL_19700406 | A/Hooded_crane/Kagoshima/KU-24 | 2.3.4.4b | Asia / Japan | Avian | 2024/11/20 | A / H5N1 |
| EPI_ISL_19700407 | A/Hooded_crane/Kagoshima/KU-24 | 2.3.4.4b | Asia / Japan | Avian | 2024/11/20 | A / H5N1 |
| EPI_ISL_19700408 | A/Hooded_crane/Kagoshima/KU-24 | 2.3.4.4b | Asia / Japan | Avian | 2024/11/21 | A / H5N1 |
| EPI_ISL_19700409 | A/Hooded_crane/Kagoshima/KU-24 | 2.3.4.4b | Asia / Japan | Avian | 2024/11/21 | A / H5N1 |
| EPI_ISL_19700410 | A/Hooded_crane/Kagoshima/KU-24 | 2.3.4.4b | Asia / Japan | Avian | 2024/11/21 | A / H5N1 |
| EPI_ISL_19700683 | A/Hooded_crane/Kagoshima/KU-24 | 2.3.4.4b | Asia / Japan | Avian | 2024/11/19 | A / H5N1 |
| EPI_ISL_19700684 | A/Hooded_crane/Kagoshima/KU-24 | 2.3.4.4b | Asia / Japan | Avian | 2024/11/19 | A / H5N1 |
| EPI_ISL_19700685 | A/Hooded_crane/Kagoshima/KU-24 | 2.3.4.4b | Asia / Japan | Avian | 2024/11/22 | A / H5N1 |
| EPI_ISL_19700686 | A/Hooded_crane/Kagoshima/KU-24 | 2.3.4.4b | Asia / Japan | Avian | 2024/11/22 | A / H5N1 |
| EPI_ISL_19700687 | A/Hooded_crane/Kagoshima/KU-24 | 2.3.4.4b | Asia / Japan | Avian | 2024/11/22 | A / H5N1 |
| EPI_ISL_19700691 | A/Hooded_crane/Kagoshima/KU-24 | 2.3.4.4b | Asia / Japan | Avian | 2024/11/19 | A / H5N1 |
| EPI_ISL_19700692 | A/Hooded_crane/Kagoshima/KU-24 | 2.3.4.4b | Asia / Japan | Avian | 2024/11/23 | A / H5N1 |
| EPI_ISL_19700693 | A/Hooded_crane/Kagoshima/KU-24 | 2.3.4.4b | Asia / Japan | Avian | 2024/11/24 | A / H5N1 |
| EPI_ISL_19700694 | A/Hooded_crane/Kagoshima/KU-24 | 2.3.4.4b | Asia / Japan | Avian | 2024/11/25 | A / H5N1 |
| EPI_ISL_19700695 | A/Hooded_crane/Kagoshima/KU-24 | 2.3.4.4b | Asia / Japan | Avian | 2024/11/27 | A / H5N1 |
| EPI_ISL_19700696 | A/Whooper_swan/Hokkaido/0104P1 | 2.3.4.4b | Asia / Japan | Avian | 2024/10/25 | A / H5N1 |
| EPI_ISL_19700697 | A/Peregrine_Falcon/Hokkaido/2024 | 2.3.4.4b | Asia / Japan | Avian | 2024/10/31 | A / H5N1 |
| EPI_ISL_19707505 | A/Hooded_crane/Kagoshima/KU-24 | 2.3.4.4b | Asia / Japan | Avian | 2024/11/21 | A / H5N1 |
| EPI_ISL_19707506 | A/Hooded_crane/Kagoshima/KU-24 | 2.3.4.4b | Asia / Japan | Avian | 2024/11/25 | A / H5N1 |
| EPI_ISL_19707507 | A/Hooded_crane/Kagoshima/KU-24 | 2.3.4.4b | Asia / Japan | Avian | 2024/11/27 | A / H5N1 |
| EPI_ISL_19707508 | A/Hooded_crane/Kagoshima/KU-24 | 2.3.4.4b | Asia / Japan | Avian | 2024/11/28 | A / H5N1 |
| EPI_ISL_19707509 | A/Hooded_crane/Kagoshima/KU-24 | 2.3.4.4b | Asia / Japan | Avian | 2024/11/28 | A / H5N1 |
| EPI_ISL_19707510 | A/Hooded_crane/Kagoshima/KU-24 | 2.3.4.4b | Asia / Japan | Avian | 2024/11/28 | A / H5N1 |
| EPI_ISL_19707511 | A/Hooded_crane/Kagoshima/KU-24 | 2.3.4.4b | Asia / Japan | Avian | 2024/11/29 | A / H5N1 |
| EPI_ISL_19707512 | A/White-naped_crane/Kagoshima/K | 2.3.4.4b | Asia / Japan | Avian | 2024/11/29 | A / H5N1 |
| EPI_ISL_19004930 | A/Ruddy_Shelduck/Qinghai/07-HD | 2.3.4.4b | Asia / China | Avian | 2022/7/22 | A / H5N1 |
| EPI_ISL_19004931 | A/Crested_grebe/Qinghai/03-ZZ-S/2 | 2.3.4.4b | Asia / China | Avian | 2022/7/21 | A / H5N1 |
| EPI_ISL_19004933 | A/Brown-headed_gull/Qinghai/01-Z | 2.3.4.4b | Asia / China | Avian | 2022/7/21 | A / H5N1 |
| EPI_ISL_19004934 | A/Brown-headed_gull/Qinghai/02-Z | 2.3.4.4b | Asia / China | Avian | 2022/7/21 | A / H5N1 |
| EPI_ISL_19004935 | A/Ruddy_Shelduck/Qinghai/07-HD | 2.3.4.4b | Asia / China | Avian | 2022/7/22 | A / H5N1 |
| EPI_ISL_19004936 | A/Bar-headed_goose/Qinghai/07-JX | 2.3.4.4b | Asia / China | Avian | 2022/7/22 | A / H5N1 |
| EPI_ISL_19004937 | A/Bar-headed_goose/Qinghai/07-JX | 2.3.4.4b | Asia / China | Avian | 2022/7/22 | A / H5N1 |
| EPI_ISL_19004938 | A/Brown-headed_gull/Qinghai/06-Z | 2.3.4.4b | Asia / China | Avian | 2022/7/22 | A / H5N1 |

| | | | | | | |
|---|---|---|---|---|---|---|
| EPI_ISL_19004939 | A/Crested_grebe/Qinghai/04-ZZ-F/2 | 2.3.4.4b | Asia / China | Avian | 2022/7/15 | A / H5N1 |
| EPI_ISL_19004940 | A/Bar-headed_goose/Qinghai/07-JX | 2.3.4.4b | Asia / China | Avian | 2022/7/22 | A / H5N1 |
| EPI_ISL_19004941 | A/Brown-headed_gull/Qinghai/07-Z | 2.3.4.4b | Asia / China | Avian | 2022/7/20 | A / H5N1 |
| EPI_ISL_19004942 | A/Crested_grebe/Qinghai/10-ZZ-C/2 | 2.3.4.4b | Asia / China | Avian | 2022/7/15 | A / H5N1 |
| EPI_ISL_19004943 | A/Brown-headed_gull/Qinghai/11-Z | 2.3.4.4b | Asia / China | Avian | 2022/7/23 | A / H5N1 |
| EPI_ISL_19004944 | A/Crested_grebe/Qinghai/09-ZZ-F/2 | 2.3.4.4b | Asia / China | Avian | 2022/7/15 | A / H5N1 |
| EPI_ISL_19004945 | A/Crested_grebe/Qinghai/08-ZZ-F/2 | 2.3.4.4b | Asia / China | Avian | 2022/7/15 | A / H5N1 |
| EPI_ISL_19004946 | A/Brown-headed_gull/Qinghai/12-Z | 2.3.4.4b | Asia / China | Avian | 2022/7/23 | A / H5N1 |
| EPI_ISL_19004948 | A/Bar-headed_goose/Qinghai/06-22 | 2.3.4.4b | Asia / China | Avian | 2022/6/18 | A / H5N1 |
| EPI_ISL_19004949 | A/Bar-headed_goose/Qinghai/06-SH | 2.3.4.4b | Asia / China | Avian | 2022/6/18 | A / H5N1 |
| EPI_ISL_19004950 | A/Bar-headed_goose/Qinghai/06-SH | 2.3.4.4b | Asia / China | Avian | 2022/6/18 | A / H5N1 |
| EPI_ISL_19004951 | A/Bar-headed_goose/Qinghai/06-SH | 2.3.4.4b | Asia / China | Avian | 2022/6/18 | A / H5N1 |
| EPI_ISL_19004952 | A/Bar-headed_goose/Qinghai/06-SH | 2.3.4.4b | Asia / China | Avian | 2022/6/18 | A / H5N1 |
| EPI_ISL_19004953 | A/Bar-headed_goose/Qinghai/06-SH | 2.3.4.4b | Asia / China | Avian | 2022/6/18 | A / H5N1 |
| EPI_ISL_19004954 | A/Bar-headed_goose/Qinghai/06-SH | 2.3.4.4b | Asia / China | Avian | 2022/6/18 | A / H5N1 |
| EPI_ISL_14064697 | A/crow/Hokkaido/0101Q044/2022 | 2.3.4.4b | Asia / Japan / | Avian | 2022/1/20 | A / H5N1 |
| EPI_ISL_14064890 | A/crow/Hokkaido/0103B073/2022 | 2.3.4.4b | Asia / Japan / | Avian | 2022/4/1 | A / H5N1 |
| EPI_ISL_18529943 | A/europian_wigeon/Kagoshima/KU- | 2.3.4.4b | Asia / Japan | Avian | 2023/11/13 | A / H5N1 |
| EPI_ISL_18529944 | A/europian_wigeon/Kagoshima/KU- | 2.3.4.4b | Asia / Japan | Avian | 2023/11/20 | A / H5N1 |
| EPI_ISL_18529946 | A/northern_pintail/Kagoshima/KU-1 | 2.3.4.4b | Asia / Japan | Avian | 2023/11/13 | A / H5N1 |
| EPI_ISL_14644592 | A/African_barn_owl/South_Africa/2 | 2.3.4.4b | Africa / South | Avian | 2021/5/25 | A / H5N1 |
| EPI_ISL_14646080 | A/pelican/South_Africa/21050494/2 | 2.3.4.4b | Africa / South | Avian | 2021/5/29 | A / H5N1 |
| EPI_ISL_14918031 | A/Sacred_ibis/South_Africa/210603 | 2.3.4.4b | Africa / South | Avian | 2021/6/19 | A / H5N1 |
| EPI_ISL_14918839 | A/Blue_crane/South_Africa/210604 | 2.3.4.4b | Africa / South | Avian | 2021/6/28 | A / H5N1 |
| EPI_ISL_14918893 | A/Blue_crane/South_Africa/210700 | 2.3.4.4b | Africa / South | Avian | 2021/7/1 | A / H5N1 |
| EPI_ISL_14934885 | A/Cape_cormorant/South_Africa/21 | 2.3.4.4b | Africa / South | Avian | 2021/9/13 | A / H5N1 |
| EPI_ISL_19033339 | A/buzzard/Poland/MB098-N/2024 | 2.3.4.4b | Europe / Pola | Avian | 2024/2/27 | A / H5N1 |
| EPI_ISL_19033340 | A/buzzard/Poland/MB103-N/2024 | 2.3.4.4b | Europe / Pola | Avian | 2024/3/4 | A / H5N1 |
| EPI_ISL_198837 | A/chicken/Jilin/04.12_MHK003-O/2 | 2.3.2.1c | Asia / China | Chicken | 2015/4/12 | A / H5N1 |
| EPI_ISL_198840 | A/chicken/Jilin/04.25_MHK009/201 | 2.3.2.1c | Asia / China | Chicken | 2015/4/25 | A / H5N1 |
| EPI_ISL_198845 | A/chicken/Jilin/04.12_MHK001-O/2 | 2.3.2.1c | Asia / China | Chicken | 2015/4/12 | A / H5N1 |
| EPI_ISL_198846 | A/chicken/Jilin/04.12_MHK002-O/2 | 2.3.2.1c | Asia / China | Chicken | 2015/4/12 | A / H5N1 |
| EPI_ISL_199154 | A/chicken/Shandong/12.03_YA-CK | 2.3.2.1c | Asia / China | Chicken | 2014/12/3 | A / H5N1 |
| EPI_ISL_199164 | A/chicken/Jiangsu/03.17_WX006-P/ | 2.3.2.1c | Asia / China | Chicken | 2015/3/17 | A / H5N1 |
| EPI_ISL_199261 | A/chicken/Jiangsu/12.22_TCCX035 | 2.3.2.1c | Asia / China | Chicken | 2014/12/22 | A / H5N1 |
| EPI_ISL_199390 | A/chicken/Jiangsu/12.22_TCCX003 | 2.3.2.1c | Asia / China | Chicken | 2014/12/22 | A / H5N1 |
| EPI_ISL_12572652 | A/chicken/Anhui/S1740/2022 | 2.3.4.4b | Asia / China / | Chicken | 2022/3/3 | A / H5N1 |
| EPI_ISL_12572653 | A/chicken/Jiangxi/S40653/2021 | 2.3.4.4b | Asia / China / | Chicken | 2021/12/7 | A / H5N1 |
| EPI_ISL_19033337 | A/chicken/Poland/H79-T2/2024 | 2.3.4.4b | Europe / Pola | Chicken | 2024/2/26 | A / H5N1 |
| EPI_ISL_19033549 | A/chicken/Poland/H45-NM/2024 | 2.3.4.4b | Europe / Pola | Chicken | 2024/2/12 | A / H5N1 |
| EPI_ISL_19602897 | A/laying_hen/Poland/H411NM/2024 | 2.3.4.4b | Europe / Pola | Chicken | 2024/10/18 | A / H5N1 |
| EPI_ISL_19602961 | A/laying_hen/Poland/H438NM/2024 | 2.3.4.4b | Europe / Pola | Chicken | 2024/10/26 | A / H5N1 |
| EPI_ISL_19602964 | A/laying_hen/Poland/H449NM/2024 | 2.3.4.4b | Europe / Pola | Chicken | 2024/10/28 | A / H5N1 |
| EPI_ISL_19602965 | A/laying_hen/Poland/H467NM/2024 | 2.3.4.4b | Europe / Pola | Chicken | 2024/11/3 | A / H5N1 |
| EPI_ISL_19602985 | A/chicken/Poland/H464N/2024 | 2.3.4.4b | Europe / Pola | Chicken | 2024/10/31 | A / H5N1 |
| EPI_ISL_244144 | A/chicken/Vietnam/HU3-67/2015 | 2.3.2.1c | Asia / Vietnan | Gallus ga | 2015/8/15 | A / H5N1 |
| EPI_ISL_244145 | A/chicken/Vietnam/HU3-83/2015 | 2.3.2.1c | Asia / Vietnan | Gallus ga | 2015/8/15 | A / H5N1 |
| EPI_ISL_15844573 | A/Broiler_Chicken/BC/FAV-0228/2 | 2.3.4.4b | North Americ | Gallus ga | 2022/4/12 | A / H5N1 |
| EPI_ISL_14620233 | A/Buff_Orpington_chicken/South_A | 2.3.4.4b | Africa / South | Gallus ga | 2021/5/19 | A / H5N1 |
| EPI_ISL_198764 | A/duck/Hunan/12.07_YYGK100-P/ | 2.3.2.1c | Asia / China | Duck | 2013/12/7 | A / H5N1 |
| EPI_ISL_198774 | A/duck/Hunan/12.07_YYGK007-P/ | 2.3.2.1c | Asia / China | Duck | 2013/12/7 | A / H5N1 |
| EPI_ISL_198776 | A/duck/Hunan/12.07__YYGK97-P/ | 2.3.2.1c | Asia / China | Duck | 2013/12/7 | A / H5N1 |
| EPI_ISL_199019 | A/duck/Hubei/03.06__WHWTZ015 | 2.3.2.1c | Asia / China | Duck | 2015/3/6 | A / H5N1 |
| EPI_ISL_199239 | A/duck/Jiangsu/12.18_NJLH1270-P | 2.3.2.1c | Asia / China | Duck | 2014/12/18 | A / H5N1 |
| EPI_ISL_18819797 | A/duck/Korea/D502/2023 | 2.3.4.4b | Asia / Korea, | Duck | 2023/12/20 | A / H5N1 |
| EPI_ISL_18819960 | A/duck/Korea/D448-N1/2023 | 2.3.4.4b | Asia / Korea, | Duck | 2023/12/3 | A / H5N1 |
| EPI_ISL_18508589 | A/northern_pintail/Kagoshima/KU-6 | 2.3.4.4b | Asia / Japan / | Duck | 2022/11/14 | A / H5N1 |
| EPI_ISL_14573124 | A/Egyptian_goose/South_Africa/21( | 2.3.4.4b | Africa / South | Duck | 2021/5/13 | A / H5N1 |
| EPI_ISL_14710708 | A/duck/South_Africa/21060064/202 | 2.3.4.4b | Africa / South | Duck | 2021/6/3 | A / H5N1 |
| EPI_ISL_12572654 | A/duck/Guangdong/S4518/2021 | 2.3.4.4b | Asia / China / | Duck | 2021/12/8 | A / H5N1 |
| EPI_ISL_12572655 | A/duck/Guangdong/S4525/2021 | 2.3.4.4b | Asia / China / | Duck | 2021/12/8 | A / H5N1 |
| EPI_ISL_12572656 | A/duck/Guizhou/S1321/2022 | 2.3.4.4b | Asia / China / | Duck | 2022/2/22 | A / H5N1 |

| | | | | | | |
|---|---|---|---|---|---|---|
| EPI_ISL_12572657 | A/duck/Hubei/S4465/2021 | 2.3.4.4b | Asia / China / Duck | | 2021/11/27 | A / H5N1 |
| EPI_ISL_12572658 | A/duck/Hubei/SE128/2022 | 2.3.4.4b | Asia / China / Duck | | 2022/1/10 | A / H5N1 |
| EPI_ISL_12572659 | A/duck/Hubei/SE220/2022 | 2.3.4.4b | Asia / China / Duck | | 2022/1/10 | A / H5N1 |
| EPI_ISL_12572660 | A/duck/Jiangxi/S40833/2021 | 2.3.4.4b | Asia / China / Duck | | 2021/12/8 | A / H5N1 |
| EPI_ISL_19033331 | A/domestic_duck/Poland/H60-T4/2( | 2.3.4.4b | Europe / Pola | Duck | 2024/2/15 | A / H5N1 |
| EPI_ISL_19033333 | A/domestic_duck/Poland/H69-T2/2( | 2.3.4.4b | Europe / Pola | Duck | 2024/2/23 | A / H5N1 |
| EPI_ISL_19033554 | A/domestic_duck/Poland/H52-T1K2 | 2.3.4.4b | Europe / Pola | Duck | 2024/2/16 | A / H5N1 |
| EPI_ISL_19033555 | A/domestic_duck/Poland/H57-T1/2( | 2.3.4.4b | Europe / Pola | Duck | 2024/2/15 | A / H5N1 |
| EPI_ISL_19033558 | A/domestic_duck/Poland/H66-T1/2( | 2.3.4.4b | Europe / Pola | Duck | 2024/2/20 | A / H5N1 |
| EPI_ISL_19602772 | A/domestic_duck/Poland/H361NM/ | 2.3.4.4b | Europe / Pola | Duck | 2024/10/5 | A / H5N1 |
| EPI_ISL_19602808 | A/domestic_duck/Poland/H372NM/ | 2.3.4.4b | Europe / Pola | Duck | 2024/10/10 | A / H5N1 |
| EPI_ISL_19602896 | A/domestic_duck/Poland/H391NM/ | 2.3.4.4b | Europe / Pola | Duck | 2024/10/12 | A / H5N1 |
| EPI_ISL_19602957 | A/domestic_duck/Poland/H433T1/2 | 2.3.4.4b | Europe / Pola | Duck | 2024/10/24 | A / H5N1 |
| EPI_ISL_19602962 | A/domestic_duck/Poland/H440NM/ | 2.3.4.4b | Europe / Pola | Duck | 2024/10/28 | A / H5N1 |
| EPI_ISL_244143 | A/duck/Vietnam/HU3-16/2015 | 2.3.2.1c | Asia / Vietnar | Anas plat | 2015/8/15 | A / H5N1 |
| EPI_ISL_400767 | A/duck/East_Java/Av378/2014/H5N | 2.3.2.1c | Asia / Indone: | Anas sp. | 2014/12/21 | A / H5N1 |
| EPI_ISL_400769 | A/duck/East_Java/Av731/2016/H5N | 2.3.2.1c | Asia / Indone: | Anas sp. | 2016/1/22 | A / H5N1 |
| EPI_ISL_400771 | A/duck/East_Java/Av682/2016/H5N | 2.3.2.1c | Asia / Indone: | Anas sp. | 2016/1/15 | A / H5N1 |
| EPI_ISL_14710814 | A/African_fish_eagle/South_Africa/ | 2.3.4.4b | Africa / South | Eagle | 2021/6/3 | A / H5N1 |
| EPI_ISL_14638909 | A/African_penguin/South_Africa/21 | 2.3.4.4b | Africa / South | Penguin | 2021/5/20 | A / H5N1 |
| EPI_ISL_19496405 | A/large-billed_crow/Hokkaido/0111 | 2.3.4.4b | Asia / Japan / | Wild bird | 2023/11/28 | A / H5N1 |
| EPI ISL 18717640 | A/Eurasian wigeon/korea/23WS022 | 2.3.4.4b | Asia / Korea, | Mareca p | 2023/11/27 | A / H5N1 |
| EPI_ISL_19603112 | A/greylag_goose/Poland/MB408NM | 2.3.4.4b | Europe / Pola | Greylag g | 2024/10/22 | A / H5N1 |
| EPI_ISL_19603554 | A/greylag_goose/Poland/MB441NM | 2.3.4.4b | Europe / Pola | Greylag g | 2024/11/4 | A / H5N1 |
| EPI_ISL_14973759 | A/Cape_cormorant/South_Africa/21 | 2.3.4.4b | Africa / South | Cormorar | 2021/10/7 | A / H5N1 |
| EPI_ISL_14973864 | A/Cape_cormorant/South_Africa/21 | 2.3.4.4b | Africa / South | Cormorar | 2021/10/7 | A / H5N1 |
| EPI_ISL_244141 | A/Muscovy_duck/Vietnam/HU3-97/ | 2.3.2.1c | Asia / Vietnar | Cairina m | 2015/8/15 | A / H5N1 |
| EPI_ISL_244142 | A/Muscovy_duck/Vietnam/HU3-35 | 2.3.2.1c | Asia / Vietnar | Cairina m | 2015/8/15 | A / H5N1 |
| EPI_ISL_15843373 | A/Bald_Eagle/BC/OTH-33-36/2022 | 2.3.4.4b | North Americ | Haliaeetu | 2022/2/3 | A / H5N1 |
| EPI_ISL_14573033 | A/chicken/South_Africa/690575/202 | 2.3.4.4b | Africa / South | Gallus ga | 2021/5/10 | A / H5N1 |
| EPI_ISL_14542488 | A/chicken/South_Africa/26683/2021 | 2.3.4.4b | Africa / South | Gallus ga | 2021/4/9 | A / H5N1 |
| EPI_ISL_14542530 | A/chicken/South_Africa/26700/2021 | 2.3.4.4b | Africa / South | Gallus ga | 2021/4/19 | A / H5N1 |
| EPI_ISL_14573038 | A/chicken/South_Africa/690813/202 | 2.3.4.4b | Africa / South | Gallus ga | 2021/5/10 | A / H5N1 |
| EPI_ISL_14620174 | A/chicken/South_Africa/692329/202 | 2.3.4.4b | Africa / South | Gallus ga | 2021/5/18 | A / H5N1 |
| EPI_ISL_14643779 | A/chicken/South_Africa/692881/202 | 2.3.4.4b | Africa / South | Gallus ga | 2021/5/20 | A / H5N1 |
| EPI_ISL_14644201 | A/chicken/South_Africa/693331/202 | 2.3.4.4b | Africa / South | Gallus ga | 2021/5/23 | A / H5N1 |
| EPI_ISL_14645717 | A/chicken/South_Africa/693781/202 | 2.3.4.4b | Africa / South | Gallus ga | 2021/5/27 | A / H5N1 |
| EPI_ISL_14710239 | A/chicken/South_Africa/693965/202 | 2.3.4.4b | Africa / South | Gallus ga | 2021/5/31 | A / H5N1 |
| EPI_ISL_14918217 | A/chicken/South_Africa/697352/202 | 2.3.4.4b | Africa / South | Gallus ga | 2021/6/20 | A / H5N1 |
| EPI_ISL_14918892 | A/chicken/South_Africa/683320/202 | 2.3.4.4b | Africa / South | Gallus ga | 2021/6/30 | A / H5N1 |
| EPI_ISL_14918903 | A/chicken/South_Africa/411255/202 | 2.3.4.4b | Africa / South | Gallus ga | 2021/7/19 | A / H5N1 |
| EPI_ISL_14932914 | A/chicken/South_Africa/411542/202 | 2.3.4.4b | Africa / South | Gallus ga | 2021/7/20 | A / H5N1 |
| EPI_ISL_14933095 | A/chicken/South_Africa/411554/202 | 2.3.4.4b | Africa / South | Gallus ga | 2021/7/23 | A / H5N1 |
| EPI_ISL_14933096 | A/chicken/South_Africa/33071/2021 | 2.3.4.4b | Africa / South | Gallus ga | 2021/7/25 | A / H5N1 |
| EPI_ISL_14933117 | A/chicken/South_Africa/33069/2021 | 2.3.4.4b | Africa / South | Gallus ga | 2021/7/26 | A / H5N1 |
| EPI_ISL_14933695 | A/chicken/South_Africa/412364/202 | 2.3.4.4b | Africa / South | Gallus ga | 2021/7/26 | A / H5N1 |
| EPI_ISL_14933721 | A/chicken/South_Africa/33081/2021 | 2.3.4.4b | Africa / South | Gallus ga | 2021/7/27 | A / H5N1 |
| EPI_ISL_14933723 | A/chicken/South_Africa/412372/202 | 2.3.4.4b | Africa / South | Gallus ga | 2021/7/28 | A / H5N1 |
| EPI_ISL_14933725 | A/chicken/South_Africa/412369/202 | 2.3.4.4b | Africa / South | Gallus ga | 2021/7/28 | A / H5N1 |
| EPI_ISL_14933916 | A/chicken/South_Africa/411258/202 | 2.3.4.4b | Africa / South | Gallus ga | 2021/7/29 | A / H5N1 |
| EPI_ISL_14934317 | A/chicken/South_Africa/412374/202 | 2.3.4.4b | Africa / South | Gallus ga | 2021/7/30 | A / H5N1 |
| EPI_ISL_14934318 | A/chicken/South_Africa/411262/202 | 2.3.4.4b | Africa / South | Gallus ga | 2021/8/9 | A / H5N1 |
| EPI_ISL_14934383 | A/chicken/South_Africa/10766/2021 | 2.3.4.4b | Africa / South | Gallus ga | 2021/8/16 | A / H5N1 |
| EPI_ISL_14934479 | A/chicken/South_Africa/424469/202 | 2.3.4.4b | Africa / South | Gallus ga | 2021/8/18 | A / H5N1 |
| EPI_ISL_14934883 | A/chicken/South_Africa/412380/202 | 2.3.4.4b | Africa / South | Gallus ga | 2021/9/7 | A / H5N1 |
| EPI_ISL_14934884 | A/chicken/South_Africa/412381/202 | 2.3.4.4b | Africa / South | Gallus ga | 2021/9/8 | A / H5N1 |
| EPI ISL 14542831 | A/chicken/South Africa/UP01/2021 | 2.3.4.4b | Africa / South | Gallus ga | 2021/5/1 | A / H5N1 |
| EPI_ISL_14573053 | A/chicken/South_Africa/UP481/202 | 2.3.4.4b | Africa / South | Gallus ga | 2021/5/10 | A / H5N1 |
| EPI_ISL_14934607 | A/chicken/South_Africa/MAB/2021 | 2.3.4.4b | Africa / South | Gallus ga | 2021/8/27 | A / H5N1 |
| EPI_ISL_14543004 | A/chicken/South_Africa/21050090/2 | 2.3.4.4b | Africa / South | Gallus ga | 2021/5/6 | A / H5N1 |
| EPI_ISL_14543005 | A/chicken/South_Africa/21050118/2 | 2.3.4.4b | Africa / South | Gallus ga | 2021/5/7 | A / H5N1 |
| EPI_ISL_14571452 | A/chicken/South_Africa/21050119/2 | 2.3.4.4b | Africa / South | Gallus ga | 2021/5/7 | A / H5N1 |

| | | | | | | |
|---|---|---|---|---|---|---|
| EPI_ISL_14572124 | A/chicken/South_Africa/21050125/2 | 2.3.4.4b | Africa / South | Gallus ga | 2021/5/8 | A / H5N1 |
| EPI_ISL_14572927 | A/chicken/South_Africa/21050021/2 | 2.3.4.4b | Africa / South | Gallus ga | 2021/5/10 | A / H5N1 |
| EPI_ISL_14619020 | A/chicken/South_Africa/21050293/2 | 2.3.4.4b | Africa / South | Gallus ga | 2021/5/17 | A / H5N1 |
| EPI_ISL_14619024 | A/chicken/South_Africa/21050299/2 | 2.3.4.4b | Africa / South | Gallus ga | 2021/5/18 | A / H5N1 |
| EPI_ISL_14711165 | A/chicken/South_Africa/21060265/2 | 2.3.4.4b | Africa / South | Gallus ga | 2021/6/14 | A / H5N1 |
| EPI_ISL_14918605 | A/chicken/South_Africa/21060435/2 | 2.3.4.4b | Africa / South | Gallus ga | 2021/6/24 | A / H5N1 |
| EPI_ISL_14918800 | A/chicken/South_Africa/21060469/2 | 2.3.4.4b | Africa / South | Gallus ga | 2021/6/25 | A / H5N1 |
| EPI_ISL_12572663 | A/pigeon/Jiangxi/S40784/2021 | 2.3.4.4b | Asia / China / | Pigeon | 2021/12/8 | A / H5N1 |
| EPI_ISL_19603113 | A/bean_goose/Poland/MB416NJ/20 | 2.3.4.4b | Europe / Pola | Bean goo | 2024/10/18 | A / H5N1 |
| EPI_ISL_3135897 | A/Turkey/Hungary/16603/2021 | 2.3.4.4b | Europe / Hung | Meleagris | 2021/4/13 | A / H5N1 |
| EPI_ISL_17971989 | A/domestic_cat/Poland/H255-M/202 | 2.3.4.4b | Europe / Pola | Felis catu | 2023/6/21 | A / H5N1 |
| EPI_ISL_17971990 | A/domestic_cat/Poland/H256-G/202 | 2.3.4.4b | Europe / Pola | Felis catu | 2023/6/24 | A / H5N1 |
| EPI_ISL_17971991 | A/domestic_cat/Poland/H257-G/202 | 2.3.4.4b | Europe / Pola | Felis catu | 2023/6/24 | A / H5N1 |
| EPI_ISL_17971992 | A/domestic_cat/Poland/H263-G/202 | 2.3.4.4b | Europe / Pola | Felis catu | 2023/6/26 | A / H5N1 |
| EPI_ISL_17971993 | A/domestic_cat/Poland/H266-W/202 | 2.3.4.4b | Europe / Pola | Felis catu | 2023/6/19 | A / H5N1 |
| EPI_ISL_17971994 | A/domestic_cat/Poland/H267-W/202 | 2.3.4.4b | Europe / Pola | Felis catu | 2023/6/25 | A / H5N1 |
| EPI_ISL_17971995 | A/domestic_cat/Poland/H270-W/202 | 2.3.4.4b | Europe / Pola | Felis catu | 2023/6/25 | A / H5N1 |
| EPI_ISL_17971996 | A/domestic_cat/Poland/H271-W/202 | 2.3.4.4b | Europe / Pola | Felis catu | 2023/6/25 | A / H5N1 |
| EPI_ISL_17971997 | A/domestic_cat/Poland/H277-W1/20 | 2.3.4.4b | Europe / Pola | Felis catu | 2023/6/26 | A / H5N1 |
| EPI_ISL_17971998 | A/domestic_cat/Poland/H264-G/202 | 2.3.4.4b | Europe / Pola | Felis catu | 2023/6/26 | A / H5N1 |
