## Appendix Files for "Transcontinental Spread of HPAI H5N1 from South America to Antarctica via Avian Vectors": AppendixTable3.pdf

**Appendix Table S3.** Detailed metadata of the reference sequences obtained from NCBI, which were used for multiple sequence alignment and phylogenetic analysis of the sequences generated in this study.

| Strain | Host_Name | Host_Group | Collection_I | Isolation_ | Geographic_Group |
|---|---|---|---|---|---|
| A/America | American black duck | Avian | 2022/2/6 | USA | North America |
| A/America | American black duck | Avian | 2022/2/16 | USA | North America |
| A/America | American black duck | Avian | 2022/2/16 | USA | North America |
| A/America | American black duck | Avian | 2022/2/16 | USA | North America |
| A/America | American black duck | Avian | 2022/3/4 | USA | North America |
| A/America | American black duck | Avian | 2022/3/4 | USA | North America |
| A/America | American black duck | Avian | 2022/3/9 | USA | North America |
| A/America | American crow | Avian | 2022/3/29 | USA | North America |
| A/America | American crow | Avian | 2022/4/18 | USA | North America |
| A/America | American crow | Avian | 2022/4/25 | USA | North America |
| A/America | American crow | Avian | 2022/4/20 | USA | North America |
| A/America | American crow | Avian | 2022/4/1 | USA | North America |
| A/America | American crow | Avian | 2022/4/7 | USA | North America |
| A/America | American crow | Avian | 2022/4/7 | USA | North America |
| A/America | American crow | Avian | 2022/4/7 | USA | North America |
| A/America | American crow | Avian | 2022/4/7 | USA | North America |
| A/America | American crow | Avian | 2022/4/21 | USA | North America |
| A/America | American crow | Avian | 2022/4/21 | USA | North America |
| A/America | American crow | Avian | 2022/4/21 | USA | North America |
| A/America | American crow | Avian | 2022/4/21 | USA | North America |
| A/America | American crow | Avian | 2022/4/21 | USA | North America |
| A/America | American crow | Avian | 2022/4/21 | USA | North America |
| A/America | American green-winge | Avian | 2022/2/12 | USA | North America |
| A/America | Anas carolinensis | Avian | 2022/9/21 | USA | North America |
| A/America | Pelecanus erythrorhync | Avian | 2022/3/10 | USA | North America |
| A/America | American pelican | | 2022/4/24 | USA | North America |
| A/America | American white pelica | Avian | 2022/4/12 | USA | North America |
| A/America | American white pelica | Avian | 2022/4/12 | USA | North America |
| A/America | American white pelica | Avian | 2022/4/12 | USA | North America |
| A/America | American white pelica | Avian | 2022/2/28 | USA | North America |
| A/America | American white pelica | Avian | 2022/3/29 | USA | North America |
| A/America | American wigeon | Avian | 2022/2/5 | USA | North America |
| A/America | American wigeon | Avian | 2022/1/26 | USA | North America |
| A/America | American wigeon | Avian | 2022/3/23 | USA | North America |
| A/America | American wigeon | Avian | 2022/1/8 | USA | North America |
| A/America | American wigeon | Avian | 2022/1/8 | USA | North America |
| A/America | American wigeon | Avian | 2022/1/8 | USA | North America |
| A/America | American wigeon | Avian | 2022/1/8 | USA | North America |
| A/America | American wigeon | Avian | 2022/1/8 | USA | North America |
| A/America | American wigeon | Avian | 2022/1/8 | USA | North America |
| A/America | American wigeon | Avian | 2022/1/8 | USA | North America |
| A/America | American wigeon | Avian | 2022/1/8 | USA | North America |
| A/America | American wigeon | Avian | 2022/1/8 | USA | North America |
| A/America | American wigeon | Avian | 2022/1/8 | USA | North America |
| A/America | American wigeon | Avian | 2022/1/8 | USA | North America |
| A/America | American wigeon | Avian | 2022/1/8 | USA | North America |
| A/America | American wigeon | Avian | 2022/1/8 | USA | North America |
| A/America | American wigeon | Avian | 2022/1/8 | USA | North America |
| A/America | American wigeon | Avian | 2022/1/8 | USA | North America |
| A/America | American wigeon | Avian | 2022/1/8 | USA | North America |
| A/America | American wigeon | Avian | 2022/1/8 | USA | North America |
| A/America | American wigeon | Avian | 2022/1/8 | USA | North America |
| A/America | American wigeon | Avian | 2022/1/8 | USA | North America |
| A/America | American wigeon | Avian | 2022/1/8 | USA | North America |

| | | | | | |
|---|---|---|---|---|---|
| A/America | American wigeon | Avian | 2022/1/22 | USA | North America |
| A/America | American wigeon | Avian | 2022/1/26 | USA | North America |
| A/America | American wigeon | Avian | 2021/12/30 | USA | North America |
| A/America | American wigeon | Avian | 2022/1/6 | USA | North America |
| A/America | American wigeon | Avian | 2022/1/6 | USA | North America |
| A/America | American wigeon | Avian | 2021/12/30 | USA | North America |
| A/Backyard | Backyard bird | | 2022/4/28 | USA | North America |
| A/Backyard | Backyard bird | | 2022/4/28 | USA | North America |
| A/Backyard | Backyard bird | | 2022/4/7 | USA | North America |
| A/Backyard | Backyard bird | | 2022/4/15 | USA | North America |
| A/Backyard | Backyard bird | | 2022/4/15 | USA | North America |
| A/Backyard | Backyard bird | | 2022/2/28 | USA | North America |
| A/Backyard | Backyard bird | | 2022/4/18 | USA | North America |
| A/Backyard | Backyard bird | | 2022/4/18 | USA | North America |
| A/Backyard | Backyard bird | | 2022/4/18 | USA | North America |
| A/Backyard | Backyard bird | | 2022/4/18 | USA | North America |
| A/Backyard | Backyard bird | | 2022/3/29 | USA | North America |
| A/Backyard | Backyard bird | | 2022/3/29 | USA | North America |
| A/Backyard | Backyard bird | | 2022/2/28 | USA | North America |
| A/Backyard | Backyard bird | | 2022/3/19 | USA | North America |
| A/Backyard | Backyard bird | | 2022/4/25 | USA | North America |
| A/Backyard | Backyard bird | | 2022/4/25 | USA | North America |
| A/Backyard | Backyard bird | | 2022/3/16 | USA | North America |
| A/Backyard | Backyard bird | | 2022/3/16 | USA | North America |
| A/Backyard | Backyard bird | | 2022/3/16 | USA | North America |
| A/Backyard | Backyard bird | | 2022/3/16 | USA | North America |
| A/Backyard | Backyard bird | | 2022/3/24 | USA | North America |
| A/Backyard | Backyard bird | | 2022/4/26 | USA | North America |
| A/Backyard | Backyard bird | | 2022/2/20 | USA | North America |
| A/Backyard | Backyard bird | | 2022/3/10 | USA | North America |
| A/Backyard | Backyard bird | | 2022/3/10 | USA | North America |
| A/Backyard | Backyard bird | | 2022/3/18 | USA | North America |
| A/Backyard | Backyard bird | | 2022/3/18 | USA | North America |
| A/Backyard | Backyard bird | | 2022/3/25 | USA | North America |
| A/Backyard | Backyard bird | | 2022/3/25 | USA | North America |
| A/Backyard | Backyard bird | | 2022/3/25 | USA | North America |
| A/Backyard | Backyard bird | | 2022/2/22 | USA | North America |
| A/Backyard | Backyard bird | | 2022/2/22 | USA | North America |
| A/Backyard | Backyard bird | | 2022/3/22 | USA | North America |
| A/Backyard | Backyard bird | | 2022/3/22 | USA | North America |
| A/Backyard | Backyard bird | | 2022/3/22 | USA | North America |
| A/Backyard | Backyard bird | | 2022/4/7 | USA | North America |
| A/Backyard | Backyard bird | | 2022/4/7 | USA | North America |
| A/Backyard | Backyard bird | | 2022/4/11 | USA | North America |
| A/Backyard | Backyard bird | | 2022/4/11 | USA | North America |
| A/Backyard | Backyard bird | | 2022/4/13 | USA | North America |
| A/Backyard | Backyard bird | | 2022/4/13 | USA | North America |
| A/Backyard | Backyard bird | | 2022/4/13 | USA | North America |
| A/Backyard | Backyard bird | | 2022/4/13 | USA | North America |
| A/Backyard | Backyard bird | | 2022/4/14 | USA | North America |
| A/Backyard | Backyard bird | | 2022/4/14 | USA | North America |
| A/Backyard | Backyard bird | | 2022/4/14 | USA | North America |
| A/Backyard | Backyard bird | | 2022/4/22 | USA | North America |
| A/Backyard | Backyard bird | | 2022/4/22 | USA | North America |
| A/Backyard | Backyard bird | | 2022/3/25 | USA | North America |
| A/Backyard | Backyard bird | | 2022/3/25 | USA | North America |
| A/Backyard | Backyard bird | | 2022/3/29 | USA | North America |
| A/Backyard | Backyard bird | | 2022/3/29 | USA | North America |

| | | | | |
|---|---|---|---|---|
| A/Backyard | Backyard bird | 2022/4/11 | USA | North America |
| A/Backyard | Backyard bird | 2022/4/11 | USA | North America |
| A/Backyard | Backyard bird | 2022/4/11 | USA | North America |
| A/Backyard | Backyard bird | 2022/4/28 | USA | North America |
| A/Backyard | Backyard bird | 2022/4/28 | USA | North America |
| A/Backyard | Backyard bird | 2022/4/28 | USA | North America |
| A/Backyard | Backyard bird | 2022/4/28 | USA | North America |
| A/Backyard | Backyard bird | 2022/4/28 | USA | North America |
| A/Backyard | Backyard bird | 2022/4/28 | USA | North America |
| A/Backyard | Backyard bird | 2022/4/28 | USA | North America |
| A/Backyard | Backyard bird | 2022/4/29 | USA | North America |
| A/Backyard | Backyard bird | 2022/4/29 | USA | North America |
| A/Backyard | Backyard bird | 2022/3/3 | USA | North America |
| A/Backyard | Backyard bird | 2022/3/14 | USA | North America |
| A/Backyard | Backyard bird | 2022/3/25 | USA | North America |
| A/Backyard | Backyard bird | 2022/4/5 | USA | North America |
| A/Backyard | Backyard bird | 2022/4/5 | USA | North America |
| A/Backyard | Backyard bird | 2022/3/14 | USA | North America |
| A/Backyard | Backyard bird | 2022/3/14 | USA | North America |
| A/Backyard | Backyard bird | 2022/3/24 | USA | North America |
| A/Backyard | Backyard bird | 2022/3/24 | USA | North America |
| A/Backyard | Backyard bird | 2022/3/31 | USA | North America |
| A/Backyard | Backyard bird | 2022/4/26 | USA | North America |
| A/Backyard | Backyard bird | 2022/2/22 | USA | North America |
| A/Backyard | Backyard bird | 2022/2/22 | USA | North America |
| A/Backyard | Backyard bird | 2022/3/28 | USA | North America |
| A/Backyard | Backyard bird | 2022/3/28 | USA | North America |
| A/Backyard | Backyard bird | 2022/4/4 | USA | North America |
| A/Backyard | Backyard bird | 2022/4/4 | USA | North America |
| A/Backyard | Backyard bird | 2022/4/4 | USA | North America |
| A/Backyard | Backyard bird | 2022/4/4 | USA | North America |
| A/Backyard | Backyard bird | 2022/4/4 | USA | North America |
| A/Backyard | Backyard bird | 2022/3/27 | USA | North America |
| A/Backyard | Backyard bird | 2022/4/4 | USA | North America |
| A/Backyard | Backyard bird | 2022/4/6 | USA | North America |
| A/Backyard | Backyard bird | 2022/4/6 | USA | North America |
| A/Backyard | Backyard bird | 2022/4/10 | USA | North America |
| A/Backyard | Backyard bird | 2022/4/10 | USA | North America |
| A/Backyard | Backyard bird | 2022/4/18 | USA | North America |
| A/Backyard | Backyard bird | 2022/4/19 | USA | North America |
| A/Backyard | Backyard bird | 2022/3/26 | USA | North America |
| A/Backyard | Backyard bird | 2022/3/25 | USA | North America |
| A/Backyard | Backyard bird | 2022/2/11 | USA | North America |
| A/Backyard | Backyard bird | 2022/3/31 | USA | North America |
| A/Backyard | Backyard bird | 2022/4/5 | USA | North America |
| A/Backyard | Backyard bird | 2022/4/5 | USA | North America |
| A/Backyard | Backyard bird | 2022/4/11 | USA | North America |
| A/Backyard | Backyard bird | 2022/4/28 | USA | North America |
| A/Backyard | Backyard bird | 2022/4/28 | USA | North America |
| A/Backyard | Backyard bird | 2022/4/28 | USA | North America |
| A/Backyard | Backyard bird | 2022/4/26 | USA | North America |
| A/Backyard | Backyard bird | 2022/4/26 | USA | North America |
| A/Backyard | Backyard bird | 2022/4/26 | USA | North America |
| A/Backyard | Backyard bird | 2022/4/26 | USA | North America |
| A/Backyard | Backyard bird | 2022/3/29 | USA | North America |
| A/Backyard | Backyard bird | 2022/3/26 | USA | North America |
| A/Backyard | Backyard bird | 2022/3/26 | USA | North America |
| A/Backyard | Backyard bird | 2022/3/26 | USA | North America |

| | | | | | |
|---|---|---|---|---|---|
| A/Backyarc | Backyard bird | | 2022/3/28 | USA | North America |
| A/Backyarc | Backyard bird | | 2022/3/31 | USA | North America |
| A/Backyarc | Backyard bird | | 2022/3/31 | USA | North America |
| A/Backyarc | Backyard bird | | 2022/3/31 | USA | North America |
| A/Baikal T | Baikal teal | Avian | 2022/3/23 | USA | North America |
| A/Bald_Ea | Bald eagle | Avian | 2022/2/16 | USA | North America |
| A/Bald Ea | Bald eagle | Avian | 2022/2/23 | USA | North America |
| A/Bald Ea | Bald eagle | Avian | Feb-22 | USA | North America |
| A/Bald_Ea | Bald eagle | Avian | Feb-22 | USA | North America |
| A/Bald Ea | Bald eagle | Avian | 2022/2/28 | USA | North America |
| A/Bald Ea | Bald eagle | Avian | 2022/2/28 | USA | North America |
| A/Bald Ea | Bald eagle | Avian | 2022/2/23 | USA | North America |
| A/Bald Ea | Bald eagle | Avian | Mar-22 | USA | North America |
| A/Bald Ea | Bald eagle | Avian | Mar-22 | USA | North America |
| A/Bald_Ea | Bald eagle | Avian | 2022/1/25 | USA | North America |
| A/bald eag | Haliaeetus leucocephalus | | 2022/2/12 | USA | North America |
| A/bald eag | Haliaeetus leucocephalus | | 2022/2/12 | | |
| A/bald_eag | Haliaeetus leucocephalus | | 2022/2/28 | | |
| A/bald eag | Haliaeetus leucocephalus | | 2022/2/28 | | |
| A/bald eag | Haliaeetus leucocephalus | | 2022/3/3 | USA | North America |
| A/bald_eag | Haliaeetus leucocephalus | | 2022/3/8 | USA | North America |
| A/bald eag | Haliaeetus leucocephalus | | 2022/3/13 | USA | North America |
| A/bald_eag | Haliaeetus leucocephalus | | 2022/2/17 | USA | North America |
| A/Bald_Ea | Bald eagle | Avian | 2022/3/8 | USA | North America |
| A/Bald Ea | Bald eagle | Avian | 2022/3/1 | USA | North America |
| A/Bald Ea | Bald eagle | Avian | 2022/3/1 | USA | North America |
| A/bald_eag | Haliaeetus leucocephalus | | 2022/3/8 | USA | North America |
| A/bald eag | Haliaeetus leucocephalus | | 2022/3/14 | USA | North America |
| A/bald_eag | Haliaeetus leucocephalus | | 2022/3/1 | USA | North America |
| A/Bald_Ea | Bald eagle | Avian | 2022/3/31 | USA | North America |
| A/Bald Ea | Bald eagle | Avian | 2022/4/13 | USA | North America |
| A/Bald_Ea | Bald eagle | Avian | 2022/4/13 | USA | North America |
| A/Bald Ea | Bald eagle | Avian | 2022/3/6 | USA | North America |
| A/Bald Ea | Bald eagle | Avian | 2022/3/14 | USA | North America |
| A/bald eag | Haliaeetus leucocephalus | | 2022/3/6 | | |
| A/bald_eag | Haliaeetus leucocephalus | | 2022/3/13 | USA | North America |
| A/bald eag | Haliaeetus leucocephalus | | Apr-22 | USA | North America |
| A/Bald_Ea | Bald eagle | Avian | 2022/3/4 | USA | North America |
| A/Bald_Ea | Bald eagle | Avian | 2022/4/11 | USA | North America |
| A/Bald Ea | Bald eagle | Avian | 2022/4/11 | USA | North America |
| A/Bald_Ea | Bald eagle | Avian | 2022/4/11 | USA | North America |
| A/Bald Ea | Bald eagle | Avian | 2022/4/14 | USA | North America |
| A/Bald Ea | Bald eagle | Avian | 2022/4/22 | USA | North America |
| A/Bald_Ea | Bald eagle | Avian | 2022/4/25 | USA | North America |
| A/Bald_Ea | Bald eagle | Avian | 2022/3/20 | USA | North America |
| A/Bald Ea | Bald eagle | Avian | 2022/3/30 | USA | North America |
| A/Bald_Ea | Bald eagle | Avian | 2022/3/30 | USA | North America |
| A/Bald_Ea | Bald eagle | Avian | 2022/3/30 | USA | North America |
| A/Bald Ea | Bald eagle | Avian | 2022/4/6 | USA | North America |
| A/Bald_Ea | Bald eagle | Avian | 2022/4/5 | USA | North America |
| A/Bald Ea | Bald eagle | Avian | 2022/4/8 | USA | North America |
| A/Bald Ea | Bald eagle | Avian | 2022/4/6 | USA | North America |
| A/Bald_Ea | Bald eagle | Avian | 2022/4/7 | USA | North America |
| A/Bald_Ea | Bald eagle | Avian | 2022/4/8 | USA | North America |
| A/Bald Ea | Bald eagle | Avian | 2022/4/11 | USA | North America |
| A/Bald Ea | Bald eagle | Avian | 2022/4/11 | USA | North America |
| A/Bald_Ea | Bald eagle | Avian | 2022/4/14 | USA | North America |
| A/Bald Ea | Bald eagle | Avian | 2022/4/11 | USA | North America |

| | | | | | |
|---|---|---|---|---|---|
| A/Bald Ea | Bald eagle | Avian | 2022/4/14 | USA | North America |
| A/Bald Ea | Bald eagle | Avian | 2022/4/25 | USA | North America |
| A/Bald Ea | Bald eagle | Avian | 2022/4/25 | USA | North America |
| A/Bald Ea | Bald eagle | Avian | 2022/4/25 | USA | North America |
| A/Bald Ea | Bald eagle | Avian | 2022/4/22 | USA | North America |
| A/Bald_Ea | Bald eagle | Avian | Mar-22 | USA | North America |
| A/Bald Ea | Bald eagle | Avian | 2022/3/18 | USA | North America |
| A/Bald Ea | Bald eagle | Avian | Mar-22 | USA | North America |
| A/Bald_Ea | Bald eagle | Avian | 2022/4/5 | USA | North America |
| A/Bald Ea | Bald eagle | Avian | 2022/3/15 | USA | North America |
| A/Bald Ea | Bald eagle | Avian | 2022/4/19 | USA | North America |
| A/Bald Ea | Bald eagle | Avian | 2022/3/30 | USA | North America |
| A/Bald Ea | Bald eagle | Avian | 2022/4/6 | USA | North America |
| A/Bald Ea | Bald eagle | Avian | 2022/4/12 | USA | North America |
| A/Bald_Ea | Bald eagle | Avian | 2022/4/21 | USA | North America |
| A/Bald Ea | Bald eagle | Avian | Feb-22 | USA | North America |
| A/Bald Ea | Bald eagle | Avian | 2022/3/8 | USA | North America |
| A/Bald_Ea | Bald eagle | Avian | 2022/3/8 | USA | North America |
| A/Bald Ea | Bald eagle | Avian | 2022/3/31 | USA | North America |
| A/Bald Ea | Bald eagle | Avian | 2022/3/31 | USA | North America |
| A/bald_eag | Haliaeetus leucocephalus | | 2022/2/22 | | |
| A/bald eag | Haliaeetus leucocephalus | | 2022/3/8 | | |
| A/bald_eag | Haliaeetus leucocephalus | | 2022/3/12 | | |
| A/bald_eag | Haliaeetus leucocephalus | | 2022/3/10 | USA | North America |
| A/bald eag | Haliaeetus leucocephalus | | 2022/12/30 | USA | North America |
| A/bald eag | Haliaeetus leucocephalus | | 2023/2/16 | USA | North America |
| A/bald_eag | Haliaeetus leucocephalus | | 2023/2/19 | USA | North America |
| A/Bald Ea | Bald eagle | Avian | 2022/3/28 | USA | North America |
| A/Bald_Ea | Bald eagle | Avian | 2022/3/27 | USA | North America |
| A/Bald_Ea | Bald eagle | Avian | 2022/4/11 | USA | North America |
| A/Bald Ea | Bald eagle | Avian | 2022/4/8 | USA | North America |
| A/Bald_Ea | Bald eagle | Avian | 2022/4/21 | USA | North America |
| A/Bald Ea | Bald eagle | Avian | 2022/3/2 | USA | North America |
| A/Bald Ea | Bald eagle | Avian | 2022/3/7 | USA | North America |
| A/Bald Ea | Bald eagle | Avian | 2022/3/4 | USA | North America |
| A/Bald_Ea | Bald eagle | Avian | 2022/3/25 | USA | North America |
| A/Bald Ea | Bald eagle | Avian | 2022/4/2 | USA | North America |
| A/Bald_Ea | Bald eagle | Avian | 2022/4/13 | USA | North America |
| A/Bald_Ea | Bald eagle | Avian | 2022/3/19 | USA | North America |
| A/Bald Ea | Bald eagle | Avian | 2022/3/25 | USA | North America |
| A/Bald_Ea | Bald eagle | Avian | 2022/2/26 | USA | North America |
| A/Bald Ea | Bald eagle | Avian | 2022/2/26 | USA | North America |
| A/Bald Ea | Bald eagle | Avian | 2022/3/4 | USA | North America |
| A/Bald_Ea | Bald eagle | Avian | 2022/3/4 | USA | North America |
| A/Bald_Ea | Bald eagle | Avian | 2022/3/1 | USA | North America |
| A/Bald Ea | Bald eagle | Avian | 2022/3/17 | USA | North America |
| A/bald_eag | Haliaeetus leucocephalus | | 2022/3/1 | USA | North America |
| A/bald_eag | Haliaeetus leucocephalus | | 2023/3/4 | USA | North America |
| A/bald eag | Haliaeetus leucocephalus | | 2023/3/4 | USA | North America |
| A/Bald_Ea | Bald eagle | Avian | 2022/3/23 | USA | North America |
| A/Bald Ea | Bald eagle | Avian | 2022/3/9 | USA | North America |
| A/Bald Ea | Bald eagle | Avian | Mar-22 | USA | North America |
| A/bald_eag | Haliaeetus leucocephalus | | 2022/12/29 | USA | North America |
| A/Bald_Ea | Bald eagle | Avian | 2022/3/28 | USA | North America |
| A/Bald Ea | Bald eagle | Avian | 2022/3/28 | USA | North America |
| A/Bald Ea | Bald eagle | Avian | 2022/3/30 | USA | North America |
| A/bald_eag | Haliaeetus leucocephalus | | 2023/1/3 | USA | North America |
| A/bald eag | Haliaeetus leucocephalus | | 2022/12/8 | USA | North America |

| | | | | | |
|---|---|---|---|---|---|
| A/bald eag | Haliaeetus leucocephalus | | 2022/4/15 | | |
| A/Bald Ea | Bald eagle | Avian | 2022/3/14 | USA | North America |
| A/Bald Ea | Bald eagle | Avian | 2022/3/28 | USA | North America |
| A/Bald Ea | Bald eagle | Avian | 2022/4/9 | USA | North America |
| A/Bald Ea | Bald eagle | Avian | 2022/4/26 | USA | North America |
| A/Bald_Ea | Bald eagle | Avian | 2022/4/22 | USA | North America |
| A/Barred ( | Barred owl | | 2022/4/15 | USA | North America |
| A/Barred ( | Barred owl | | 2022/4/27 | USA | North America |
| A/Belchers | Belcher's_gull | | Nov-22 | Peru | South America |
| A/Belchers | Belcher's gull | | Dec-22 | Peru | South America |
| A/Black-cr | Nycticorax nycticorax | Avian | 2022/12/20 | Chile | South America |
| A/Black-cr | Nycticorax nycticorax | Avian | 2022/12/20 | Chile | South America |
| A/Blackish | Haematopus ater | Avian | 2023/3/6 | Chile | South America |
| A/black sk | Rynchops niger | Avian | 2022/12/2 | Chile | South America |
| A/Black_Sl | Rynchops niger | Avian | 2023/3/6 | Chile | South America |
| A/Black S | Black swift | | 2022/3/9 | USA | North America |
| A/Black S | Black swift | | 2022/3/9 | USA | North America |
| A/Black_V | Black vulture | Avian | 2022/2/14 | USA | North America |
| A/Black V | Black vulture | Avian | 2022/2/14 | USA | North America |
| A/Black V | Black vulture | Avian | 2022/3/3 | USA | North America |
| A/Black_V | Black vulture | Avian | 2022/3/2 | USA | North America |
| A/Black V | Black vulture | Avian | Mar-22 | USA | North America |
| A/Black_V | Black vulture | Avian | 2022/3/5 | USA | North America |
| A/Black_V | Black vulture | Avian | 2022/3/7 | USA | North America |
| A/Black V | Black vulture | Avian | 2022/3/7 | USA | North America |
| A/Black V | Black vulture | Avian | 2022/3/9 | USA | North America |
| A/Black_V | Black vulture | Avian | 2022/3/9 | USA | North America |
| A/Black V | Black vulture | Avian | 2022/2/25 | USA | North America |
| A/Black_V | Black vulture | Avian | 2022/3/7 | USA | North America |
| A/Black_V | Black vulture | Avian | 2022/3/7 | USA | North America |
| A/Black V | Black vulture | Avian | 2022/3/14 | USA | North America |
| A/Black_V | Black vulture | Avian | 2022/3/14 | USA | North America |
| A/Black V | Black vulture | Avian | 2022/3/14 | USA | North America |
| A/Black V | Black vulture | Avian | 2022/3/14 | USA | North America |
| A/Black V | Black vulture | Avian | 2022/3/14 | USA | North America |
| A/Black_V | Black vulture | Avian | 2022/3/14 | USA | North America |
| A/Black V | Black vulture | Avian | 2022/3/14 | USA | North America |
| A/Black_V | Black vulture | Avian | 2022/3/17 | USA | North America |
| A/Black_V | Black vulture | Avian | 2022/3/18 | USA | North America |
| A/Black V | Black vulture | Avian | 2022/3/21 | USA | North America |
| A/Black_V | Black vulture | Avian | 2022/3/21 | USA | North America |
| A/Black V | Black vulture | Avian | 2022/3/21 | USA | North America |
| A/Black V | Black vulture | Avian | 2022/3/21 | USA | North America |
| A/Black_V | Black vulture | Avian | 2022/3/21 | USA | North America |
| A/Black_V | Black vulture | Avian | 2022/3/21 | USA | North America |
| A/Black V | Black vulture | Avian | 2022/3/25 | USA | North America |
| A/Black_V | Black vulture | Avian | 2022/3/25 | USA | North America |
| A/Black_V | Black vulture | Avian | 2022/3/27 | USA | North America |
| A/Black V | Black vulture | Avian | 2022/3/24 | USA | North America |
| A/Black_V | Black vulture | Avian | 2022/3/30 | USA | North America |
| A/Black V | Black vulture | Avian | 2022/3/30 | USA | North America |
| A/Black V | Black vulture | Avian | 2022/3/31 | USA | North America |
| A/Black_V | Black vulture | Avian | 2022/3/31 | USA | North America |
| A/Black_V | Black vulture | Avian | 2022/3/31 | USA | North America |
| A/Black V | Black vulture | Avian | 2022/3/31 | USA | North America |
| A/Black V | Black vulture | Avian | 2022/3/31 | USA | North America |
| A/Black_V | Black vulture | Avian | 2022/4/4 | USA | North America |
| A/Black V | Black vulture | Avian | 2022/4/5 | USA | North America |

| | | | | | |
|---|---|---|---|---|---|
| A/Black V | Black vulture | Avian | 2022/4/4 | USA | North America |
| A/Black V | Black vulture | Avian | 2022/4/5 | USA | North America |
| A/Black V | Black vulture | Avian | 2022/4/5 | USA | North America |
| A/Black V | Black vulture | Avian | 2022/4/6 | USA | North America |
| A/Black V | Black vulture | Avian | 2022/4/6 | USA | North America |
| A/Black_V | Black vulture | Avian | 2022/4/8 | USA | North America |
| A/Black V | Black vulture | Avian | 2022/4/5 | USA | North America |
| A/Black V | Black vulture | Avian | 2022/4/13 | USA | North America |
| A/Black_V | Black vulture | Avian | 2022/4/15 | USA | North America |
| A/Black V | Black vulture | Avian | 2022/4/16 | USA | North America |
| A/Black V | Black vulture | Avian | 2022/4/15 | USA | North America |
| A/Black V | Black vulture | Avian | 2022/4/19 | USA | North America |
| A/Black V | Black vulture | Avian | 2022/4/20 | USA | North America |
| A/Black V | Black vulture | Avian | 2022/4/20 | USA | North America |
| A/Black_V | Black vulture | Avian | 2022/4/20 | USA | North America |
| A/Black V | Black vulture | Avian | 2022/4/21 | USA | North America |
| A/Black V | Black vulture | Avian | 2022/4/21 | USA | North America |
| A/Black_V | Black vulture | Avian | 2022/4/21 | USA | North America |
| A/Black V | Black vulture | Avian | 2022/4/22 | USA | North America |
| A/Black V | Black vulture | Avian | 2022/4/22 | USA | North America |
| A/Black_V | Black vulture | Avian | 2022/4/22 | USA | North America |
| A/black vu | Coragyps atratus | | 2022/2/25 | | |
| A/black_vu | Coragyps atratus | | 2022/3/4 | | |
| A/black_vu | Coragyps atratus | | 2022/3/7 | | |
| A/black vu | Coragyps atratus | Avian | 2022/10/21 | USA | North America |
| A/black vu | Coragyps atratus | Avian | 2022/10/21 | USA | North America |
| A/black_vu | Coragyps atratus | Avian | 2022/11/2 | USA | North America |
| A/black vu | Coragyps atratus | Avian | 2022/11/3 | USA | North America |
| A/black_vu | Coragyps atratus | Avian | 2022/11/3 | USA | North America |
| A/black_vu | Coragyps atratus | Avian | Apr-22 | USA | North America |
| A/black vu | Coragyps atratus | Avian | Apr-22 | USA | North America |
| A/black_vu | Coragyps atratus | Avian | Apr-22 | USA | North America |
| A/black vu | Coragyps atratus | Avian | 2022/6/23 | USA | North America |
| A/black vu | Coragyps atratus | Avian | 2022/7/1 | USA | North America |
| A/black vu | Coragyps atratus | Avian | 2022/8/22 | USA | North America |
| A/black_vu | Coragyps atratus | Avian | 2022/8/22 | USA | North America |
| A/black vu | Coragyps atratus | Avian | 2022/8/24 | USA | North America |
| A/black_vu | Coragyps atratus | Avian | 2022/8/23 | USA | North America |
| A/black_vu | Coragyps atratus | Avian | 2022/8/23 | USA | North America |
| A/black vu | Coragyps atratus | Avian | 2022/8/23 | USA | North America |
| A/black_vu | Coragyps atratus | Avian | 2022/8/31 | USA | North America |
| A/black vu | Coragyps atratus | Avian | 2022/8/31 | USA | North America |
| A/black vu | Coragyps atratus | Avian | 2022/8/31 | USA | North America |
| A/black_vu | Coragyps atratus | Avian | 2022/8/30 | USA | North America |
| A/black_vu | Coragyps atratus | Avian | 2022/8/30 | USA | North America |
| A/black vu | Coragyps atratus | Avian | 2022/8/31 | USA | North America |
| A/black_vu | Coragyps atratus | Avian | 2022/8/31 | USA | North America |
| A/black_vu | Coragyps atratus | Avian | 2022/9/6 | USA | North America |
| A/black vu | Coragyps atratus | Avian | 2022/9/6 | USA | North America |
| A/black_vu | Coragyps atratus | Avian | Sep-22 | USA | North America |
| A/black vu | Coragyps atratus | Avian | Sep-22 | USA | North America |
| A/black vu | Coragyps atratus | Avian | Sep-22 | USA | North America |
| A/black_vu | Coragyps atratus | Avian | Sep-22 | USA | North America |
| A/black_vu | Coragyps atratus | Avian | Sep-22 | USA | North America |
| A/black vu | Coragyps atratus | Avian | Sep-22 | USA | North America |
| A/black vu | Coragyps atratus | Avian | 2022/9/29 | USA | North America |
| A/black_vu | Coragyps atratus | Avian | Oct-22 | USA | North America |
| A/black vu | Coragyps atratus | Avian | Oct-22 | USA | North America |

| | | | | | |
|---|---|---|---|---|---|
| A/black vu | Coragyps atratus | Avian | 2023/2/7 | USA | North America |
| A/black vu | Coragyps atratus | Avian | 2023/2/8 | USA | North America |
| A/black vu | Coragyps atratus | Avian | 2023/2/18 | USA | North America |
| A/Black V | Black vulture | Avian | 2022/4/21 | USA | North America |
| A/Black V | Black vulture | Avian | 2022/4/26 | USA | North America |
| A/black_vu | Coragyps atratus | Avian | 2022/10/31 | USA | North America |
| A/black vu | Coragyps atratus | Avian | 2022/11/4 | USA | North America |
| A/black vu | Coragyps atratus | Avian | 2022/11/4 | USA | North America |
| A/black_vu | Coragyps atratus | Avian | 2022/11/22 | USA | North America |
| A/black vu | Coragyps atratus | | 2022/5/10 | | |
| A/black vu | Coragyps atratus | Avian | 2022/11/2 | USA | North America |
| A/black vu | Coragyps atratus | Avian | 2022/11/2 | USA | North America |
| A/black vu | Coragyps atratus | Avian | 2022/12/1 | USA | North America |
| A/black vu | Coragyps atratus | Avian | 2022/8/22 | USA | North America |
| A/black_vu | Coragyps atratus | Avian | 2022/8/25 | USA | North America |
| A/black vu | Coragyps atratus | Avian | 2022/12/5 | USA | North America |
| A/black vu | Coragyps atratus | Avian | 2022/12/5 | USA | North America |
| A/black_vu | Coragyps atratus | | 2022/6/30 | | |
| A/black vu | Coragyps atratus | | 2022/6/30 | | |
| A/black vu | Coragyps atratus | | 2022/6/30 | | |
| A/black_vu | Coragyps atratus | Avian | 2022/8/22 | USA | North America |
| A/black vu | Coragyps atratus | Avian | 2022/8/22 | USA | North America |
| A/black_vu | Coragyps atratus | Avian | 2022/8/22 | USA | North America |
| A/black_vu | Coragyps atratus | Avian | 2022/8/22 | USA | North America |
| A/black vu | Coragyps atratus | Avian | 2022/8/22 | USA | North America |
| A/black vu | Coragyps atratus | Avian | 2022/8/22 | USA | North America |
| A/Blue-wir | Blue-winged and cinnamon teal | | 2022/1/6 | USA | North America |
| A/Blue-wir | Blue-winged teal | Avian | 2022/1/22 | USA | North America |
| A/blue-win | Spatula discors | Avian | 2022/9/23 | USA | North America |
| A/blue-win | Spatula discors | Avian | 2022/9/23 | USA | North America |
| A/blue-win | Spatula discors | Avian | Sep-22 | USA | North America |
| A/blue-win | Spatula discors | Avian | Sep-22 | USA | North America |
| A/blue-win | Spatula discors | Avian | Sep-22 | USA | North America |
| A/blue-win | Spatula discors | Avian | 2022/9/15 | USA | North America |
| A/blue-win | Spatula discors | Avian | 2022/9/15 | USA | North America |
| A/blue-win | Spatula discors | Avian | 2022/9/15 | USA | North America |
| A/blue-win | Spatula discors | Avian | 2022/9/14 | USA | North America |
| A/Blue-wir | Blue-winged teal | Avian | 2021/12/30 | USA | North America |
| A/blue-win | Spatula discors | Avian | Sep-22 | USA | North America |
| A/blue-win | Spatula discors | Avian | Sep-22 | USA | North America |
| A/blue-win | Spatula discors | Avian | Oct-22 | USA | North America |
| A/blue-win | Spatula discors | Avian | Oct-22 | USA | North America |
| A/blue-win | Spatula discors | Avian | Oct-22 | USA | North America |
| A/blue-win | Spatula discors | Avian | Oct-22 | USA | North America |
| A/blue-win | Spatula discors | Avian | Oct-22 | USA | North America |
| A/bottlenos | Tursiops truncatus | Sea Mammal | 2022/3/30 | USA | North America |
| A/Brown_I | Brown pelican | | 2022/3/4 | USA | North America |
| A/Brown_I | Brown pelican | | 2022/2/20 | USA | North America |
| A/brown p | Pelecanus occidentalis | | 2022/2/20 | | |
| A/Canada_ | Canada goose | Avian | 2022/4/1 | USA | North America |
| A/Canada | Canada goose | Avian | 2022/4/26 | USA | North America |
| A/Canada | Canada goose | Avian | 2022/2/4 | USA | North America |
| A/Canada_ | Canada goose | Avian | 2022/4/5 | USA | North America |
| A/Canada_ | Canada goose | Avian | 2022/3/16 | USA | North America |
| A/Canada | Canada goose | Avian | 2022/2/3 | USA | North America |
| A/Canada | Canada goose | Avian | 2022/2/3 | USA | North America |
| A/Canada_ | Canada goose | Avian | 2022/3/8 | USA | North America |
| A/Canada | Canada goose | Avian | 2022/3/17 | USA | North America |

| | | | | | |
|---|---|---|---|---|---|
| A/Canada | Canada goose | Avian | 2022/3/17 | USA | North America |
| A/Canada | Canada goose | Avian | 2022/3/17 | USA | North America |
| A/Canada | Canada goose | Avian | 2022/3/17 | USA | North America |
| A/Canada | Canada goose | Avian | 2022/3/17 | USA | North America |
| A/Canada | Canada goose | Avian | 2022/3/17 | USA | North America |
| A/Canada_ | Canada goose | Avian | 2022/3/25 | USA | North America |
| A/Canada | Canada goose | Avian | 2022/3/25 | USA | North America |
| A/Canada | Canada goose | Avian | 2022/4/5 | USA | North America |
| A/Canada_ | Canada goose | Avian | 2022/3/18 | USA | North America |
| A/Canada | Canada goose | Avian | 2022/3/18 | USA | North America |
| A/Canada | Canada goose | Avian | Mar-22 | USA | North America |
| A/Canada | Canada goose | Avian | Apr-22 | USA | North America |
| A/Canada | Canada goose | Avian | Mar-22 | USA | North America |
| A/Canada | Canada goose | Avian | 2022/4/11 | USA | North America |
| A/Canada_ | Canada goose | Avian | 2022/4/20 | USA | North America |
| A/Canada | Canada goose | Avian | 2022/3/7 | USA | North America |
| A/Canada | Canada goose | Avian | 2022/3/7 | USA | North America |
| A/Canada_ | Canada goose | Avian | 2022/2/22 | USA | North America |
| A/Canada | Canada goose | Avian | 2022/2/22 | USA | North America |
| A/Canada | Canada goose | Avian | 2022/2/22 | USA | North America |
| A/Canada_ | Canada goose | Avian | 2022/4/7 | USA | North America |
| A/Canada | Canada goose | Avian | 2022/3/21 | USA | North America |
| A/Canada_ | Branta canadensis | Avian | 2022/11/1 | USA | North America |
| A/Canada_ | Canada goose | Avian | 2022/3/25 | USA | North America |
| A/Canada | Canada goose | Avian | 2022/3/1 | USA | North America |
| A/Canada | Canada goose | Avian | 2022/3/15 | USA | North America |
| A/Canada_ | Canada goose | Avian | 2022/3/15 | USA | North America |
| A/Canada | Canada goose | Avian | 2022/3/9 | USA | North America |
| A/Canada_ | Canada goose | Avian | 2022/3/16 | USA | North America |
| A/Canada_ | Canada goose | Avian | 2022/3/14 | USA | North America |
| A/Canada | Canada goose | Avian | 2022/3/14 | USA | North America |
| A/Canada_ | Canada goose | Avian | 2022/3/14 | USA | North America |
| A/Canada | Canada goose | Avian | 2022/4/1 | USA | North America |
| A/Canada | Canada goose | Avian | 2022/4/18 | USA | North America |
| A/Canada | Branta canadensis | Avian | 2022/9/29 | USA | North America |
| A/Canada_ | Canada goose | Avian | 2022/3/20 | USA | North America |
| A/Canada | Canada goose | Avian | 2022/3/28 | USA | North America |
| A/Canada_ | Canada goose | Avian | 2022/4/2 | USA | North America |
| A/Canada_ | Canada goose | Avian | 2022/4/4 | USA | North America |
| A/Canada | Canada goose | Avian | 2022/3/31 | USA | North America |
| A/Canada_ | Canada goose | Avian | 2022/4/10 | USA | North America |
| A/Canada | Canada goose | Avian | 2022/4/13 | USA | North America |
| A/Canada | Canada goose | Avian | 2022/4/18 | USA | North America |
| A/Canada_ | Canada goose | Avian | 2022/4/21 | USA | North America |
| A/chicken/. | Gallus gallus | Avian | 2023/2/28 | Chile | South America |
| A/chicken/. | Gallus gallus | Avian | 2023/3/2 | Chile | South America |
| A/chicken/. | Gallus gallus | Avian | 2023/3/2 | Chile | South America |
| A/chicken/. | Gallus gallus | Avian | 2023/3/7 | Chile | South America |
| A/chicken/. | Gallus gallus | Avian | 2023/3/14 | Chile | South America |
| A/chicken/. | Gallus gallus | Avian | 2023/3/14 | Chile | South America |
| A/chicken/( | Chicken | Avian | 2022/11/8 | Colombi: | South America |
| A/chicken/( | Chicken | Avian | 2022/11/13 | Colombi: | South America |
| A/chicken/( | Chicken | Avian | 2022/11/1 | Colombi: | South America |
| A/chicken/( | Chicken | Avian | 2022/11/26 | Colombi: | South America |
| A/chicken/( | Chicken | Avian | 2022/11/18 | Colombi: | South America |
| A/chicken/] | Chicken | Avian | 2021/3/1 | Nigeria | Africa |
| A/chicken/] | Gallus gallus | Avian | 2023/2/27 | Chile | South America |
| A/chicken/] | Gallus gallus | Avian | 2023/3/6 | Chile | South America |

| | | | | | |
|---|---|---|---|---|---|
| A/chicken/I | Gallus gallus | Avian | 2023/3/8 | Chile | South America |
| A/chicken/I | Gallus gallus | Avian | 2023/3/10 | Chile | South America |
| A/chicken/I | Gallus gallus | Avian | 2023/3/10 | Chile | South America |
| A/chicken/( | Gallus gallus | Avian | 2023/3/12 | Chile | South America |
| A/chicken/( | Gallus gallus | Avian | 2023/3/12 | Chile | South America |
| A/chicken/( | Gallus gallus | Avian | 2023/3/12 | Chile | South America |
| A/chicken/( | Chicken | Avian | 2022/9/1 | USA | North America |
| A/chicken/( | Chicken | Avian | 2022/9/1 | USA | North America |
| A/chicken/( | Chicken | Avian | 2022/9/12 | USA | North America |
| A/chicken/( | Chicken | Avian | 2022/11/3 | USA | North America |
| A/chicken/( | Chicken | Avian | 2022/11/3 | USA | North America |
| A/chicken/( | Chicken | Avian | 2022/11/3 | USA | North America |
| A/chicken/( | Chicken | Avian | 2022/3/28 | USA | North America |
| A/Chiloe v | Mareca sibilatrix | | 2023/3/9 | Chile | South America |
| A/common | Somateria mollissima | | 2022/6/23 | USA | North America |
| A/common | Somateria mollissima | | 2022/6/23 | USA | North America |
| A/common | Somateria mollissima | | 2022/6/23 | USA | North America |
| A/common | Somateria mollissima | | 2022/6/23 | USA | North America |
| A/common | Somateria mollissima | | 2022/6/12 | | |
| A/common | Somateria mollissima | | 2022/6/12 | | |
| A/Commor | Common goldeneye | Avian | 2022/3/31 | USA | North America |
| A/Commor | Common loon | Avian | 2022/3/31 | USA | North America |
| A/Commor | Common raven | Avian | 2022/3/24 | USA | North America |
| A/common | Sterna hirundo | | 2022/6/23 | | |
| A/common | Sterna hirundo | | 2022/6/23 | | |
| A/Coopers | Accipiter cooperii | | 2022/12/2 | USA | North America |
| A/Coopers_ | Cooper's hawk | Avian | 2022/4/25 | USA | North America |
| A/Coopers | Cooper's hawk | Avian | 2022/4/22 | USA | North America |
| A/Coopers_ | Cooper's hawk | Avian | 2022/4/27 | USA | North America |
| A/Coopers_ | Cooper's hawk | Avian | 2022/3/31 | USA | North America |
| A/Coopers | Cooper's hawk | Avian | 2022/4/8 | USA | North America |
| A/Coopers_ | Cooper's hawk | Avian | 2022/4/21 | USA | North America |
| A/Coopers | Cooper's hawk | Avian | 2022/3/13 | USA | North America |
| A/Cormora | Cormorant | Avian | Apr-22 | USA | North America |
| A/Cormora | Cormorant | Avian | Apr-22 | USA | North America |
| A/dolphin/I | Common dolphin | | 2022/11/22 | Peru | South America |
| A/Domesti | Anas platyrhynchos do | Avian | 2023/2/28 | Chile | South America |
| A/Domesti | Anas platyrhynchos do | Avian | 2023/3/7 | Chile | South America |
| A/Domesti | Anas platyrhynchos do | Avian | 2023/3/14 | Chile | South America |
| A/Domesti | Anas platyrhynchos do | Avian | 2023/3/7 | Chile | South America |
| A/double-c | Phalacrocorax auritus | | 2022/12/6 | USA | North America |
| A/Duck/Fl | Duck | Avian | 2022/3/3 | USA | North America |
| A/Duck/Ida | Duck | Avian | 2022/4/18 | USA | North America |
| A/Duck/Ida | Duck | Avian | 2022/4/18 | USA | North America |
| A/Duck/Inc | Duck | Avian | 2022/4/7 | USA | North America |
| A/Duck/Inc | Duck | Avian | 2022/4/12 | USA | North America |
| A/Duck/Inc | Duck | Avian | 2022/4/17 | USA | North America |
| A/Duck/Inc | Duck | Avian | 2022/4/17 | USA | North America |
| A/Duck/M | Duck | Avian | 2022/4/6 | USA | North America |
| A/Duck/M | Duck | Avian | 2022/4/6 | USA | North America |
| A/Duck/M | Duck | Avian | 2022/4/19 | USA | North America |
| A/Duck/M | Duck | Avian | 2022/4/19 | USA | North America |
| A/Duck/M | Duck | Avian | 2022/4/27 | USA | North America |
| A/Duck/M | Duck | Avian | 2022/4/27 | USA | North America |
| A/Duck/M | Duck | Avian | 2022/4/27 | USA | North America |
| A/Duck/Pe | Duck | Avian | 2022/4/28 | USA | North America |
| A/Duck/Pe | Duck | Avian | 2022/4/28 | USA | North America |
| A/Eagle/M | Eagle | Avian | 2022/3/15 | USA | North America |

| | | | | | |
|---|---|---|---|---|---|
| A/Elegant | Thalasseus elegans | | 2022/12/24 | Chile | South America |
| A/environn | Environment | Environment | 2022/4/14 | USA | North America |
| A/Falco ru | Falco rusticolus | | Oct-22 | Mexico | North America |
| A/Fish Cro | Fish crow | | 2022/4/4 | USA | North America |
| A/Gadwall/ | Gadwall | Avian | 2022/1/30 | USA | North America |
| A/Gadwall/ | Gadwall | Avian | 2022/2/8 | USA | North America |
| A/Gadwall/ | Gadwall | Avian | 2022/1/8 | USA | North America |
| A/Gadwall/ | Gadwall | Avian | 2022/1/8 | USA | North America |
| A/Gadwall/ | Gadwall | Avian | 2022/1/8 | USA | North America |
| A/Gadwall/ | Gadwall | Avian | 2022/1/8 | USA | North America |
| A/Gadwall/ | Gadwall | Avian | 2022/1/8 | USA | North America |
| A/Gadwall/ | Gadwall | Avian | 2022/1/15 | USA | North America |
| A/Gadwall/ | Gadwall | Avian | 2022/1/15 | USA | North America |
| A/Gadwall/ | Gadwall | Avian | 2022/1/22 | USA | North America |
| A/Gadwall/ | Gadwall | Avian | 2022/1/6 | USA | North America |
| A/Gadwall/ | Gadwall | Avian | 2022/1/6 | USA | North America |
| A/Gadwall/ | Gadwall | Avian | 2022/1/6 | USA | North America |
| A/Gadwall/ | Gadwall | Avian | 2022/1/6 | USA | North America |
| A/Gadwall/ | Gadwall | Avian | 2022/1/6 | USA | North America |
| A/Gadwall/ | Gadwall | Avian | 2022/1/29 | USA | North America |
| A/Gallus_g | Gallus gallus | Avian | 2022/11/18 | Peru | South America |
| A/Gallus g | Gallus gallus | Avian | 2022/11/28 | Peru | South America |
| A/Gallus_g | Gallus gallus | Avian | 2022/12/1 | Peru | South America |
| A/Gallus_g | Gallus gallus | Avian | 2022/12/1 | Peru | South America |
| A/Gallus g | Gallus gallus | Avian | 2022/12/3 | Peru | South America |
| A/Gallus g | Gallus gallus | Avian | 2022/12/18 | Peru | South America |
| A/Gallus_g | Gallus gallus | Avian | 2022/12/22 | Peru | South America |
| A/Gallus g | Gallus gallus | Avian | 2022/12/22 | Peru | South America |
| A/Gallus_g | Gallus gallus | Avian | 2022/12/12 | Peru | South America |
| A/Gallus_g | Gallus gallus | Avian | 2022/12/12 | Peru | South America |
| A/goose/An | Anser sp. | Avian | 2023/2/28 | Chile | South America |
| A/goose/Cz | Goose | Avian | 2021/9/27 | Czech Re | Europe |
| A/goose/Ol | Goose | Avian | 2022/9/13 | USA | North America |
| A/gray gul | Leucophaeus modestus | | 2022/12/2 | Chile | South America |
| A/Gray gu | Leucophaeus modestus | | 2023/1/18 | Chile | South America |
| A/Graylag_ | Graylag goose | | 2022/3/14 | USA | North America |
| A/Graylag | Graylag goose | | 2022/3/14 | USA | North America |
| A/great_bla | Larus marinus | Avian | 2022/12/6 | USA | North America |
| A/Great_Bl | Great blue heron | | 2022/2/27 | USA | North America |
| A/great blu | Ardea herodias | | 2022/2/27 | | |
| A/Great_Bl | Great blue heron | | 2022/3/2 | USA | North America |
| A/Great eg | Ardea alba | | 2023/3/7 | Chile | South America |
| A/Greater | Greater white-fronted g | Avian | 2022/4/6 | USA | North America |
| A/Great_H | Great horned owl | Avian | 2022/4/11 | USA | North America |
| A/great_ho | Bubo virginianus | | 2022/2/24 | | |
| A/great ho | Bubo virginianus | | 2022/2/23 | | |
| A/Great_H | Great horned owl | Avian | 2022/2/28 | USA | North America |
| A/Great_H | Great horned owl | Avian | 2022/2/24 | USA | North America |
| A/Great H | Great horned owl | Avian | 2022/2/25 | USA | North America |
| A/Great_H | Great horned owl | Avian | 2022/4/7 | USA | North America |
| A/Great H | Great horned owl | Avian | 2022/4/7 | USA | North America |
| A/Great H | Great horned owl | Avian | 2022/4/15 | USA | North America |
| A/Great_H | Great horned owl | Avian | 2022/4/14 | USA | North America |
| A/Great_H | Great horned owl | Avian | 2022/3/30 | USA | North America |
| A/Great H | Great horned owl | Avian | 2022/3/30 | USA | North America |
| A/Great H | Great horned owl | Avian | 2022/4/3 | USA | North America |
| A/Great_H | Great horned owl | Avian | 2022/4/6 | USA | North America |
| A/Great H | Great horned owl | Avian | 2022/3/31 | USA | North America |

| | | | | | |
|---|---|---|---|---|---|
| A/Great H | Great horned owl | Avian | 2022/4/11 | USA | North America |
| A/Great H | Great horned owl | Avian | 2022/4/11 | USA | North America |
| A/Great H | Great horned owl | Avian | 2022/4/15 | USA | North America |
| A/Great H | Great horned owl | Avian | 2022/4/15 | USA | North America |
| A/Great H | Great horned owl | Avian | 2022/4/15 | USA | North America |
| A/Great_H | Great horned owl | Avian | 2022/4/15 | USA | North America |
| A/Great H | Great horned owl | Avian | 2022/4/15 | USA | North America |
| A/Great H | Great horned owl | Avian | 2022/4/15 | USA | North America |
| A/Great_H | Great horned owl | Avian | 2022/4/15 | USA | North America |
| A/Great H | Great horned owl | Avian | Apr-22 | USA | North America |
| A/Great H | Great horned owl | Avian | 2022/4/11 | USA | North America |
| A/Great H | Great horned owl | Avian | 2022/4/14 | USA | North America |
| A/Great H | Great horned owl | Avian | 2022/4/12 | USA | North America |
| A/Great H | Great horned owl | Avian | 2022/4/18 | USA | North America |
| A/Great_H | Great horned owl | Avian | 2022/4/13 | USA | North America |
| A/Great H | Great horned owl | Avian | 2022/4/25 | USA | North America |
| A/Great H | Great horned owl | Avian | 2022/4/25 | USA | North America |
| A/Great_H | Great horned owl | Avian | 2022/4/25 | USA | North America |
| A/Great H | Great horned owl | Avian | 2022/4/25 | USA | North America |
| A/Great H | Great horned owl | Avian | 2022/4/25 | USA | North America |
| A/Great_H | Great horned owl | Avian | 2022/4/21 | USA | North America |
| A/Great H | Great horned owl | Avian | 2022/4/22 | USA | North America |
| A/Great_H | Great horned owl | Avian | 2022/4/22 | USA | North America |
| A/Great_H | Great horned owl | Avian | 2022/4/22 | USA | North America |
| A/Great H | Great horned owl | Avian | 2022/4/27 | USA | North America |
| A/Great H | Great horned owl | Avian | 2022/4/27 | USA | North America |
| A/Great_H | Great horned owl | Avian | 2022/4/7 | USA | North America |
| A/Great H | Great horned owl | Avian | 2022/4/7 | USA | North America |
| A/Great_H | Great horned owl | Avian | Apr-22 | USA | North America |
| A/Great_H | Great horned owl | Avian | 2022/3/27 | USA | North America |
| A/Great H | Great horned owl | Avian | 2022/4/5 | USA | North America |
| A/Great_H | Great horned owl | Avian | 2022/4/11 | USA | North America |
| A/Great H | Great horned owl | Avian | 2022/4/21 | USA | North America |
| A/Great H | Great horned owl | Avian | 2022/4/21 | USA | North America |
| A/Great H | Great horned owl | Avian | 2022/3/18 | USA | North America |
| A/Great_H | Great horned owl | Avian | 2022/3/22 | USA | North America |
| A/Great H | Great horned owl | Avian | 2022/3/29 | USA | North America |
| A/Great_H | Great horned owl | Avian | 2022/4/4 | USA | North America |
| A/Great_H | Great horned owl | Avian | 2022/4/4 | USA | North America |
| A/Great H | Great horned owl | Avian | 2022/4/13 | USA | North America |
| A/Great_H | Great horned owl | Avian | 2022/4/22 | USA | North America |
| A/Great H | Great horned owl | Avian | 2022/4/25 | USA | North America |
| A/great-tail | Quiscalus mexicanus | | 2022/12/13 | USA | North America |
| A/great-tail | Quiscalus mexicanus | | 2022/12/13 | USA | North America |
| A/great-tail | Quiscalus mexicanus | | 2022/12/13 | USA | North America |
| A/Green-w | Green-winged teal | Avian | 2022/1/8 | USA | North America |
| A/Green-w | Green-winged teal | Avian | 2022/1/29 | USA | North America |
| A/Green-w | Green-winged teal | Avian | 2022/1/29 | USA | North America |
| A/Green-w | Green-winged teal | Avian | 2022/1/29 | USA | North America |
| A/Green-w | Green-winged teal | Avian | 2022/1/29 | USA | North America |
| A/Green-w | Green-winged teal | Avian | 2022/1/29 | USA | North America |
| A/Green-w | Green-winged teal | Avian | 2022/1/31 | USA | North America |
| A/Green-w | Green-winged teal | Avian | 2022/2/5 | USA | North America |
| A/Green-w | Green-winged teal | Avian | 2022/1/19 | USA | North America |
| A/Guanay | Guanay cormorant | | 2022/11/24 | Peru | South America |
| A/Guineafc | Guineafowl | Avian | 2022/3/9 | USA | North America |
| A/Gull/CH | Larus sp. | Avian | 2022/12/7 | Chile | South America |
| A/Gull/CH | Larus sp. | Avian | 2022/12/7 | Chile | South America |

| | | | | | |
|---|---|---|---|---|---|
| A/Gull/Flo | Gull | Avian | 2022/3/1 | USA | North America |
| A/Gull/Flo | Gull | Avian | 2022/3/31 | USA | North America |
| A/gull/Flor | Larus sp. | | 2022/3/1 | | |
| A/harbor s | Phoca vitulina | Sea Mammal | 2022/6/24 | USA | North America |
| A/harbor s | Phoca vitulina | Sea Mammal | 2022/7/2 | USA | North America |
| A/harbor_s | Phoca vitulina | Sea Mammal | 2022/7/2 | USA | North America |
| A/harbor s | Phoca vitulina | Sea Mammal | 2022/7/2 | USA | North America |
| A/harbor s | Phoca vitulina | Sea Mammal | 2022/7/7 | USA | North America |
| A/harbor_s | Phoca vitulina | Sea Mammal | 2022/7/7 | USA | North America |
| A/Hawk/M | Hawk | Avian | 2022/4/27 | USA | North America |
| A/Hawk/N | Hawk | Avian | 2022/4/8 | USA | North America |
| A/Hawk/N | Hawk | Avian | 2022/4/9 | USA | North America |
| A/Hawk/N | Hawk | Avian | 2022/4/21 | USA | North America |
| A/Heron/M | Heron | Avian | 2022/4/24 | USA | North America |
| A/Herring_ | Herring gull | Avian | 2022/4/11 | USA | North America |
| A/herring | Larus argentatus | Avian | 2022/12/6 | USA | North America |
| A/Herring | Herring gull | Avian | 2022/2/25 | USA | North America |
| A/Herring_ | Herring gull | Avian | 2022/4/20 | USA | North America |
| A/Herring | Herring gull | Avian | 2022/4/20 | USA | North America |
| A/Herring | Herring gull | Avian | 2022/4/20 | USA | North America |
| A/Herring_ | Herring gull | Avian | 2022/4/20 | USA | North America |
| A/Hooded | Hooded merganser | Avian | 2022/3/1 | USA | North America |
| A/hooded_ | Lophodytes cucullatus | | 2022/3/1 | | |
| A/Hooded_ | Hooded merganser | Avian | 2022/4/14 | USA | North America |
| A/Hooded | Hooded merganser | Avian | 2022/4/12 | USA | North America |
| A/Hooded | Hooded merganser | Avian | 2022/2/28 | USA | North America |
| A/Hooded_ | Hooded merganser | Avian | 2022/4/4 | USA | North America |
| A/Hooded | Hooded merganser | Avian | 2022/4/4 | USA | North America |
| A/Hooded_ | Hooded merganser | Avian | 2022/4/4 | USA | North America |
| A/Hooded_ | Hooded merganser | Avian | 2022/4/3 | USA | North America |
| A/Humbold | Spheniscus humboldti | | 2023/2/28 | Chile | South America |
| A/Kelp_gu | Larus dominicanus | Avian | 2023/3/1 | Chile | South America |
| A/Lesser S | Lesser scaup | Avian | 2022/2/8 | USA | North America |
| A/Lesser S | Lesser scaup | Avian | 2022/2/16 | USA | North America |
| A/Lesser S | Lesser scaup | Avian | 2022/2/18 | USA | North America |
| A/Lesser_S | Lesser scaup | Avian | 2022/2/13 | USA | North America |
| A/Lesser S | Lesser scaup | Avian | 2022/2/24 | USA | North America |
| A/Lesser_S | Lesser scaup | Avian | 2022/3/18 | USA | North America |
| A/Lesser_S | Lesser scaup | Avian | 2022/3/23 | USA | North America |
| A/Lesser S | Lesser scaup | Avian | 2022/3/23 | USA | North America |
| A/Lesser_S | Lesser scaup | Avian | 2022/4/4 | USA | North America |
| A/lesser sc | Aythya affinis | Avian | 2022/2/12 | USA | North America |
| A/Lesser S | Lesser scaup | Avian | 2022/2/24 | USA | North America |
| A/Lesser_S | Lesser scaup | Avian | 2022/2/24 | USA | North America |
| A/Lesser_S | Lesser scaup | Avian | 2022/2/24 | USA | North America |
| A/Lesser S | Lesser scaup | Avian | 2022/2/24 | USA | North America |
| A/Lesser_S | Lesser scaup | Avian | 2022/2/24 | USA | North America |
| A/lesser_sc | Aythya affinis | Avian | 2022/2/12 | USA | North America |
| A/lesser sc | Aythya affinis | Avian | 2022/2/12 | USA | North America |
| A/lesser_sc | Aythya affinis | Avian | 2022/2/12 | USA | North America |
| A/lesser sc | Aythya affinis | Avian | 2022/2/12 | USA | North America |
| A/lesser sc | Aythya affinis | Avian | 2022/2/12 | USA | North America |
| A/lesser_sc | Aythya affinis | Avian | 2022/2/12 | USA | North America |
| A/Lesser_S | Lesser scaup | Avian | 2022/1/23 | USA | North America |
| A/Lesser s | Aythya affinis | Avian | 2022/1/23 | USA | North America |
| A/Lesser S | Lesser scaup | Avian | 2022/3/18 | USA | North America |
| A/Lesser_S | Lesser snow goose blue-morph | | 2022/3/14 | USA | North America |
| A/Lesser S | Lesser snow goose blue-morph | | 2022/3/14 | USA | North America |

| | | | | | |
|---|---|---|---|---|---|
| A/lesser sr | Anser caerulescens | | 2022/4/8 | | |
| A/Lesser S | Lesser snow goose white-morph | | 2022/3/14 | USA | North America |
| A/Lesser S | Lesser snow goose white-morph | | 2022/3/14 | USA | North America |
| A/Lesser S | Lesser snow goose white-morph | | 2022/3/14 | USA | North America |
| A/Lesser S | Lesser snow goose white-morph | | 2022/3/14 | USA | North America |
| A/Lesser_S | Lesser snow goose white-morph | | 2022/3/14 | USA | North America |
| A/Lesser S | Lesser snow goose white-morph | | 2022/3/14 | USA | North America |
| A/Magpie/ | Magpie | Avian | Apr-22 | USA | North America |
| A/Magpie/ | Magpie | Avian | 2022/4/21 | USA | North America |
| A/Mallard/ | Mallard | Avian | 2022/3/22 | USA | North America |
| A/Mallard/ | Mallard | Avian | 2022/2/3 | USA | North America |
| A/Mallard/ | Mallard | Avian | 2022/2/3 | USA | North America |
| A/Mallard/ | Mallard | Avian | 2022/2/3 | USA | North America |
| A/Mallard/ | Mallard | Avian | 2022/2/3 | USA | North America |
| A/Mallard/ | Mallard | Avian | 2022/2/3 | USA | North America |
| A/Mallard | Anas platyrhynchos | Avian | 2019/8/22 | Canada | North America |
| A/Mallard | Anas platyrhynchos | Avian | 2019/8/22 | Canada | North America |
| A/Mallard_ | Anas platyrhynchos | Avian | 2019/8/22 | Canada | North America |
| A/mallard | Anas platyrhynchos | Avian | Sep-22 | USA | North America |
| A/mallard | Anas platyrhynchos | Avian | 2022/9/14 | USA | North America |
| A/Mallard/ | Mallard | Avian | 2022/3/2 | USA | North America |
| A/Mallard/ | Mallard | Avian | 2022/4/18 | USA | North America |
| A/Mallard/ | Mallard | Avian | 2022/3/9 | USA | North America |
| A/Mallard/ | Mallard | Avian | 2022/2/7 | USA | North America |
| A/Mallard/ | Mallard | Avian | 2022/3/22 | USA | North America |
| A/Mallard/ | Mallard | Avian | 2022/3/25 | USA | North America |
| A/Mallard/ | Mallard | Avian | 2022/3/25 | USA | North America |
| A/Mallard/ | Mallard | Avian | 2022/3/11 | USA | North America |
| A/Mallard/ | Mallard | Avian | 2022/1/26 | USA | North America |
| A/Mallard/ | Mallard | Avian | 2022/1/26 | USA | North America |
| A/Mallard/ | Mallard | Avian | 2022/1/26 | USA | North America |
| A/Mallard/ | Mallard | Avian | 2022/1/26 | USA | North America |
| A/Mallard/ | Mallard | Avian | 2022/1/26 | USA | North America |
| A/Mallard/ | Mallard | Avian | 2022/1/26 | USA | North America |
| A/Mallard/ | Mallard | Avian | 2022/1/26 | USA | North America |
| A/Mallard/ | Mallard | Avian | 2022/1/26 | USA | North America |
| A/Mallard/ | Mallard | Avian | 2022/1/26 | USA | North America |
| A/Mallard/ | Mallard | Avian | 2022/1/26 | USA | North America |
| A/Mallard/ | Mallard | Avian | 2022/2/7 | USA | North America |
| A/Mallard/ | Mallard | Avian | 2022/2/7 | USA | North America |
| A/Mallard/ | Mallard | Avian | 2022/2/7 | USA | North America |
| A/Mallard/ | Mallard | Avian | 2022/2/7 | USA | North America |
| A/Mallard/ | Mallard | Avian | 2022/1/8 | USA | North America |
| A/Mallard/ | Mallard | Avian | 2022/1/29 | USA | North America |
| A/Mallard/ | Mallard | Avian | 2022/2/1 | USA | North America |
| A/Mallard/ | Mallard | Avian | 2022/2/1 | USA | North America |
| A/Mallard/ | Mallard | Avian | 2022/2/1 | USA | North America |
| A/Mallard/ | Mallard | Avian | 2022/2/1 | USA | North America |
| A/mallard/l | Anas platyrhynchos | Avian | 2022/12/1 | USA | North America |
| A/Mallard/ | Mallard | Avian | 2022/3/8 | USA | North America |
| A/Mallard/ | Mallard | Avian | 2022/3/8 | USA | North America |
| A/Mallard/ | Mallard | Avian | 2022/3/8 | USA | North America |
| A/Mallard/ | Mallard | Avian | 2022/3/9 | USA | North America |
| A/Mallard/ | Mallard | Avian | 2022/3/9 | USA | North America |
| A/Mallard/ | Mallard | Avian | 2022/2/12 | USA | North America |
| A/Mergans | Merganser | | 2022/3/23 | USA | North America |
| A/Mergans | Merganser | | 2022/4/11 | USA | North America |
| A/Mergans | Merganser | | 2022/3/31 | USA | North America |

| | | | | | |
|---|---|---|---|---|---|
| A/Mergans | Merganser | | 2022/4/18 | USA | North America |
| A/Mergans | Merganser | | 2022/4/26 | USA | North America |
| A/Mergans | Merganser | | 2022/3/2 | USA | North America |
| A/Mergans | Merganser | | 2022/3/21 | USA | North America |
| A/Mergans | Merganser | | 2022/4/5 | USA | North America |
| A/Mergans | Merganser | | 2022/3/22 | USA | North America |
| A/Mergans | Merganser | | 2022/3/22 | USA | North America |
| A/Mergans | Merganser | | 2022/3/22 | USA | North America |
| A/Mergans | Merganser | | 2022/3/22 | USA | North America |
| A/Mergus/l | Mergus | | 2022/4/5 | USA | North America |
| A/Mergus/l | Mergus | | 2022/4/7 | USA | North America |
| A/Mergus/l | Mergus | | 2022/4/6 | USA | North America |
| A/Muscovy | Muscovy duck | Avian | 2022/4/4 | USA | North America |
| A/Muscovy | Muscovy duck | Avian | 2022/4/4 | USA | North America |
| A/Muscovy | Muscovy duck | Avian | 2022/4/5 | USA | North America |
| A/Muscovy | Muscovy duck | Avian | 2022/4/5 | USA | North America |
| A/Muscovy | Muscovy duck | Avian | 2022/4/13 | USA | North America |
| A/Muscovy | Muscovy duck | Avian | 2022/4/13 | USA | North America |
| A/Muscovy | Muscovy duck | Avian | 2022/4/13 | USA | North America |
| A/Muscovy | Muscovy duck | Avian | 2022/4/22 | USA | North America |
| A/muscovy | Cairina moschata | | 2022/4/18 | | |
| A/Mute_Sv | Mute swan | Avian | 2022/4/11 | USA | North America |
| A/Mute_Sv | Mute swan | Avian | 2022/3/30 | USA | North America |
| A/Nannopt | Phalacrocorax brasilianus | | 2022/11/22 | Peru | South America |
| A/Nannopt | Phalacrocorax brasilianus | | 2022/11/22 | Peru | South America |
| A/Northern | Northern pintail | Avian | 2022/1/8 | USA | North America |
| A/Northern | Northern shoveler | Avian | 2022/1/31 | USA | North America |
| A/Northern | Northern shoveler | Avian | 2022/1/8 | USA | North America |
| A/Northern | Northern shoveler | Avian | 2022/1/29 | USA | North America |
| A/Owl/Tex | Owl | Avian | 2022/4/26 | USA | North America |
| A/Owl/Wy | Owl | Avian | 2022/4/26 | USA | North America |
| A/Panthera | Panthera leo | Nonhuman M | 2023/2/8 | Peru | South America |
| A/Parrot/M | Parrot | Avian | 2022/4/4 | USA | North America |
| A/Pekin_dı | Anas platyrhynchos | Avian | 2022/11/14 | USA | North America |
| A/Pelecanu | Pelecanus | | Dec-22 | Peru | South America |
| A/Pelecanu | Pelecanus thagus | | 2022/11/10 | Peru | South America |
| A/Pelecanu | Pelecanus thagus | | 2022/11/16 | Peru | South America |
| A/Pelecanu | Pelecanus thagus | | 2022/11/22 | Peru | South America |
| A/Pelican/ | Pelecanus sp. | | 2022/12/16 | Chile | South America |
| A/Pelican/ | Pelecanus sp. | | 2022/12/15 | Chile | South America |
| A/Pelican/ | Pelecanus sp. | | 2022/12/15 | Chile | South America |
| A/Pelican/ | Pelecanus sp. | | 2022/12/16 | Chile | South America |
| A/Pelican/ | Pelecanus sp. | | 2022/12/22 | Chile | South America |
| A/Pelican/ | Pelecanus sp. | | 2022/12/26 | Chile | South America |
| A/Pelican/ | Pelecanus sp. | | 2022/12/30 | Chile | South America |
| A/Pelican/( | Pelecanus thagus | | 2022/12/5 | Chile | South America |
| A/Pelican/( | Pelecanus thagus | | 2022/12/5 | Chile | South America |
| A/Pelican/( | Pelecanus thagus | | 2022/12/6 | Chile | South America |
| A/Pelican/( | Pelecanus thagus | | 2022/12/6 | Chile | South America |
| A/Pelican/( | Pelecanus thagus | | 2022/12/6 | Chile | South America |
| A/Pelican/( | Pelecanus thagus | | 2022/12/6 | Chile | South America |
| A/Pelican/( | Pelecanus thagus | | 2022/12/7 | Chile | South America |
| A/Pelican/l | Pelican | Avian | Mar-22 | USA | North America |
| A/Pelican/l | Pelican | Avian | Mar-22 | USA | North America |
| A/Pelican/l | Pelican | Avian | 2022/4/4 | USA | North America |
| A/Pelican/l | Pelecanus sp. | | 2023/1/26 | Chile | South America |
| A/Pelican/( | Pelecanus sp. | | 2023/1/24 | Chile | South America |
| A/Pelican/( | Pelecanus sp. | | 2023/1/24 | Chile | South America |

| | | | | | |
|---|---|---|---|---|---|
| A/pelican/I | Pelican | Avian | 2022/11/23 | Peru | South America |
| A/pelican/I | Pelican | Avian | 2022/11/24 | Peru | South America |
| A/pelican/I | Pelican | Avian | 2022/11/24 | Peru | South America |
| A/pelican/I | Pelican | Avian | 2022/11/24 | Peru | South America |
| A/Pelican/T | Pelecanus sp. | | 2022/12/9 | Chile | South America |
| A/Pelican/\ | Pelecanus sp. | | 2023/1/20 | Chile | South America |
| A/Pelican/\ | Pelecanus sp. | | 2023/1/20 | Chile | South America |
| A/Pelican/\ | Pelecanus sp. | | 2023/1/20 | Chile | South America |
| A/Pelican/\ | Pelecanus sp. | | 2023/1/23 | Chile | South America |
| A/Pelican/\ | Pelecanus sp. | | 2023/1/25 | Chile | South America |
| A/Peregrin | Peregrine falcon | Avian | 2022/4/15 | USA | North America |
| A/Peregrin | Peregrine falcon | Avian | 2022/3/11 | USA | North America |
| A/Peregrin | Peregrine falcon | Avian | 2022/4/19 | USA | North America |
| A/Peruvian | Peruvian_pelican | | Nov-22 | Peru | South America |
| A/Peruvian | Peruvian_pelican | | Nov-22 | Peru | South America |
| A/Pheasant | Pheasant | Avian | 2022/3/22 | USA | North America |
| A/Pheasant | Pheasant | Avian | 2022/4/12 | USA | North America |
| A/Pheasant | Pheasant | Avian | 2022/3/29 | USA | North America |
| A/Pheasant | Pheasant | Avian | 2022/3/29 | USA | North America |
| A/Pheasant | Pheasant | Avian | 2022/3/29 | USA | North America |
| A/poultry/I | Env | Environment | 2021/8/1 | Benin | Africa |
| A/poultry/I | Env | Environment | 2021/9/1 | Benin | Africa |
| A/poultry/I | Env | Environment | 2021/9/1 | Benin | Africa |
| A/poultry/I | Env | Environment | 2021/9/1 | Benin | Africa |
| A/Poultry/I | Poultry | Avian | 2022/3/9 | USA | North America |
| A/Redhead | Redhead | Avian | 2022/3/23 | USA | North America |
| A/Redhead | Redhead | Avian | 2022/3/23 | USA | North America |
| A/Redhead | Redhead | Avian | 2022/3/23 | USA | North America |
| A/Redhead | Redhead | Avian | 2022/3/23 | USA | North America |
| A/Redhead | Redhead | Avian | 2022/3/23 | USA | North America |
| A/Redhead | Redhead | Avian | Mar-22 | USA | North America |
| A/Red-shou | Red-shouldered hawk | | 2022/2/10 | USA | North America |
| A/Red-shou | Red-shouldered hawk | | 2022/4/18 | USA | North America |
| A/Red-shou | Red-shouldered hawk | | 2022/4/22 | USA | North America |
| A/Red-shou | Red-shouldered hawk | | 2022/4/22 | USA | North America |
| A/Red-shou | Red-shouldered hawk | | Feb-22 | USA | North America |
| A/red-shou | Buteo lineatus | | 2022/2/12 | USA | North America |
| A/Red-taile | Red-tailed hawk | Avian | 2022/4/20 | USA | North America |
| A/Red-taile | Red-tailed hawk | Avian | 2022/4/14 | USA | North America |
| A/Red-taile | Red-tailed hawk | Avian | 2022/4/14 | USA | North America |
| A/Red-taile | Red-tailed hawk | Avian | 2022/4/13 | USA | North America |
| A/Red-taile | Red-tailed hawk | Avian | 2022/4/27 | USA | North America |
| A/Red-taile | Red-tailed hawk | Avian | 2022/3/14 | USA | North America |
| A/red-tailed | Buteo jamaicensis | Avian | 2022/12/5 | USA | North America |
| A/red-tailed | Buteo jamaicensis | Avian | 2022/12/5 | USA | North America |
| A/red-tailed | Buteo jamaicensis | Avian | 2022/3/14 | USA | North America |
| A/red-tailed | Buteo jamaicensis | Avian | 2022/12/22 | USA | North America |
| A/Red-taile | Red-tailed hawk | Avian | 2022/3/6 | USA | North America |
| A/Red-taile | Red-tailed hawk | Avian | 2022/4/11 | USA | North America |
| A/Red-taile | Red-tailed hawk | Avian | 2022/4/22 | USA | North America |
| A/Red-taile | Red-tailed hawk | Avian | 2022/3/30 | USA | North America |
| A/Red-taile | Red-tailed hawk | Avian | 2022/3/30 | USA | North America |
| A/Red-taile | Red-tailed hawk | Avian | 2022/4/15 | USA | North America |
| A/Red-taile | Red-tailed hawk | Avian | 2022/4/15 | USA | North America |
| A/Red-taile | Red-tailed hawk | Avian | 2022/4/15 | USA | North America |
| A/Red-taile | Red-tailed hawk | Avian | 2022/4/14 | USA | North America |
| A/Red-taile | Red-tailed hawk | Avian | 2022/4/12 | USA | North America |
| A/Red-taile | Red-tailed hawk | Avian | 2022/4/13 | USA | North America |

| | | | | | |
|---|---|---|---|---|---|
| A/Red-taile | Red-tailed hawk | Avian | 2022/4/25 | USA | North America |
| A/Red-taile | Red-tailed hawk | Avian | 2022/4/25 | USA | North America |
| A/Red-taile | Red-tailed hawk | Avian | 2022/4/25 | USA | North America |
| A/Red-taile | Red-tailed hawk | Avian | 2022/4/22 | USA | North America |
| A/Red-taile | Red-tailed hawk | Avian | 2022/4/7 | USA | North America |
| A/Red-taile | Red-tailed hawk | Avian | 2022/3/29 | USA | North America |
| A/Red-taile | Red-tailed hawk | Avian | 2022/4/6 | USA | North America |
| A/Red-taile | Red-tailed hawk | Avian | 2022/4/1 | USA | North America |
| A/Red-taile | Red-tailed hawk | Avian | 2022/4/5 | USA | North America |
| A/Red-taile | Red-tailed hawk | Avian | 2022/4/6 | USA | North America |
| A/Red-taile | Red-tailed hawk | Avian | 2022/4/5 | USA | North America |
| A/Red-taile | Red-tailed hawk | Avian | 2022/4/8 | USA | North America |
| A/Red-taile | Red-tailed hawk | Avian | 2022/4/8 | USA | North America |
| A/Red-taile | Red-tailed hawk | Avian | 2022/4/10 | USA | North America |
| A/Red-taile | Red-tailed hawk | Avian | 2022/4/10 | USA | North America |
| A/Red-taile | Red-tailed hawk | Avian | 2022/4/25 | USA | North America |
| A/Red-taile | Red-tailed hawk | Avian | 2022/4/21 | USA | North America |
| A/Red-taile | Red-tailed hawk | Avian | 2022/4/21 | USA | North America |
| A/Red-taile | Red-tailed hawk | Avian | 2022/4/21 | USA | North America |
| A/Red-taile | Red-tailed hawk | Avian | 2022/4/21 | USA | North America |
| A/Red-taile | Red-tailed hawk | Avian | 2022/4/21 | USA | North America |
| A/Red-taile | Red-tailed hawk | Avian | 2022/4/21 | USA | North America |
| A/Red-taile | Red-tailed hawk | Avian | 2022/3/23 | USA | North America |
| A/Red-taile | Red-tailed hawk | Avian | 2022/3/23 | USA | North America |
| A/Red-taile | Red-tailed hawk | Avian | 2022/4/10 | USA | North America |
| A/Red-taile | Red-tailed hawk | Avian | 2022/4/9 | USA | North America |
| A/Red-taile | Red-tailed hawk | Avian | 2022/4/22 | USA | North America |
| A/Ring-bill | Ring-billed gull | Avian | 2022/3/7 | USA | North America |
| A/ring-bill | Larus delawarensis | | 2022/3/7 | | |
| A/Ring-nec | Ring-necked duck | Avian | 2022/3/23 | USA | North America |
| A/Ring-nec | Ring-necked duck | Avian | 2022/3/23 | USA | North America |
| A/Ring-nec | Ring-necked duck | Avian | 2022/3/23 | USA | North America |
| A/Ring-nec | Ring-necked duck | Avian | 2022/3/11 | USA | North America |
| A/Ring-nec | Ring-necked pheasant | Avian | 2022/4/26 | USA | North America |
| A/Rosss G | Ross's goose | Avian | 2022/3/18 | USA | North America |
| A/Rosss_G | Ross's goose | Avian | 2022/3/6 | USA | North America |
| A/Rosss G | Ross's goose | Avian | 2022/3/8 | USA | North America |
| A/Rosss_G | Ross's goose | Avian | 2022/3/8 | USA | North America |
| A/Rosss_G | Ross's goose | Avian | 2022/3/15 | USA | North America |
| A/Rosss G | Ross's goose | Avian | 2022/3/15 | USA | North America |
| A/Rosss_G | Ross's goose | Avian | 2022/3/12 | USA | North America |
| A/Rosss g | Anser rossii | | 2022/12/5 | USA | North America |
| A/Rosss G | Ross's goose | Avian | 2022/4/5 | USA | North America |
| A/Rosss_G | Ross's goose | Avian | 2022/4/6 | USA | North America |
| A/Rosss_G | Ross's goose | Avian | 2022/4/14 | USA | North America |
| A/Rosss G | Ross's goose | Avian | 2022/2/22 | USA | North America |
| A/Rosss_G | Ross's goose | Avian | Mar-22 | USA | North America |
| A/Rosss_G | Ross's goose | Avian | Mar-22 | USA | North America |
| A/Rosss G | Ross's goose | Avian | 2022/3/4 | USA | North America |
| A/Rosss_G | Ross's goose | Avian | 2022/3/23 | USA | North America |
| A/Rosss G | Ross's goose | Avian | 2022/3/23 | USA | North America |
| A/Rosss G | Ross's goose | Avian | 2022/3/23 | USA | North America |
| A/Rosss_G | Ross's goose | Avian | 2022/3/23 | USA | North America |
| A/Rosss_G | Ross's goose | Avian | 2022/3/23 | USA | North America |
| A/Rosss G | Ross's goose | Avian | 2022/3/22 | USA | North America |
| A/Rosss G | Ross's goose | Avian | 2022/3/25 | USA | North America |
| A/Rosss_G | Ross's goose | Avian | 2022/3/25 | USA | North America |
| A/Rosss G | Ross's goose | Avian | 2022/3/25 | USA | North America |

| | | | | | |
|---|---|---|---|---|---|
| A/Rosss G | Ross's goose | Avian | 2022/3/25 | USA | North America |
| A/Rosss G | Ross's goose | Avian | 2022/4/1 | USA | North America |
| A/Rosss G | Ross's goose | Avian | 2022/4/1 | USA | North America |
| A/Rosss G | Ross's goose | Avian | 2022/4/1 | USA | North America |
| A/Rosss G | Ross's goose | Avian | 2022/4/1 | USA | North America |
| A/Rosss_G | Ross's goose | Avian | 2022/4/1 | USA | North America |
| A/Rosss G | Ross's goose | Avian | 2022/4/1 | USA | North America |
| A/Rosss G | Ross's goose | Avian | 2022/4/1 | USA | North America |
| A/Rosss_G | Ross's goose | Avian | 2022/4/1 | USA | North America |
| A/Rosss G | Ross's goose | Avian | 2022/4/1 | USA | North America |
| A/Rosss G | Ross's goose | Avian | 2022/4/5 | USA | North America |
| A/Rosss G | Ross's goose | Avian | 2022/4/5 | USA | North America |
| A/Rosss G | Ross's goose | Avian | 2022/4/4 | USA | North America |
| A/Rosss G | Ross's goose | Avian | 2022/4/4 | USA | North America |
| A/Rosss_G | Ross's goose | Avian | 2022/4/4 | USA | North America |
| A/Rosss G | Ross's goose | Avian | 2022/4/5 | USA | North America |
| A/Rosss G | Ross's goose | Avian | 2022/4/6 | USA | North America |
| A/Rosss_G | Ross's goose | Avian | 2022/4/6 | USA | North America |
| A/Rosss G | Ross's goose | Avian | 2022/4/7 | USA | North America |
| A/Rosss G | Ross's goose | Avian | 2022/4/7 | USA | North America |
| A/Rosss_G | Ross's goose | Avian | 2022/4/10 | USA | North America |
| A/Rosss G | Ross's goose | Avian | 2022/4/10 | USA | North America |
| A/Rosss_g | Anser rossii | | 2022/4/8 | | |
| A/Rosss_G | Ross's goose | Avian | 2022/3/16 | USA | North America |
| A/Rosss G | Ross's goose | Avian | 2022/3/16 | USA | North America |
| A/Rosss G | Ross's goose | Avian | 2022/3/16 | USA | North America |
| A/Royal_T | Royal tern | | 2022/2/18 | USA | North America |
| A/Royal T | Royal tern | | 2022/2/18 | USA | North America |
| A/Royal_T | Royal tern | | 2022/3/16 | USA | North America |
| A/Royal_T | Royal tern | | 2022/3/21 | USA | North America |
| A/Royal T | Royal tern | | 2022/3/17 | USA | North America |
| A/Royal_T | Royal tern | | 2022/3/17 | USA | North America |
| A/royal ter | Thalasseus maximus | | 2022/3/12 | | |
| A/royal ter | Thalasseus maximus | | 2022/3/17 | | |
| A/Sanderli | Calidris alba | Avian | 2022/12/30 | Chile | South America |
| A/Sanderli | Calidris alba | Avian | 2023/3/3 | Chile | South America |
| A/Sanderli | Sanderling | Avian | 2022/3/1 | USA | North America |
| A/Sanderli | Sanderling | Avian | 2022/3/2 | USA | North America |
| A/Sanderli | Sanderling | Avian | 2022/3/2 | USA | North America |
| A/Sanderli | Sanderling | Avian | 2022/3/3 | USA | North America |
| A/Sanderli | Sanderling | Avian | 2022/3/4 | USA | North America |
| A/Sanderli | Sanderling | Avian | 2022/3/6 | USA | North America |
| A/Sanderli | Sanderling | Avian | 2022/2/21 | USA | North America |
| A/Sanderli | Sanderling | Avian | 2022/4/11 | USA | North America |
| A/Sanderli | Sanderling | Avian | 2022/11/22 | Peru | South America |
| A/Sandhill | Sandhill crane | | 2022/4/21 | USA | North America |
| A/Sharp-sh | Sharp-shinned hawk | | 2022/3/11 | USA | North America |
| A/Silver_P | Silver pheasant | Avian | 2022/8/31 | USA | North America |
| A/Snow G | Snow goose | Avian | 2022/3/18 | USA | North America |
| A/Snow_G | Snow goose | Avian | 2022/3/18 | USA | North America |
| A/Snow G | Snow goose | Avian | 2022/3/18 | USA | North America |
| A/Snow G | Snow goose | Avian | 2022/3/18 | USA | North America |
| A/Snow_G | Snow goose | Avian | 2022/3/18 | USA | North America |
| A/Snow_G | Snow goose | Avian | 2022/3/23 | USA | North America |
| A/Snow G | Snow goose | Avian | 2022/3/8 | USA | North America |
| A/Snow G | Snow goose | Avian | 2022/3/15 | USA | North America |
| A/Snow_G | Snow goose | Avian | 2022/3/22 | USA | North America |
| A/Snow G | Snow goose | Avian | 2022/3/22 | USA | North America |

| | | | | | |
|---|---|---|---|---|---|
| A/Snow G | Snow goose | Avian | 2022/3/22 | USA | North America |
| A/Snow G | Snow goose | Avian | 2022/2/28 | USA | North America |
| A/Snow G | Snow goose | Avian | 2022/2/28 | USA | North America |
| A/Snow G | Snow goose | Avian | 2022/3/6 | USA | North America |
| A/Snow G | Snow goose | Avian | 2022/3/4 | USA | North America |
| A/Snow_G | Snow goose | Avian | 2022/3/4 | USA | North America |
| A/Snow G | Snow goose | Avian | 2022/3/12 | USA | North America |
| A/Snow G | Snow goose | Avian | 2022/3/12 | USA | North America |
| A/Snow_G | Snow goose | Avian | 2022/3/12 | USA | North America |
| A/Snow G | Snow goose | Avian | 2022/3/12 | USA | North America |
| A/Snow G | Snow goose | Avian | 2022/3/25 | USA | North America |
| A/snow go | Anser caerulescens | Avian | 2022/12/5 | USA | North America |
| A/snow go | Anser caerulescens | | 2022/3/6 | | |
| A/snow go | Anser caerulescens | | 2022/3/4 | | |
| A/snow_go | Anser caerulescens | Avian | 2022/3/14 | USA | North America |
| A/snow go | Anser caerulescens | Avian | 2022/3/12 | USA | North America |
| A/snow go | Anser caerulescens | Avian | 2022/3/12 | USA | North America |
| A/snow_go | Anser caerulescens | Avian | 2022/3/12 | USA | North America |
| A/snow go | Anser caerulescens | Avian | 2022/3/12 | USA | North America |
| A/snow go | Anser caerulescens | Avian | 2022/3/12 | USA | North America |
| A/snow_go | Anser caerulescens | | 2022/3/25 | | |
| A/Snow G | Snow goose | Avian | 2022/2/8 | USA | North America |
| A/Snow_G | Snow goose | Avian | 2022/2/8 | USA | North America |
| A/Snow_G | Snow goose | Avian | 2022/2/18 | USA | North America |
| A/snow go | Anser caerulescens | Avian | 2022/2/8 | USA | North America |
| A/snow go | Anser caerulescens | Avian | 2022/12/5 | USA | North America |
| A/snow_go | Anser caerulescens | Avian | 2022/12/5 | USA | North America |
| A/Snow G | Snow goose | Avian | 2022/3/28 | USA | North America |
| A/Snow_G | Snow goose | Avian | 2022/4/7 | USA | North America |
| A/Snow_G | Snow goose | Avian | 2022/2/22 | USA | North America |
| A/Snow G | Snow goose | Avian | 2022/2/22 | USA | North America |
| A/Snow_G | Snow goose | Avian | 2022/2/22 | USA | North America |
| A/Snow G | Snow goose | Avian | 2022/2/22 | USA | North America |
| A/Snow G | Snow goose | Avian | 2022/2/22 | USA | North America |
| A/Snow G | Snow goose | Avian | 2022/2/23 | USA | North America |
| A/Snow_G | Snow goose | Avian | 2022/3/4 | USA | North America |
| A/Snow G | Snow goose | Avian | 2022/3/4 | USA | North America |
| A/Snow_G | Snow goose | Avian | 2022/3/4 | USA | North America |
| A/Snow_G | Snow goose | Avian | 2022/3/8 | USA | North America |
| A/Snow G | Snow goose | Avian | 2022/3/8 | USA | North America |
| A/Snow_G | Snow goose | Avian | 2022/3/8 | USA | North America |
| A/Snow G | Snow goose | Avian | 2022/3/8 | USA | North America |
| A/Snow G | Snow goose | Avian | 2022/3/12 | USA | North America |
| A/Snow_G | Snow goose | Avian | 2022/3/12 | USA | North America |
| A/Snow_G | Snow goose | Avian | 2022/3/22 | USA | North America |
| A/Snow G | Snow goose | Avian | 2022/4/8 | USA | North America |
| A/Snow_G | Snow goose | Avian | 2022/4/9 | USA | North America |
| A/Snow_G | Snow goose | Avian | 2022/3/23 | USA | North America |
| A/Snow G | Snow goose | Avian | 2022/2/24 | USA | North America |
| A/Snow_G | Snow goose | Avian | Mar-22 | USA | North America |
| A/Snow G | Snow goose | Avian | Mar-22 | USA | North America |
| A/Snow G | Snow goose | Avian | 2022/1/21 | USA | North America |
| A/Snow_G | Snow goose | Avian | 2022/3/21 | USA | North America |
| A/Snow_G | Snow goose | Avian | 2022/3/29 | USA | North America |
| A/Snow G | Snow goose | Avian | 2022/3/29 | USA | North America |
| A/Snow G | Snow goose | Avian | 2022/3/29 | USA | North America |
| A/Snow_G | Snow goose | Avian | 2022/3/21 | USA | North America |
| A/Snow G | Snow goose | Avian | 2022/3/23 | USA | North America |

| | | | | | |
|---|---|---|---|---|---|
| A/Snow_G | Snow goose | Avian | 2022/3/23 | USA | North America |
| A/Snow_G | Snow goose | Avian | 2022/3/23 | USA | North America |
| A/Snow_G | Snow goose | Avian | 2022/3/23 | USA | North America |
| A/Snow_G | Snow goose | Avian | 2022/3/23 | USA | North America |
| A/Snow_G | Snow goose | Avian | 2022/3/23 | USA | North America |
| A/Snow_G | Snow goose | Avian | 2022/3/23 | USA | North America |
| A/Snow_G | Snow goose | Avian | 2022/3/23 | USA | North America |
| A/Snow_G | Snow goose | Avian | 2022/3/23 | USA | North America |
| A/Snow_G | Snow goose | Avian | 2022/3/23 | USA | North America |
| A/Snow_G | Snow goose | Avian | 2022/3/23 | USA | North America |
| A/Snow_G | Snow goose | Avian | 2022/3/23 | USA | North America |
| A/Snow_G | Snow goose | Avian | 2022/3/23 | USA | North America |
| A/Snow_G | Snow goose | Avian | 2022/3/22 | USA | North America |
| A/Snow_G | Snow goose | Avian | 2022/3/22 | USA | North America |
| A/Snow_G | Snow goose | Avian | 2022/3/22 | USA | North America |
| A/Snow_G | Snow goose | Avian | 2022/3/22 | USA | North America |
| A/Snow_G | Snow goose | Avian | 2022/3/25 | USA | North America |
| A/Snow_G | Snow goose | Avian | 2022/3/25 | USA | North America |
| A/Snow_G | Snow goose | Avian | 2022/3/26 | USA | North America |
| A/Snow_G | Snow goose | Avian | 2022/3/26 | USA | North America |
| A/Snow_G | Snow goose | Avian | 2022/3/26 | USA | North America |
| A/Snow_G | Snow goose | Avian | 2022/4/1 | USA | North America |
| A/Snow_G | Snow goose | Avian | 2022/4/1 | USA | North America |
| A/snow_go | Anser caerulescens | | 2022/4/8 | | |
| A/snow_go | Anser caerulescens | | 2022/4/8 | | |
| A/snow_go | Anser caerulescens | | 2022/4/8 | | |
| A/South_A | Otaria byronia | Nonhuman M | 2023/3/1 | Chile | South America |
| A/south_an | South american sea lion | | 2023/2/7 | Peru | South America |
| A/South_A | South american sea lion | | 2023/1/23 | Peru | South America |
| A/South_A | Sterna hirundinacea | | 2023/2/23 | Chile | South America |
| A/striped_s | Mephitis mephitis | Nonhuman M | 2023/1/27 | USA | North America |
| A/striped_s | Mephitis mephitis | Nonhuman M | 2023/2/24 | USA | North America |
| A/Swan/OI | Swan | Avian | 2022/9/25 | USA | North America |
| A/Turkey/A | Meleagris gallopavo | Avian | 2023/3/14 | Chile | South America |
| A/Turkey/N | Turkey | Avian | 2022/4/15 | USA | North America |
| A/Turkey/N | Turkey | Avian | 2022/4/15 | USA | North America |
| A/Turkey/N | Meleagris gallopavo | Avian | 2023/3/7 | Chile | South America |
| A/Turkey/N | Meleagris gallopavo | Avian | 2023/3/10 | Chile | South America |
| A/Turkey/C | Turkey | Avian | 2022/9/12 | USA | North America |
| A/Turkey_\ | Cathartes aura | | 2022/12/17 | Chile | South America |
| A/Vulpes_\ | Vulpes vulpes | Nonhuman M | 2022/4/11 | USA | North America |
| A/Vulpes_\ | Vulpes vulpes | Nonhuman M | 2022/4/25 | USA | North America |
| A/Vulpes_\ | Vulpes vulpes | Nonhuman M | 2022/5/4 | USA | North America |
| A/Vulpes_\ | Vulpes vulpes | Nonhuman M | 2022/4/27 | USA | North America |
| A/Vulpes_\ | Vulpes vulpes | Nonhuman M | 2022/5/20 | USA | North America |
| A/wild_duc | Wild duck | Avian | 2022/10/9 | Colombia | South America |
| A/Wood_D | Wood duck | Avian | 2022/8/31 | USA | North America |
