## Supplementary figures and images for "Transcontinental Spread of HPAI H5N1 from South America to Antarctica via Avian Vectors"

### AppendixFigure1.tif

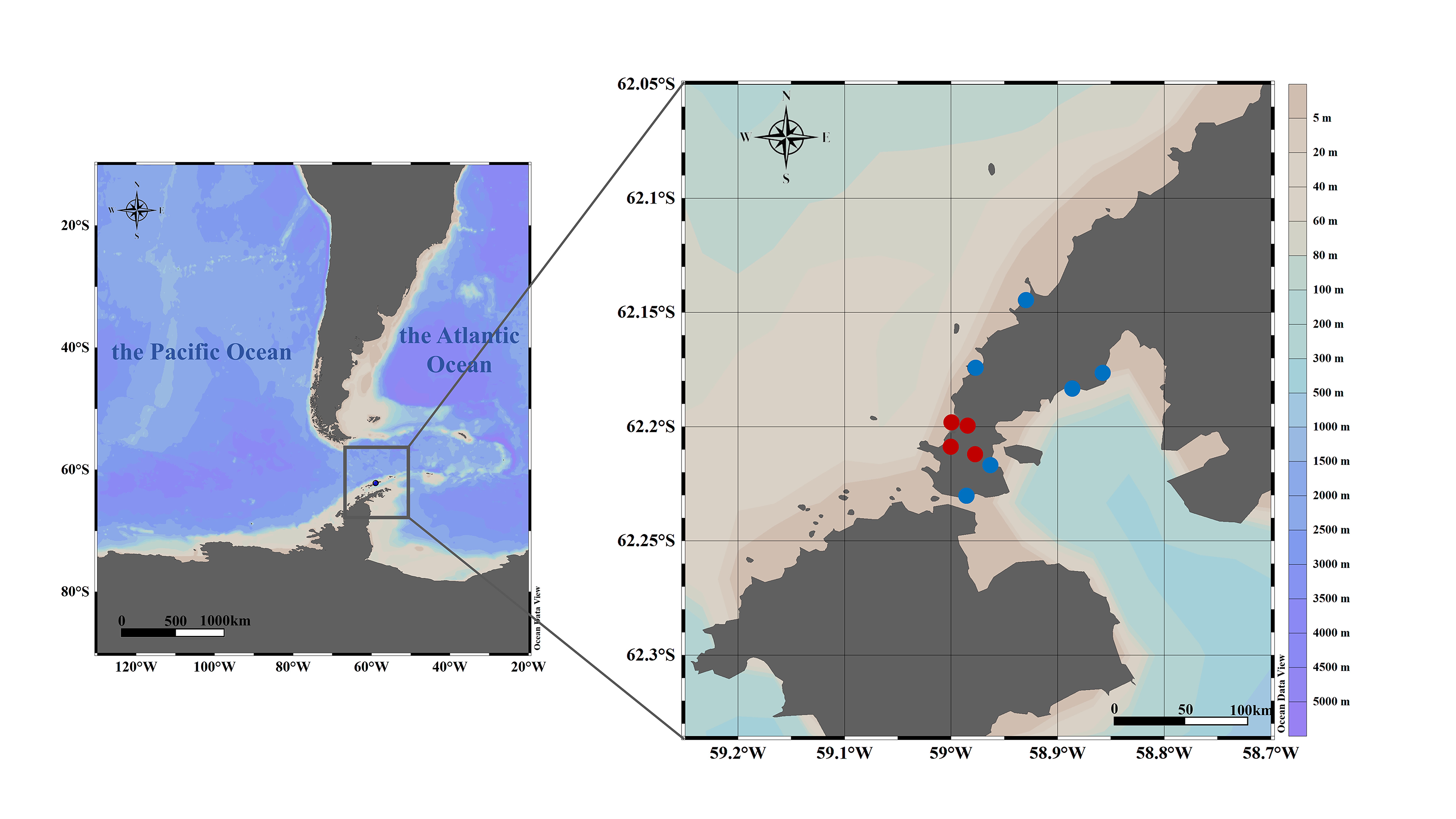

### AppendixFigure2.tif

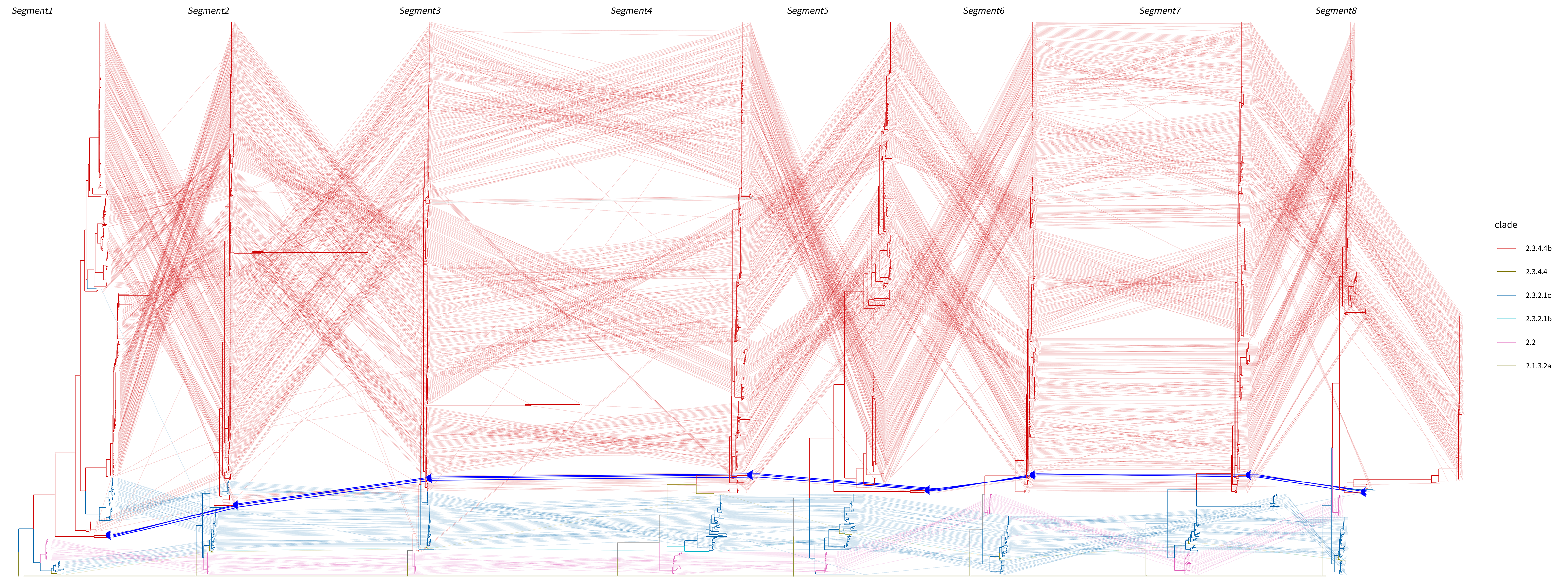

### AppendixFigure3 .tif

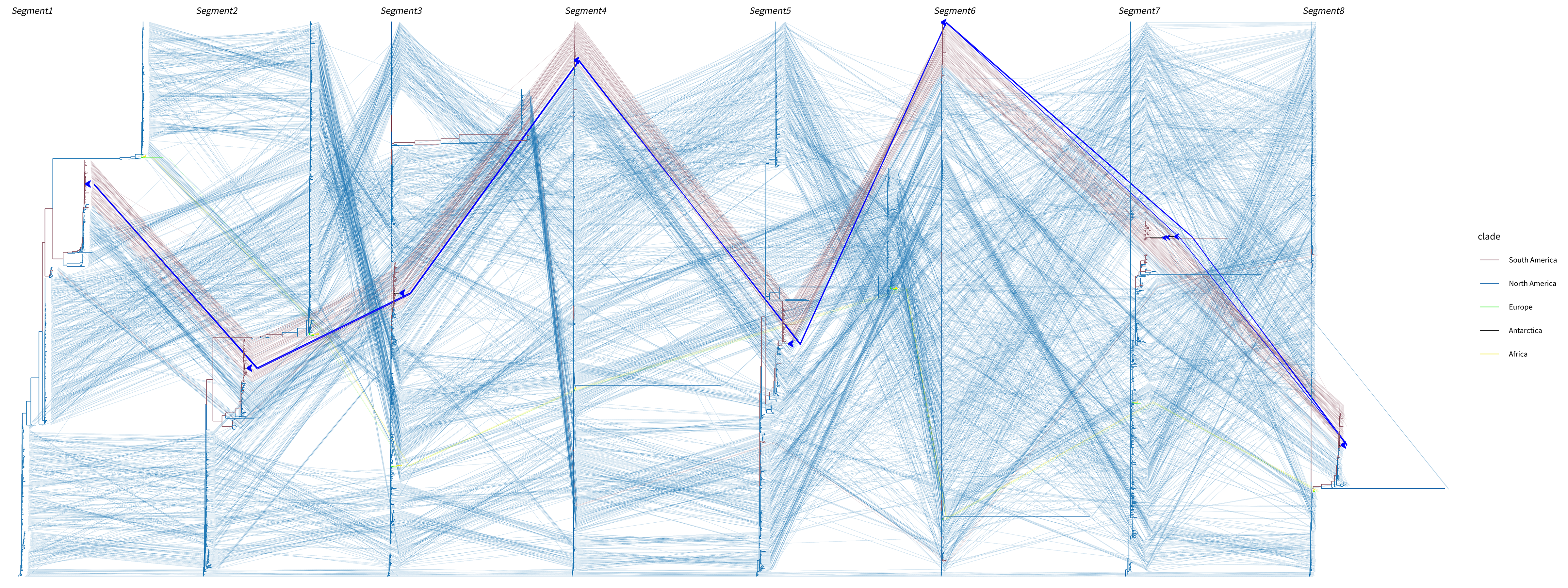

### AppendixFigure4.tif

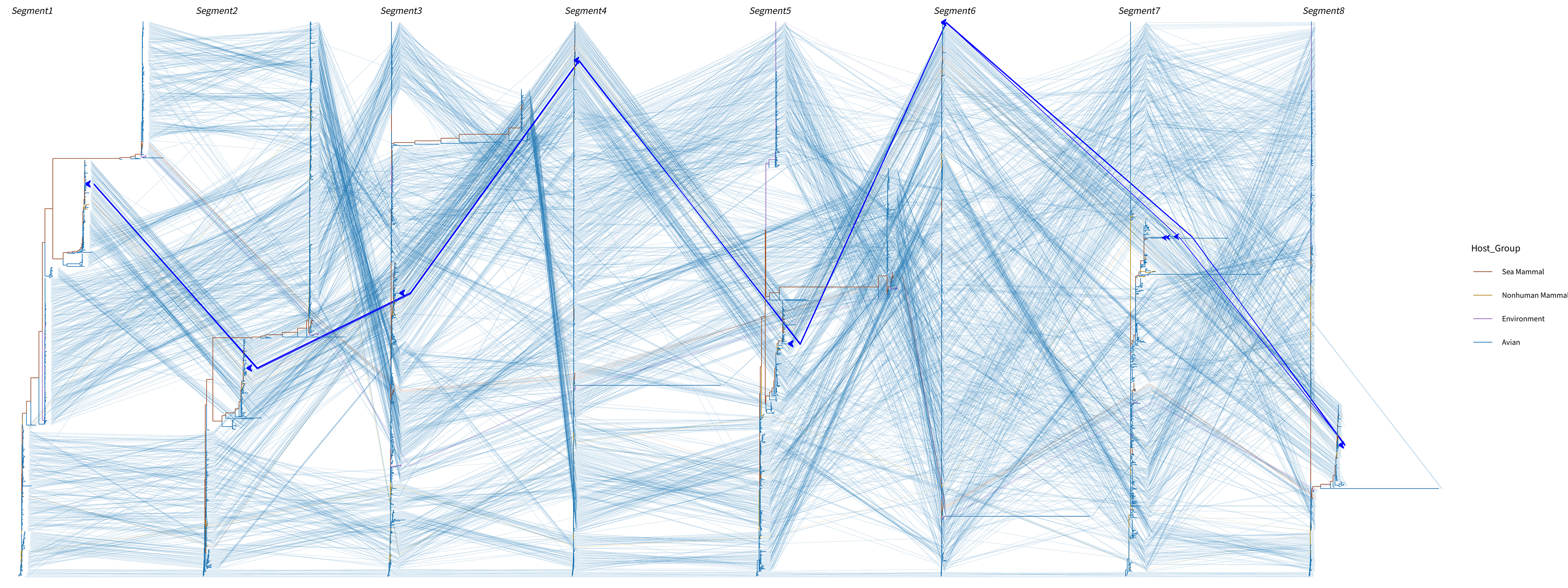

### Appendixfigure5.tif

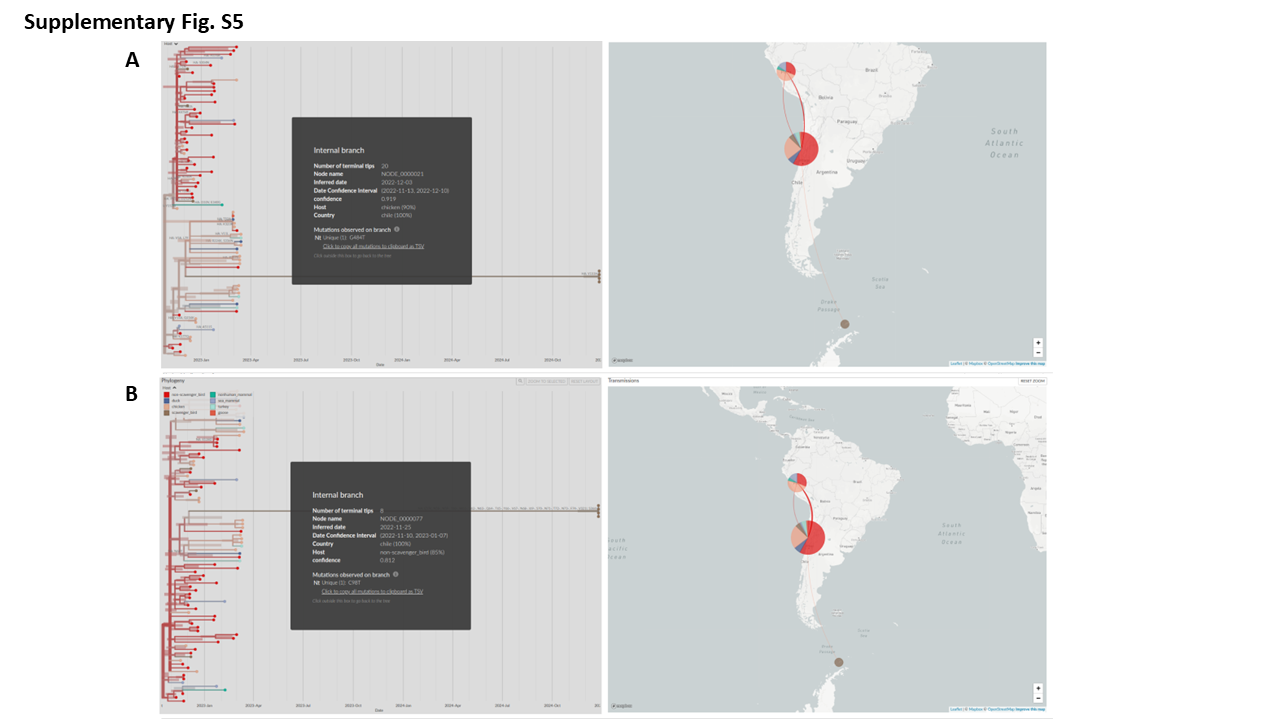
